# Genome-Wide Identification of Epigenetic Regulators in *Quercus suber*

**DOI:** 10.1101/2020.02.15.948950

**Authors:** HG Silva, RS Sobral, AP Magalhães, L Morais-Cecílio, MMR Costa

## Abstract

Modifications of DNA and histones, including methylation and acetylation, are critical for the epigenetic regulation of gene expression during plant development, particularly during environmental adaptation processes. However, information on the enzymes catalyzing all these modifications in perennial trees, such as *Quercus suber*, is still not available. In this study, several epigenetic modifier proteins, including eight DNA methyltransferases (DNA Mtases), three DNA demethylases (DDMEs) and ninety-one histone modifiers including thirty-five histone methyltransferases (HMTs), twenty-six histone demethylases (HDMTs), eight histone acetyltransferases (HATs) and twenty-two histone acetylases (HDACs) were identified in *Q. suber*. Phylogenetic analyses of the DNA and histone modifier proteins were performed using several plant species homologs, enabling the classification of the *Q. suber* proteins. Additional *in silico* analysis showed that some *Q. suber* DNA Mtases, DMEs and histone modifiers have the typical domains found in the plant model Arabidopsis, which might suggest a conserved functional role. A link between the expression levels of each gene in different *Q. suber* tissues (buds, flowers, acorns, embryos, cork and roots) with the functions already known for their closest homologs in other species was also established. Therefore, the data generated here are important for future studies exploring the role of epigenetic regulators in this economically important species.

## Introduction

Many essential biological processes are dependent on regulatory gene networks, due to the activation and/or silencing of key genes. This is achieved by the activity of particular transcription factors but also, by DNA and histone-modification enzymes that alter chromatin conformation (euchromatin or heterochromatin), in a process classically identified as epigenetic regulation. This control at the chromatin level is very dynamic and can be reversible, which means that inactive condensed regions of chromatin may be easily decondensed allowing for subsequent gene expression.

The best-known DNA modification that regulates gene expression is cytosine methylation that is catalyzed by several DNA methyltransferases. In plants, DNA methyltransferases methylate DNA in the carbon-5 of cytosine (5mC) in distinct sequence contexts such as CG and CHG, but also in the asymmetrical context CHH, in which H is any nucleotide but G [1]. While in plants 5mC in gene promoter regions is often associated with gene silencing, in coding regions, although yet controversial, 5mC has been correlated with active gene transcription [2,3]. Methylation events can be divided into two mechanisms: *de novo* and maintenance of the methylation status. *De novo* methylation implies the methylation of previously unmethylated cytosine residues without a “template” resulting in new methylation patterns. The maintenance of the methylation status is the process by which preexisting methylated residues serve as a template to the replication-coupled DNA methylation [4]. The methylation process is controlled by four families of DNA methyltransferases (Mtases), which are classified by their linear domain arrangement: Methyltransferase (MET), Chromomethylase (CMT), Domains Rearranged Methyltransferase (DRM) and DNA Methyltransferase homolog 2 (DNMT2). MET maintains CG methylation of heterochromatic regions enriched with transposable elements (TEs) and repeats, and in intragenic regions [5,6]. CMT and DRM mediate CHG (CMT3) and CHH (CMT2 and DRM2) methylation [7,8]. While the majority of the methyltransferases maintain the DNA methylation status, only DRM2 establishes *de novo* methylation in all three motif contexts [9]. *De novo* DNA methylation is triggered by the activation of RNA-directed DNA methylation (RdDM), in which small interfering RNAs (siRNAs) are generated, ending with a downstream methylation targeting phase that is mediated by DRM2 [8]. CMT family genes can initiate *de novo* DNA methylation at sites with specific histone modifications and target transposons, as well as heterochromatin during replication [10]. The role of DNMT2 family in plant DNA methylation remains unclear.

DNA methylation of 5mC can be reversed passively during DNA replication or actively through base excision repair mechanisms initiated by DNA glycosylases [11–14]. Functional studies of the DNA glycosylases REPRESSOR OF SILENCING 1 (ROS1) [11], and DEMETER (DME) [12,13] suggested roles in the inhibition of gene silencing by demethylating DNA. DEMETER-LIKE 2 and 3 (DML2 and 3) are also DNA demethylases (DDME) that, instead of reactivating gene expression, prevent the deleterious effect resulted from the accumulation of DNA methylation at or near some genes [14].

Epigenetic regulation is also accomplished by the acetylation and methylation of histones. Histone modifier enzymes add or remove chemical groups, mainly onto the lysine residues, in the amino-terminal tails of histone 3 (H3) and 4 (H4). The deposition of acetylation marks is mediated by histone acetyltransferases (HATs), a mark that can be removed by histone deacetylases (HDACs). Histone acetylation is frequently related with increased gene expression while deacetylation is associated to transcriptional repression [15]. HATs are grouped in four families: GNAT (Gcn5-related N-acetyltransferase); MYST (MOZ, Ybf2/Sas3, Sas2, and Tip60), CBP (cAMP-responsive element-binding protein-binding protein) and TAFII250 (TATA-binding protein-associated factor family) based on sequence homology and mode of action [16] while HDACs are divided in three families: RPD3/HDA1 (REDUCED POTASSIUM DEPENDENCY 3/ HISTONE DEACETYLASE 1), HD2 (HISTONE DEACETYLASE 2) and SIR2 (SILENT INFORMATION REGULATOR 2) based on sequence similarity and cofactor dependency [17]. Like acetylation, histone methylation is also a reversible process catalyzed by histone methyltransferases (HMTs) and histone demethylases (HDMTs). HMTs are mainly divided in five classes, based on the aminoacid conservation of the SET (Suppressor of variegation, Enhancer of zeste, and Trithorax in Drosophila) domain [18]. Each class of SET domain proteins has specificity for a particular histone residue. For example, while class I proteins transfer three methyl groups to lysine 27 of H3 (H3K27me3), which is correlated with gene silencing, class III proteins are responsible for the deposition of methyl groups associated with active gene expression like mono-, di- and tri-methylation at the lysine 4 of H3 (H3K4me1/2/3) site. Histone methylation marks are removed by two types of HDMTs, histone KDM1/LSD1 (lysine demethylase 1) and JmjC (Jumonji C) domain-containing proteins [19,20], depending on the cofactor required to act [21].

Plant development is controlled not only by endogenous but also by environmental cues, which requires a tight control of gene regulation. Adverse environmental factors, such as severe temperature, drought, saline and biotic stress can alter the longevity of many species, specially of long living species such as *Quercus suber*, able to live over a span of 200 years. *Q. suber* belongs to the Fagaceae family and is one of the most economically and ecologically important forest species in the Mediterranean basin, being the dominant tree of the Portugal oak woodlands - *Montados*. To provide for their unique attributes, both their longevity and annual developmental transitions, *Q. suber* trees need to have highly dynamic regulatory mechanisms in order to adapt and survive to a variable climate. So, the identification and characterization of chromatin modifier proteins is of great importance. In *Q. suber*, mutant lines are impossible to be created, due to its long life cycle and its recalcitrant behavior to transformation; however, the relevance of the genes codifying for these enzymes could be studied by gene expression analysis.

In this study, the genome-wide identification of DNA and histone modification enzymes in *Q. suber* was obtained by data mining available genomes and transcriptomes through blast analysis. The phylogeny and composition of each gene family was identified and a brief overview of the transcriptional regulation of these newly identified genes in *Q. suber* was obtained from different tissues (buds, flowers, acorns, embryos, roots and cork) using publicly available data. Thus, this work provides a valuable source of information on genes potentially involved in epigenetic regulation and should greatly facilitate further studies in *Q. suber*.

## Materials and Methods

### Identification of *Q. suber* DNA (De)Methyltransferases and Histone modifiers

The protein sequences of *Arabidopsis thaliana* DNA Mtases and DDME, as well as some histone modifiers (HATs, HDACs, HMTs and HDMTs) were retrieved from The Arabidopsis Information Resource (TAIR) database (http://www.arabidopsis.org/). These sequences were used as queries to search for *Q. suber* homologs using BLASTp. To ensure that all *Q. suber* DNA Mtases genes were identified, an additional search was performed in the CorkOakDB (http://www.corkoakdb.org/search) using the protein domains identification (InterPro ID) common to DNA Mtases (C-5 cytosine methyltransferase IPR001525), HATs (histone acetyltransferase domain, MYST-type IPR002717 and histone acetyltransferase Rtt109/CBP type IPR013178), HDACs (Histone deacetylase IPR000286 and Sirtuin IPR003000), HMTs(SET domain IPR001214) and HDMTs (JmjC domain IPR003347, Amine oxidase IPR002937 and SWIRM domain IPR007526). Proteins identified by these approaches were recorded and redundancy removed. Each corresponding *Q. suber* EST sequence and Arabidopsis protein were used to perform a search in the *Q. suber* genome (CorkOak1.0 version released on January 2018 under the RefSeq assembly: GCF_002906115.1) using the BLASTn and BLASTp algorithm, and the complete protein sequences were retrieved. All the epigenetic modifier enzymes identified in *Q. suber* are listed in table S1.

### Prediction of protein domains

*Q. suber* protein sequences were analyzed for recognizable domains using NCBI Batch Conserved Domain search tool (http://www.ncbi.nlm.nih.gov/Structure/bwrpsb/bwrpsb.cgi). The presence of specific domains and their organization was also verified using the Simple Modular Architecture Research Tool (SMART) (http://smart.embl-heidelberg.de). Schematic diagram of protein domain structures with its functional domains were constructed using Illustrator for Biological Sequences (IBS) (version IBS 1.0.1) (http://ibs.biocuckoo.org/index.php).

### Phylogenetic analysis

*A. thaliana* (At) protein sequences were used as queries to search for homologs in other species such as *Brassica rapa* (Br), *Glycine max* (Gm), *Jatropha curcas* (Jc), *Populus trichocarpa* (Pt), *Prunus persica* (Pp), *Prunus mume* (Pm), *Vitis vinifera* (Vv), *Cucumis melo* (Cm), *Cucumis sativus* (Cs) and *Juglans regia* (Jr) using the BLASTp tool in NCBI (https://blast.ncbi.nlm.nih.gov/). Protein homologs from other *Fagaceae* species (*Castanea mollissima* (Cmo) and *Quercus robur* (Qr)) were also included by performing a local BLASTp (prfectBLAST 2.0) against the Draft of Whole Genome Sequence (v1.1) of *C. molissima* (http://www.hardwoodgenomics.org/chinese-chestnut-genome), and a tBLASTn (prfectBLAST 2.0) search against *Quercus robur* data (Oak assemby V3_OCV3-91k). Protein data from *Oryza sativa* (Os) was retrieved from RGAP release 7 (http://rice.plantbiology.msu.edu/) to be used as an out-group. All the amino acid sequences were aligned using the alignment tool CLUSTAL W [22], using the Gonnet Protein Weight Matrix, and changing the default parameters of the Multiple Alignment Gap Opening penalty to 3 and the Multiple Alignment Gap Extension penalty to 1.8, more appropriate to proteins [23]. Phylogenetic trees were constructed using the Maximum-likelihood algorithm with the Jones-Taylor-Thornton (JTT) correction model (MEGA 7.0 software). The bootstrap consensus tree was inferred from 1000 replicates to obtain a support value for each branch.

### Transcriptome data analysis

RNA sequencing data was acquired from the Sequencing Read Archive of NCBI (https://www.ncbi.nlm.nih.gov/sra/) using the experiments provided by the high-throughput sequencing of the *Q. suber* transcriptome [24]. The *Q. suber* transcriptome was obtained using tissues at different stages/conditions of development such as: acorns from developmental stage 2 (ERR490202), 3 and 4 (ERR490203) and 5 (ERR490204) [25]; a pool of embryos collected from the acorns of stage 1 to 8 (ERR490207) [25]; cork of bad (SRR1009171) and good (SRR1009172) quality [26]; male (SRR1609152) and female (SRR1609153) flowers [27]; roots of plants with different degrees of watering: medium (SRR1812375), low (SRR1812376) and abundant (SRR1812377) [28]; red and opened buds (SRR5345606) and dormant and swollen buds (SRR5345607) [29]. The libraries where first trimmed to remove SMART adapters, present in the 454 libraries, using AlienTrimmer [30] and then filtered by QTRIM [31] using default quality parameters. Alignment against the *Q. suber* published genome CorkOak1.0 was performed with Burrows–Wheeler Aligner (BWA) using the BWA-MEM algorithm [32]. The alignment files were used to quantify gene expression using feature Counts [33].

### Differential expression analysis

Gene expression was analyzed in different *Q. suber* libraries using the read counts generated after the mapping of 454 reads into the CorkOak1.0 genome. Each expression pattern was normalized and estimated by DESeq2, a R package for differential expression analysis [34]. The Rlog function of DESeq2 was also used to transform the count data to the log2 scale in order to minimize variances between samples with fewer counts, taking into account each library size. The resulting values were Z scored, scaled and used for visualization and clustering. The R package NMF was used to plot heatmaps using the *aheatmap* function that is based on the Euclidian algorithm. Gene expression was also studied in the phylogenetically closed species *Q. robur* using transcriptomic data obtained by Lesur *et al*. (2015), in which the number of normalized read counts for root, eco-dormant bud, swelling bud, leaf, *in vitro* dedifferentiated *calli* and secondary differentiating xylem is available.

## Results

Epigenetic regulation plays an important role in the control of gene expression during plant growth and development. Although, significant advances in this field have been made in the model species *A. thaliana*, less has been reported in perennial tree species. In *Q. suber*, there is no knowledge about the genes that encode chromatin regulators. Therefore, in order to identify histone and DNA modifier enzymes with a potential role in epigenetic mark deposition, sequence-based searches and phylogenetic analysis were performed.

### Identification and classification of *Q. suber* DNA methyltransferases

Eight DNA Mtases proteins were identified based on a whole sequence similarity search and a C-5 cytosine methyltransferase (IPR001525) domain search. The DNA Mtases of each subfamily in *Q. suber* share the conserved catalytic domain with other plant species homologues and harbor their characteristic small motifs (Fig S1, Fig S2, Fig S3 and Fig S4) [36–38]. A single *Q. suber* gene was identified within the MET subfamily (QsMET1), as well as in the DNMT2 subfamily (QsDNMT2). QsMET1 contains the expected domains: replication foci domain RFD (PF12047), bromo adjacent homology (BAH) domain (PF01426) and DNA methyltransferase domain (PF00145) (Fig 1); a domain topology observed in the closest homologs of Arabidopsis, globe artichoke, strawberry, soybean, carrot, tomato, tobacco, pea, poplar, peach and rice [38–44]. QsDNMT2 contains a single DNA mtase domain, as previously identified in the *A. thaliana* counterpart (Fig 1) [38]. In the CMT subfamily, four homolog proteins were identified, two different putative CMT2 proteins (named in this work as QsCMT2.1 and QsCMT2.2), QsCMT1 and QsCMT3. Members of the CMT subfamily have the C-terminal DNA Mtase domain, a chromodomain (CHR) (PF00385), which has been proposed as critical to transport these proteins to heterochromatic regions [45] and, the BAH domain, crucial for DNA methylation maintenance during DNA replication [46] (Fig 1). Two *Q. suber* proteins were identified belonging to DRM subfamily, QsDRM2 and QsDRM3 (Fig 1). Both QsDRM2 and QsDRM3 have the C-terminal DNA Mtase domain but only QsDRM2 contains the UBA domain in their N-terminal region (PF00627), essential to its *de novo* methylation activity (Fig 1) [7,8,47]. QsDRM3 does not possess any UBA domain (Fig 1), what is also described for other DRM-like proteins of maize, rice and wild peanut [38,48].

**Figure 1.**
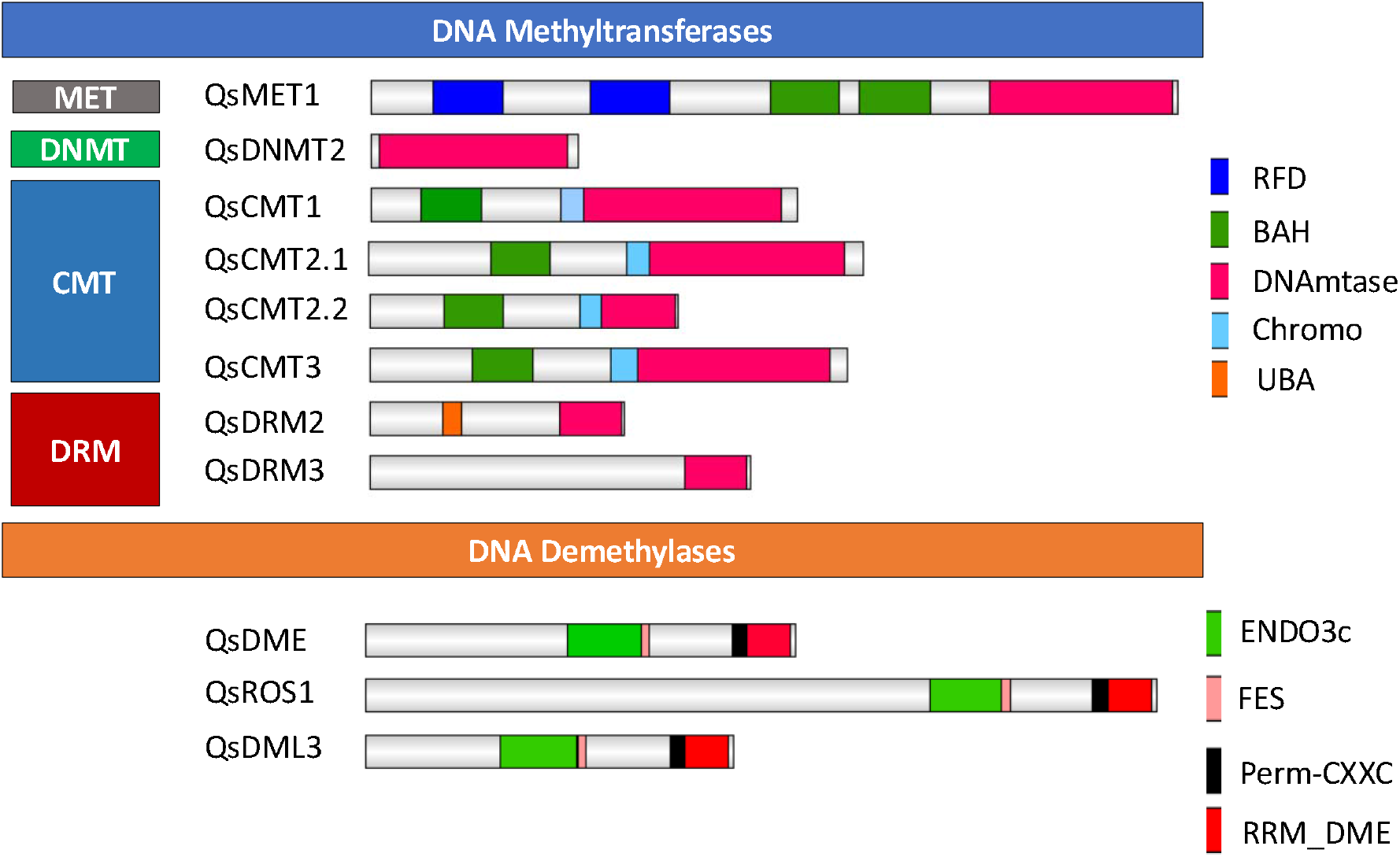
Domain organization of the *Quercus suber* DNA methyltransferases and demethylases proteins. Schematic diagrams show the domain organization of the enzymes. The domain analysis was performed using the NCBI conserved domain and the SMART websites. The proteins and the family they belong to are indicated on the left side of each corresponding schematic diagram. The different conserved protein domains are depicted in different colors as indicated in the legend on the right. C-5 cytosine methyltransferase (IPR001525) is the conserved domain common to all proteins. Replication foci domain RFD (PF12047) and bromo adjacent homology (BAH) domain (PF01426) are present in QsMET1, while chromodomain (CHR) (PF00385) are conserved in chromomethylase proteins. The UBA domain (PF00627) is identified in QsDRM2. The ENDO3c (PF00730), FES (SM000525), Perm-CXXC (PF15629) and RRM_DME (PF15628) domains are present in all DDME.

A phylogenetic analysis was performed (Fig 2) using the amino acid sequences containing the conserved C-5 Mtase domain of the predicted DNA Mtases of *Q. suber* and of other plant species. QsMET1, as well as QsDNMT2, clustered together with homologs orthologs of the closely related Fagaceae species, *Q. robur* and *C. molissima* (Fig 2A). Phylogenetic trees were constructed separately for the CMT (Fig 2B) and DRM (Fig 2C) subfamilies. The analysis suggests that the *Q. suber* CMT2 paralogues might represent duplicated genes. However, only CMT2.1 contains the complete catalytic domain (Fig 1). The *Q. suber* genome lacks a DRM1 homolog like the other Fagaceae family members, *C. molissima* and *Q. robur* (Fig 2C). QsDRM2 groups together with the *A. thaliana* DRM1 and DRM2 proteins in a clade supported with a bootstrap value of 100 (Fig 2C).

**Figure 2.**
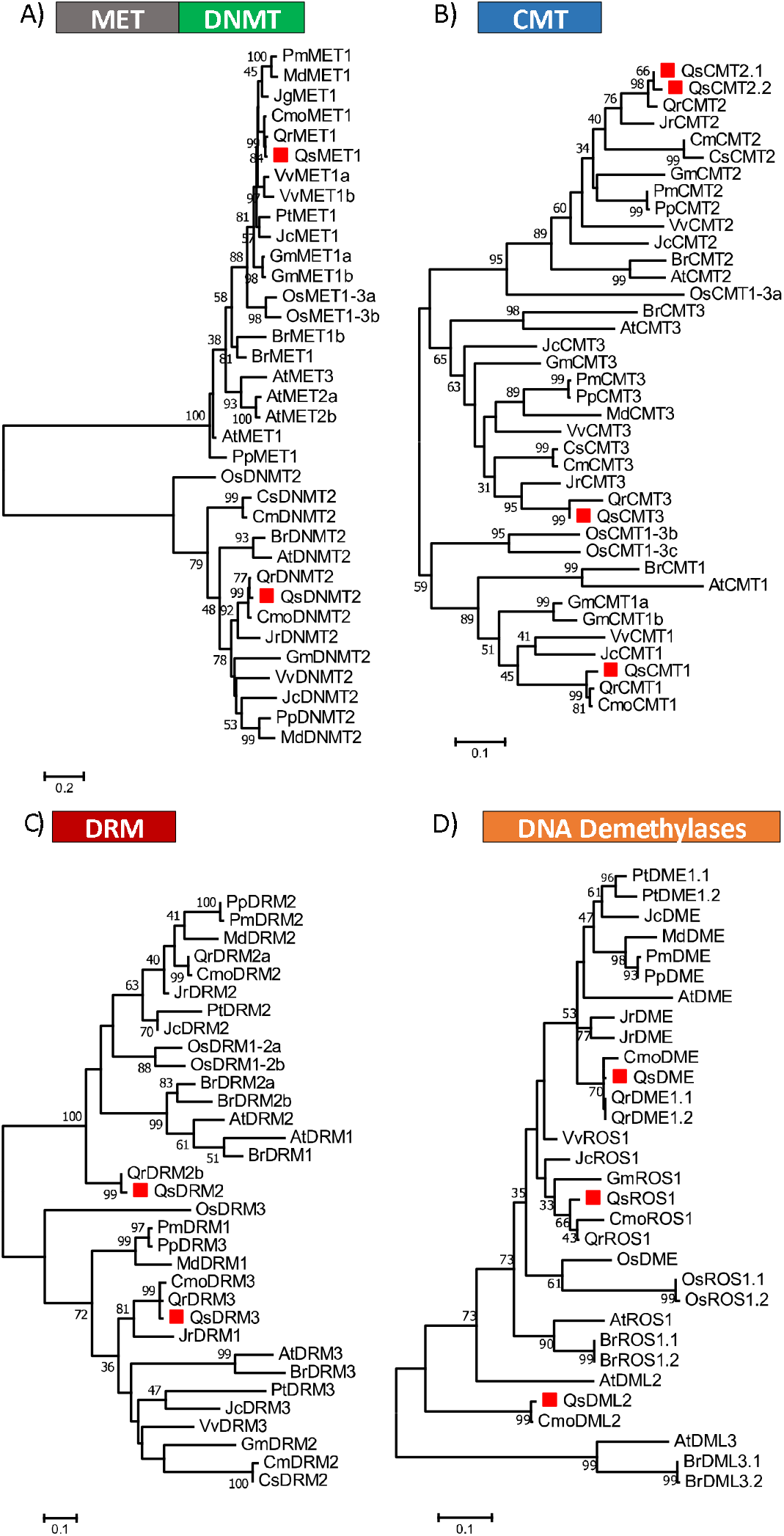
Phylogeny of DNA methyltransferase proteins of A) MET, DNMT, B) CMT family, C) DRM family and of D) DNA Demethylase proteins. The DNA Mtase domain and all DNA demethylase conserved domains of *Quercus suber* (Qs), *Arabidopsis thaliana* (At), *Brassica rapa* (Br), *Glycine max* (Gm), *Jatropha curcas* (Jc), *Populus trichocarpa* (Pt), *Prunus persica* (Pp), *Prunus mume* (Pm), *Vitis vinifera* (Vv), *Cucumis melo* (Cm), *Cucumis sativus* (Cs), *Juglans regia* (Jr), *Castanea mollissima* (Cmo), *Quercus robur* (Qr) and *Oryza sativa* (Os) were aligned using ClustalW and used to infer the evolutionary history using the Maximumlikelihood method. The evolutionary distances (left side scale bar) were computed using the Jones-Taylor-Thornton (JTT) correction model. The numbers at the nodes represent bootstrap values from 1000 replicates. The *Q. suber* proteins are indicated with a red square.

### Identification and classification of *Q. suber* DNA Demethylases

Three *Q. suber* DDME were identified according to their conserved domains: QsDME, QsROS1 and QsDML2 (Fig 1). Four characteristic domains ENDO3c (PF00730), FES (SM000525), Perm-CXXC (PF15629) and RR_ DME (PF15628) are found in all of these three putative DDME proteins (Fig 1), which are involved in base excision DNA repair [11]. A phylogenetic tree was constructed using all DDME proteins from *Q. suber* and from other species (Fig 2D). QsDME and QsROS1 cluster with the respective DME homologs from other Fagaceae trees, all being more closely related to AtROS1, whereas more distantly related to the other DDME members such as QsDML2, AtDML2 and AtDML3 (Fig 2D).

### Identification and classification of *Q. suber* Histone acetyltransferases

Only eight HATs were identified in the *Q. suber* genome based on homology analysis and protein domain identification (Fig 3). Members of the GNAT family are characterized by a Acetyltransf_1 domain (PF00583) [17]. All the members of the GNAT family in Arabidopsis have a homolog in *Q. suber*: QsELP3, QsGCN5 and QsHAG2. QsELP3 and QsGCN5 present two domains each, the ELP3 and BROMO domain, respectively, and the Acetyltransf_1 domain (PF00583). QsHAG2 contains a N-terminal domain HAT1-N (Fig 3) like other homologs in rice, Arabidopsis, soybean [17,49] and a MOZ_SAS domain (PF01853) (Fig 3), a combination not seen in AtHAG2 but already reported in tomato and litchi [49,50]. The *Q. suber* genome contains genes that code for three CBP class proteins, QsHAC1, QsHAC1-like1 and QsHAC1-like2. Domain analysis reveals that the three CBP present the expected KAT11 (PF08214) domain along with several zinc finger type domains (Fig 3). However, the type and the number of zinc fingers differ in the *Q. suber* CBP members. QsHAC1-like2 has not a Transcription Adaptor putative Zinc finger (TAZ) domain (Fig 3). QsHAM1 has the characteristic C-terminal MOZ_SAS domain of MYST family members [17,50,51] (Fig 3). Additionally, QsHAM1 contains an N-terminal CHROMO domain (PF00385) described as able to recognize and bind specific histone residues [52]. One *Q. suber* protein was associated to the HAF family. The QsHAF1 protein contains the characteristic TATA box binding protein (TBP) -binding (PF09247), Ubiquitin (UBQ) (PF00240), ZnF_C2HC (PF01530) [49,50,53] and the BROMO (PF00439) domain in the C-teminal region (Fig 3) that, similarly to the CHROMO domain, is known to bind to acetylated histone lysine residues [54].

**Figure 3.**
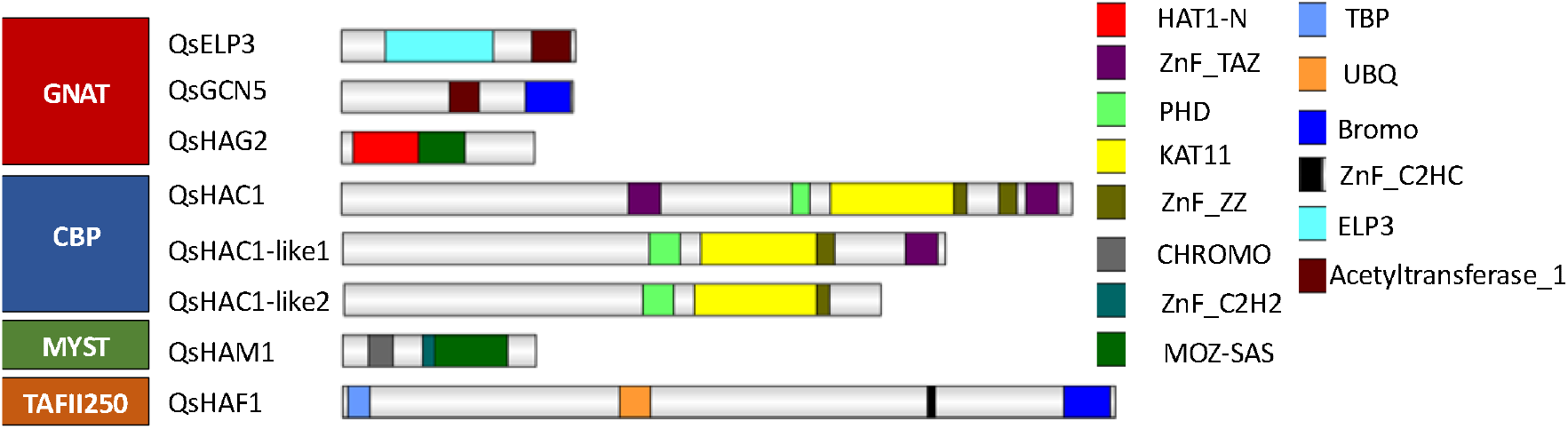
Domain organization of the *Quercus suber* histone acetyltransferases proteins. Schematic diagrams show the domain organization of the enzymes. The domain analysis was performed using the NCBI conserved domain and the SMART websites. The proteins and the family they belong to are indicated on the left side of each corresponding schematic diagram. The different conserved protein domains are depicted in different colors as indicated in the right side legend. Acetyltransferase_1 (PF00583) and C-terminal Bromo (PF00439) domains are conserved domains of QsGCN5, N-terminal ELP3 (IPR006638), and C-terminal Acetyltransferase_1 are domains of QsELP3; N-terminal Hat1_N (PF10394) along with MOZ_SAS (PF01853) are domains of QsHAG2. KAT11 (PF08214), PHD-finger (PF00628), and TAZ (PF02135) are conserved domains of the HAT proteins QsHAC1 and QsHAC1-like1. N-terminal kinase TBP (PF09247), ubiquitin UBQ (PF00240), zinc-finger C2HC (PF01530), and C-terminal bromo are conserved domains of QsHAF1. N-terminal Chromo (PF00385), zinc-finger C2H2 (PF00096), and C-terminal MOZ_SAS domains are typical of QsHAM1.

A phylogenetic tree was generated for each of the four distinct HAT classes, GNAT, CBP, MYST and HAFs, using the Acetyltransf_1 domain (PF00583), the KAT11 domain (PF08214), the MOZ_SAS domain (PF01853) and the UBQ (PF00240) together with the BROMO (PF00439) domain, respectively (Fig 4). All the members of GNAT in Arabidopsis were clearly distinguished in three different clades (Fig 4A). Each homolog is closely related to the homologs of each GNAT subfamily in other species (Fig 4A). Regarding the CBP family, QsHAC1 is more similar to AtHAC1 and OsHAC1 and more distantly related to QsHAC1-like1 (Fig 4B), that is more closely related to AtHAC2 due to the lack of the N-terminal TAZ domain. The higher number of HAC1 copies appear to be common in the majority of the species used to construct the phylogenetic tree (Fig 4B). In contrast, the existence of only one gene encoding the proteins QsHAM1 and QsHAF1 seems to be a common feature also for the other Fagaceae species represented on the phylogenetic tree (Fig 4C and Fig 4D).

**Figure 4.**
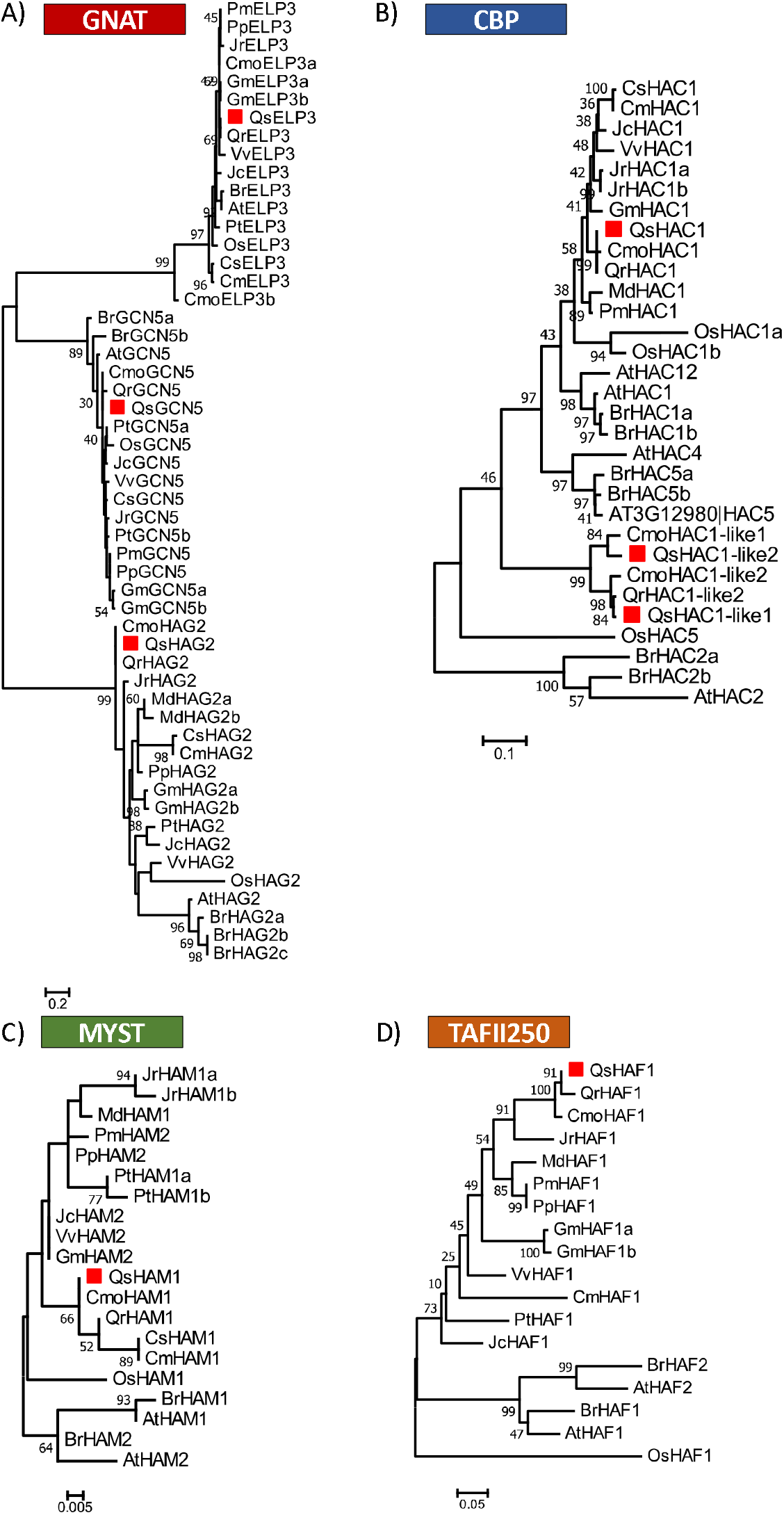
Phylogeny of HAT proteins of GNAT (A), CBP (B), MYST (C) and TAFII250 (D) family. The Acetyltransf_1 domain (GNAT) or the KAT11 domain (CBP), the MOZ_SAS domain (MYST) and the UBQ and BROMO domains (TAFII250) of *Quercus suber* (Qs), *Arabidopsis thaliana* (At), *Brassica rapa* (Br), *Glycine max* (Gm), *Jatropha curcas* (Jc), *Populus trichocarpa* (Pt), *Prunus persica* (Pp), *Prunus mume* (Pm), *Vitis vinifera* (Vv), *Cucumis melo* (Cm), *Cucumis sativus* (Cs), *Juglans regia* (Jr), *Castanea mollissima* (Cmo), *Quercus robur* (Qr) *Oryza sativa* (Os) were aligned using ClustalW and used to infer the evolutionary history using the Maximum-likelihood method. The evolutionary distances (left side scale bar) were computed using the Jones-Taylor-Thornton (JTT) correction model. The numbers at the nodes represent bootstrap values from 1000 replicates. The *Quercus suber* proteins are indicated with a red square.

### Identification and classification of *Q. suber* Histone methyltransferases

Thirty-five proteins of the SET Domain Group (SDG) were identified in *Q. suber*, according to the presence of the SET domain (PF00856) (Fig 5). Although in *A. thaliana* three class I proteins were described (AtCLF, AtSWN and AtMEA), in *Q. suber*, a homolog of AtMEA could not be identified. QsSWN and QsCLF showed a similar domain architecture to Arabidopsis class I proteins since they contained the Swi3, Ada2, N-Cor, the TFIIIB (SANT) domain, C-X(6)-C-X(3)-C-X-C (CXC) domain and SET domain. However, QsSWN and QsCLF did not show the Enhancer of Zeste Domains (EZDs) (Fig 5) [55] similar to the Litchi class I proteins [50]. In *Q. suber*, only four proteins were identified as class II: QsASHH1, QsASHH2, QsASHH3 and QsASHR3. All the *Q. suber* identified proteins have the N-terminal AWS domain (SM00570) in common, preceded by a SET and Post-SET domain (SM00508) (Fig 5), like the Arabidopsis homologs [55,56]. In addition, QsASHR3 has an extra N-terminal PHD domain (PF00628), as reported for other species [49,50,57], and QsASHH2 possess the zinc finger domain CW (PF07496) (Fig 5), which has been reported to bind monomethylated Lysine 4 of Histone 3 (H3K4me1) [58,59]. Five proteins (QsATX2, QsATX3, QsATX5, QsATXR3 and QsATXR7) were classified as class III (Fig 5). All these proteins have the same domain architecture of class III Arabidopsis counterparts [55,56], with the exception of QsATXR7 (Fig 5). QsATX3, QsATX5, QsATX2 and QsATXR7 contain SET and post-SET domains but QsATXR3 contains only the SET domain. The SET domain of QsATX2, QsATX3, QsATX5 proteins is detected along with PHD (PF00628) and PWWP (PF00855) domains in their N-terminal part and in the case of QsATXR7 with a GYF domain (SM00444) (Fig 5). AtATXR3 has also a GYF domain before the SET and Post-SET domains [56], but it was not possible to detect this domain in QsATXR3 because the protein sequence is the only one in this study that is not complete (only identified in the EST database and not in the available genome). In addition, QsATX2 contains the FYRN (SM000541) and FYRC (SM000542) domains (Fig 5). QsATXR5 and QsATXR6, belong to the class IV of HMTs. Both proteins have the N-terminal PHD and the C-terminal SET domains (Fig 5), similarly to the Arabidopsis homologs [56]. Fourteen *Q. suber* proteins were identified as class V HMTs, six SUVH (SU(VAR 3-9) and eight SUVR (SU(VAR) 3-9 related) (Fig 5). All SUVH proteins of *Q. suber* include the preSET (SM000468) and the 5mC-binding motif RING finger-associated (SRA) (SM000466). The SUVR proteins of *Q. suber* proteins contain the predicted domains and the N-terminal WIYLD (PF10440) or C2H2 (PF00096) (Fig 5), that are specific of plants [55,60,61]. In addition, eight other proteins (QsSET41, QsASHR1, QsASHR2-like, QsASHR2, QsATXR2, QsATXR1, QsSET10, QsSET40) containing the SET domain that do not belong to the above-mentioned classes were also analysed and were termed as class VI in this work (Fig 5). QsASHR1 and QsATXR1 possess a TPR (SM00028) domain in the C- and N-terminal part of the protein, respectively. QsSET40 presents a C-terminal RBS (PF09273) domain (Fig 5).

**Figure 5.**
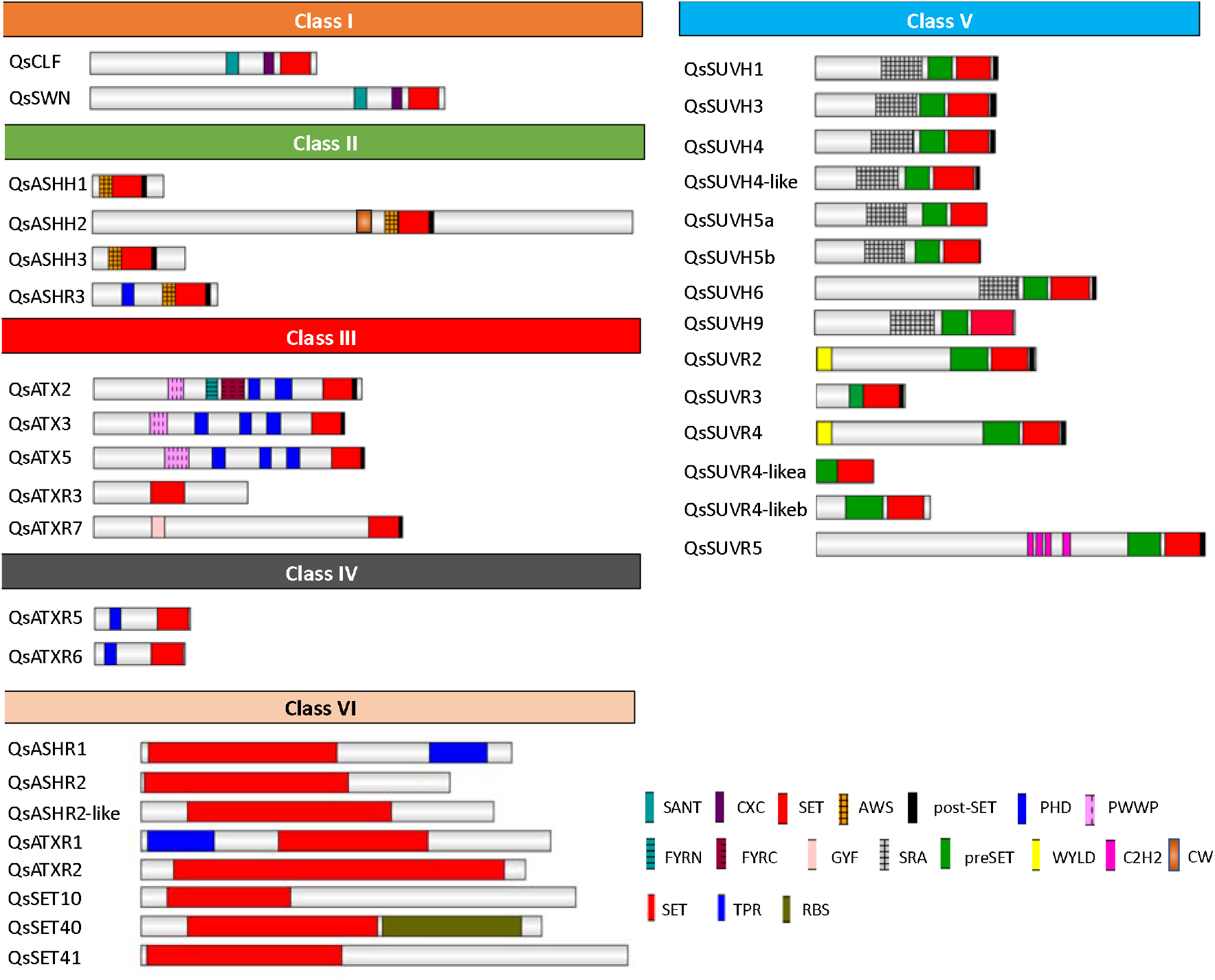
Domain organization of the *Quercus suber* histone methyltransferases proteins. Schematic diagrams show the domain organization of the enzymes. The domain analysis was performed using the NCBI conserved domain and the SMART websites. The protein names and the family they belong to are indicated on the left side and above each corresponding schematic diagram, respectively. The different conserved protein domains are shown in different colors as indicated in the legend on the right side. SANT (SM00717), CXC (PF03638), and SET (PF00856) are conserved domains of Class I. N-terminal AWS (SM00570), SET and Post-SET (SM00508) are conserved domains of Class II. N-terminal PWWP (PF00855), PHD, SET, and Post-SET are conserved domains of some members of Class III. N-terminal GYF (PF02213) is characteristic of QsATXR7. FYRN (SM000541) and FYRC (SM000542) domains are characteristic of QsATX2. N-terminal PHD along with the C-terminal SET are the conserved domains of Class IV. SRA (SM000466), Pre-SET(SM000468), SET, and Post-SET are conserved domains of the SUVH (SU(VAR 3-9) group of Class V. Pre-SET, SET, and Post-SET alongside with the N-terminal WIYLD (PF10440), or C2H2 (PF00096) or absence of domain are characteristic of Class V SUVR (SU(VAR) 3-9 related) group. Some members of the other SET domain containing proteins comprised N-terminal or C-terminal TPR domain, while other included the C-terminal RBS domain.

Phylogenetic analysis of the five HMT classes was performed. The class I proteins QsCLF and QsSWN clustered in two different subclades (Supplementary Figure S5A). The class IV proteins QsATXR5 and QsATXR6, also grouped in different clades (Supplementary Figure S5B). The class II proteins QsASHH1, QsASHH2, QsASHH3 and QsASHR3 were separated in different clades supported by bootstrap values of 100, 100, 80 and 99, respectively (Supplementary Figure S6A). The class III proteins QsATX3, QsATX5 clustered in the same clade, whereas QsATX2, QsATXR3 and QsATXR7 clustered in different clades each (Supplementary Figure S6B). Regarding class V, QsSUVH1 and QsSUVH3 belong to the clade containing the SUVH1/3/7/8 homolog proteins, positioned in different branches closely related do *Q. robur* and *C. molissima* homologs, supported by a 98 bootstrap (Supplementary Figure S7). QsSUVH9 grouped together with SUVH2 and SUVH9 homologs in another well supported clade (bootstrap value: 99). One *Q. suber* protein homolog to SUVH6 and two proteins homologs to SUVH5 were positioned in another clade. QsSUVH5 is duplicated in *Q. suber* and no homolog was found in the other Fagaceae trees *C. molissima* and *Q. robur* (Supplementary Figure S7). Two *Q. suber* proteins cluster in the SUVH4 clade. SUVH4 is duplicated in the two Quercus species but also in the annual plant *Glycina max*. Apart from these eight proteins belonging to the SUVH clade of class V, other six proteins belong to the SUVR clade (Supplementary Figure S7). QsSUVR5 and QsSUVR6 were positioned with their closest homologs in distinct clades, while one QsSUVR2 and three SUVR4-like grouped together with their respective homologs in the same clade (Supplementary Figure S7). Each class VI protein was cluster together with their closest homologs in other species (Supplementary Figure S8).

### Identification and classification of *Q. suber* Histone Demethylases

Twenty-six *Q. suber* proteins were identified as HDMTs (Fig 6A). HDMTs are divided in two main families: Lysine specific demethylase 1 (KDM/LSD1) and Jumonji C domain containing proteins (JMJ).

The amino-oxidase (PF01593) and SWIRM (PF04433) domains characteristic of the four proteins that belong to KDM1/LSD1 family: QsLDL1, QsLDL2, QsLDL3 and QsFLD (Fig 6A) were used to generate a phylogenetic tree for the KDM1/LSD1 family (Fig 6B). The four proteins were separated in four distinct clades together along with their closest homologs in other species (Fig 6B).

**Figure 6.**
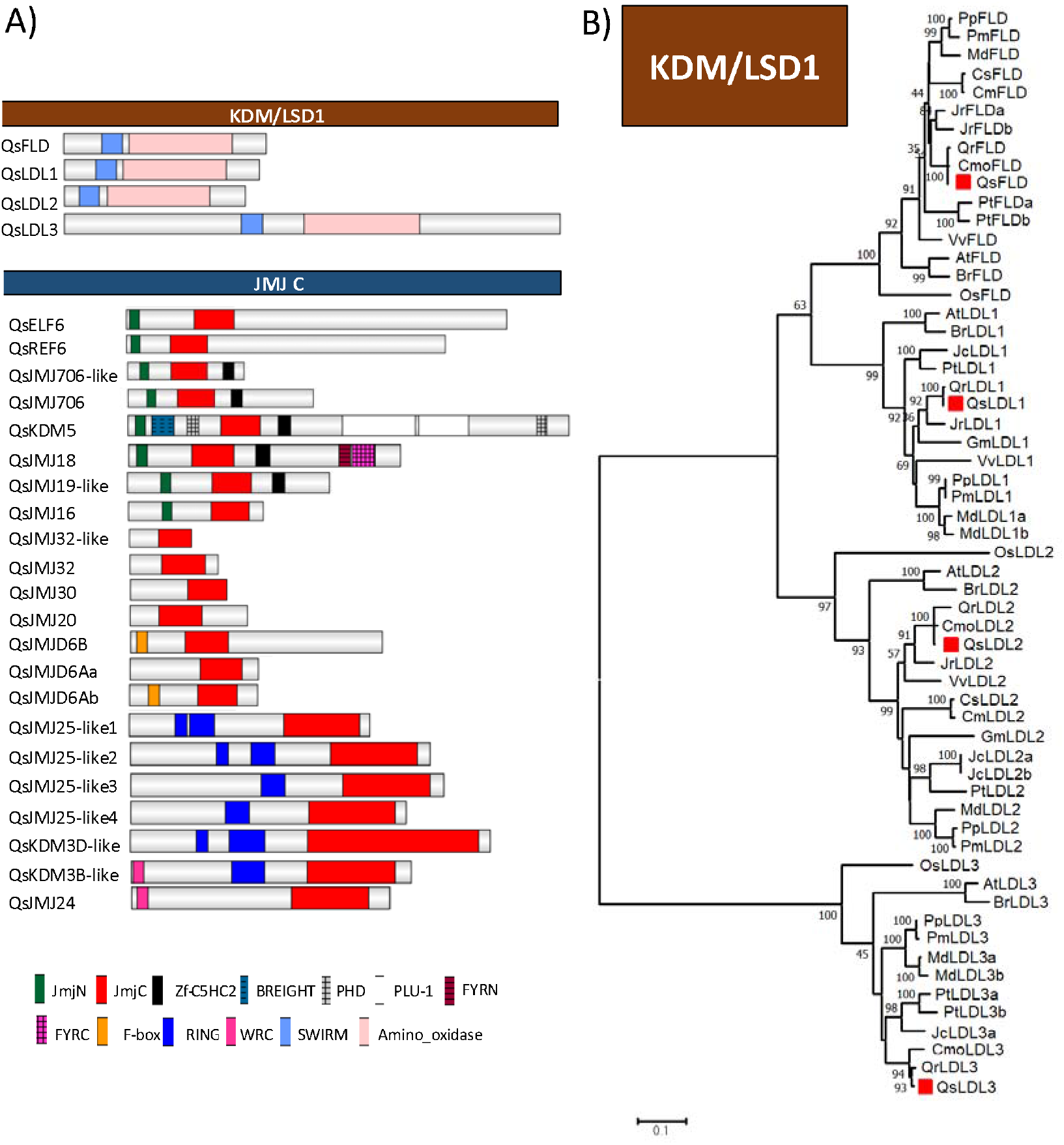
*Quercus suber* Histone Demethylases proteins with their A) domain organization and phylogeny of the B) KDM/LSD1 family. A) Schematic diagrams show the domain organization of the enzymes. The domain analysis was performed using the NCBI conserved domain and the SMART websites. The proteins and the family they belong to are indicated on left side and above each corresponding schematic diagram, respectively. The different conserved protein domains are depicted in different colors as indicated in the legend at the bottom. RING-finger (IPR001841), WRC (PF08879) and JmjC (PF02373) are conserved domains of the KDM3/JMJD1 class proteins. N-terminal JmjN (PF02375), JmjC, and the C-terminal C5HC2 (PF02928) domains are conserved typical of KDM4/JMJD2 proteins. N-terminal F-box (PF00646), and C-terminal JmjC are conserved domains of the JMJD6 class. KDM/LSD1 proteins contain only the Amino oxidase (IPR002937) and SWIRM (IPR007526) domains. B) The amino-oxidase and SWIRM domains of *Quercus suber* (Qs), *Arabidopsis thaliana* (At), *Brassica rapa* (Br), *Glycine max* (Gm), *Jatropha curcas* (Jc), *Populus trichocarpa*(Pt), *Prunus persica* (Pp), *Prunus mume* (Pm), *Vitis vinifera* (Vv), *Cucumis melo* (Cm), *Cucumis sativus* (Cs), *Juglans regia* (Jr), *Castaneaa mollissima* (Cmo), *Quercus robur* (Qr) *Oryza sativa* (Os) were aligned using ClustalW and used to infer the evolutionary history using the Maximum-likelihood method. The evolutionary distances (left side scale bar) were computed using the Jones-Taylor-Thornton (JTT) correction model. The numbers at the nodes represent bootstrap values from 1000 replicates. The *Quercus suber* proteins are indicated with a red square.

All the *Q. suber* JMJ proteins contain the JmjC domain (PF02373) (Fig 6). QsELF6, QsREF6, QsJMJ706, QsJMJ706-like, QsKDM5, QsJMJ18, QsJMJ19-like and QsJMJ16 have in common the JmjC and JmjN domains (PF02375) (Fig 6A). QsJMJ25, QsKDM3D- and QsKDM3B-like present an N-terminal RING finger domain (IPR001841). Both QsJMJ24 and QsKDM3B possess an N-terminal WRC (PF08879) domain. QsJMJ32, QsJMJ32-like, QsJMJ30 and QsJMJ20 proteins do not have any other domain other than the JmjC domain (Fig 6A), while QsJMJD6B and QsJMJD6Ab contain an N-terminal F-box domain (Fig 6A).

The amino acid sequence of the JmJC domain (PF02373) was used to generate one phylogenetic tree for the JMJ family (Supplementary Figure S9). In Arabidopsis, the JMJ family is divided into four well known classes: KDM4/JMJD2, KDM5/JARID, KDM3 and KDM2 [62]. The first clade (KDM4/JMJD2) contains the members of the sub-clades PKDM9, PKDM8: QsREF6, QsELF5, QsJMJ706 and QsJMJ706-like. The second clade (KDM5/JARID) is represented by KDM5 and PKDM7 members: QsKDM5, QsJMJ18, QsJMJ19-like and QsJMJ16 (Supplementary Figure S9). The *Q. suber* genome encodes four JMJ25-like proteins, one JMJ24-, one KDM3D- and one QsKDM3B-like proteins belonging to the KDM3/JMJD1 family, clustering in the same clade (Supplementary Figure S9). The same number of JMJ25 copies found in the *Q. suber* genome was also found in the genome of the biennial plant *B. rapa*. The fourth clade contains the remaining sub-clades PKDM12 (QsJMJ32, QsJMJ32-like and QsJMJ30), PKDM11 (QsJMJ20) and JMJD6 (QsJMJD6A and QsJMJD6B) (Supplementary Figure S9). JMJ32 is duplicated in the two Quercus species as well as in the perennial *P. trichocarpa* (Supplementary Figure S9).

### Identification and classification of *Q. suber* Histone Deacetylases

HDACs are divided into three main groups: HD2-type, Sirtuin and RPD3/HDA1. Twenty-two *Q. suber* proteins were identified as HDACs-like: three HD2-like proteins, six sirtuin-like proteins and 13 RPD3/HDA1 group proteins (Fig 7A).

**Figure 7.**
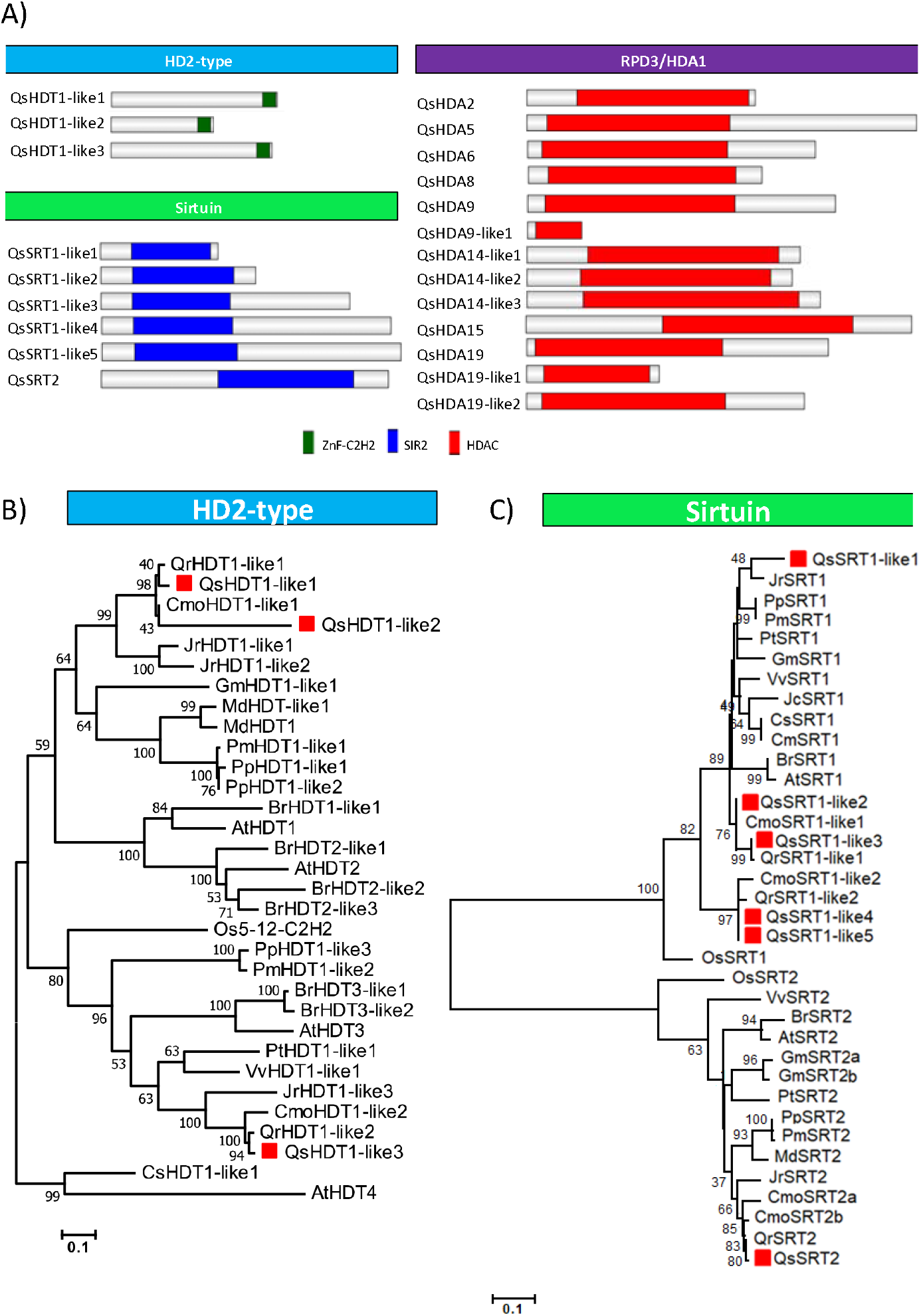
Domain organization A) of HDAC proteins in *Quercus suber* and the phylogeny of B) HD2 and C) Sirtuin family. A) The domain analysis was performed using the NCBI conserved domain and the SMART web site. The proteins and the family they belong to are indicated on left side and above each corresponding schematic diagram, respectively. The different conserved protein domains are depicted in different colors as indicated in the right-side legend. The HDAC domain (PF00850) is the conserved domain of RPD3/HDA1 proteins. C-terminal zinc finger domain C2H2 (PF00096) is typical in the HD2-type proteins. The SIR2 proteins contain an SIR2 domain (PF02146). B) All the protein of HD2-type (HD2 family) and C) SIR2 domain (Sirtuin family) of *Quercus suber* (Qs), *Arabidopsis thaliana* (At), *Brassica rapa* (Br), *Glycine max* (Gm), *Jatropha curcas* (Jc), *Populus trichocarpa* (Pt), *Prunus persica* (Pp), *Prunus mume* (Pm), *Vitis vinifera* (Vv), *Cucumis melo* (Cm), *Cucumis sativus* (Cs), *Juglans regia* (Jr), *Castanea mollissima* (Cmo), *Quercus robur* (Qr) *Oryzasativa* (Os) were aligned using ClustalW and used to infer the evolutionary history using the Maximumlikelihood method. The evolutionary distances (left side scale bar) were computed using the Jones-Taylor-Thornton (JTT) correction model. The numbers at the nodes represent bootstrap values from 1000 replicates. The *Quercus suber* proteins were indicated with a red square. Phylogenetic analyses were conducted in MEGA7.

Sirtuin-like group members are characterized by the presence of the SIR2 domain (PF02146), while RPD3/HDA1 members show only the HDAC domain (PF00850). Five genes code for QsSRT1-like proteins while only one code for QsSRT2 (Fig 7A).

The phylogenetic tree of HD2-type proteins was generated using the alignment of the conserved ZnF-C2H2 regions (Fig 7B). QsHDT1-like1 and QsHDT1-like2 group in the same clade that AtHDT1 and AtHDT2, while QsHDT1-like3 is more phylogenetically related with AtHDT3. Fig 7C illustrates the divergence of the Sirtuin family proteins based on a phylogenetic tree generated through the SIR2 domain alignment. A higher number of SRT1 copies were found in the Fagaceae genomes, with *Q. robur* and *C. molissima* having two copies, and the other species having a single copy gene.

Supplementary Figure S10 shows a phylogenetic tree illustrating the relationship among the RPD3/HDA1 superfamily proteins, produced by aligning their HDAC domains. In the first clade it was noticeable a higher number of copies of HDA19 homologs (three) that are not exclusive of Quercus species as other plants such as *V. vinifera*, *M. domestica*, *O. sativa* and *P. trichocarpa* have also a high number of copies. In contrast, three copies of HDA14 were exclusively found in *Q. suber*.

### Gene expression of epigenetic regulators during oak plant development

To infer the putative function of the identified enzymes under study, their transcript levels were analyzed in RNAseq experiments representing five *Q. suber* tissues in different developmental stages/conditions: acorns from developmental stage 2, 3/4 and 5 [25]; a pool of embryos collected from the acorns of stage 1 to [25]; cork of bad and good quality [24]; male and female flowers [27]; well-watered roots and roots with medium and severe drought [28]; red and opened buds and dormant and swollen buds [29]. The transcriptomic analysis of six tissues (root, eco-dormant bud, swelling bud, leaf, in vitro dedifferentiated callus and secondary differentiating xylem), previously done by Lesur and collaborators (2015) for the phylogenetically close species *Q. robur* were also mined to detect the expression of these regulators (Table S2).

A heatmap with the normalized expression data (Table S2) was generated for each family of epigenetic modifier enzymes. The DNA mtases *QsMET1* and *QsCMT3* gene were detected mainly in buds, acorns and roots (Fig 8, Table S). The closest homolog to *QsMET1* (*QrMET1*) was highly expressed in swelling buds when compared with the other tissues (Table S2). *QsDRM2* had highest expression in cork and buds. The DDME *QsROS1* was more expressed in roots and acorns while *QsDME* was preferentially expressed in acorns at the first developmental stages (S2, S3/S4). The HAT genes *QsHAC1, QsHAF1 and QsHAM1* showed high expression in acorns and in roots when comparing with flowers and buds. The class I HMT gene *QsCLF* was more expressed in acorns and roots (Fig 8). The class III HMT genes *QsATX3* and *QsATX5* were expressed in different tissues but remarkably in the intermediate stages (S3/S4) of fruit development (Fig 8). *QsATX2* had higher expression in the intermediate stages (S3/S4) of fruit development, similarly to *QsATX3*, but was also detected in roots, where its expression was higher during severe drought stress (Fig 8). The class II gene *QsASHH2* was highly expressed during different development stages of fruit development and in embryos (Fig 8). The class V gene *QsSUVH4* expression is higher in red and opened buds, in bad quality cork and in female flowers when compared with the other HMT genes (Fig 8). Comparing the different conditions of watering in *Q. suber* roots the other class III HMT gene *QsATXR7* was more expressed in roots under severe drought, however *QrATXR7* appears to be more expressed in *Q. robur* ecodormant buds (Table S2). In *Q. suber* buds and flowers, a higher accumulation of the HDAC *QsJMJ706-like* and *QsREF6* transcripts was observed, when comparing to all the other genes (Fig 8). Among all *Q. suber* RPD3/HDA1-like members, *QsHDA9* was less expressed in good quality cork, *QsHDA15* was less expressed in last stages of male flower development, *QsHDA5* was more expressed in the last stage of fruit development while *QsHDA2* was more expressed in the early stages (Fig 8). The HD2-type *QsHDT1-like1* was expressed at high levels in all the tissues analyzed in *Q. suber* when compared with the other putative HDAC genes (Fig 8). *Q. robur* homolog *QrHDT1-like1* is also the HDAC gene with highest expression (Table S2).

**Figure 8.**
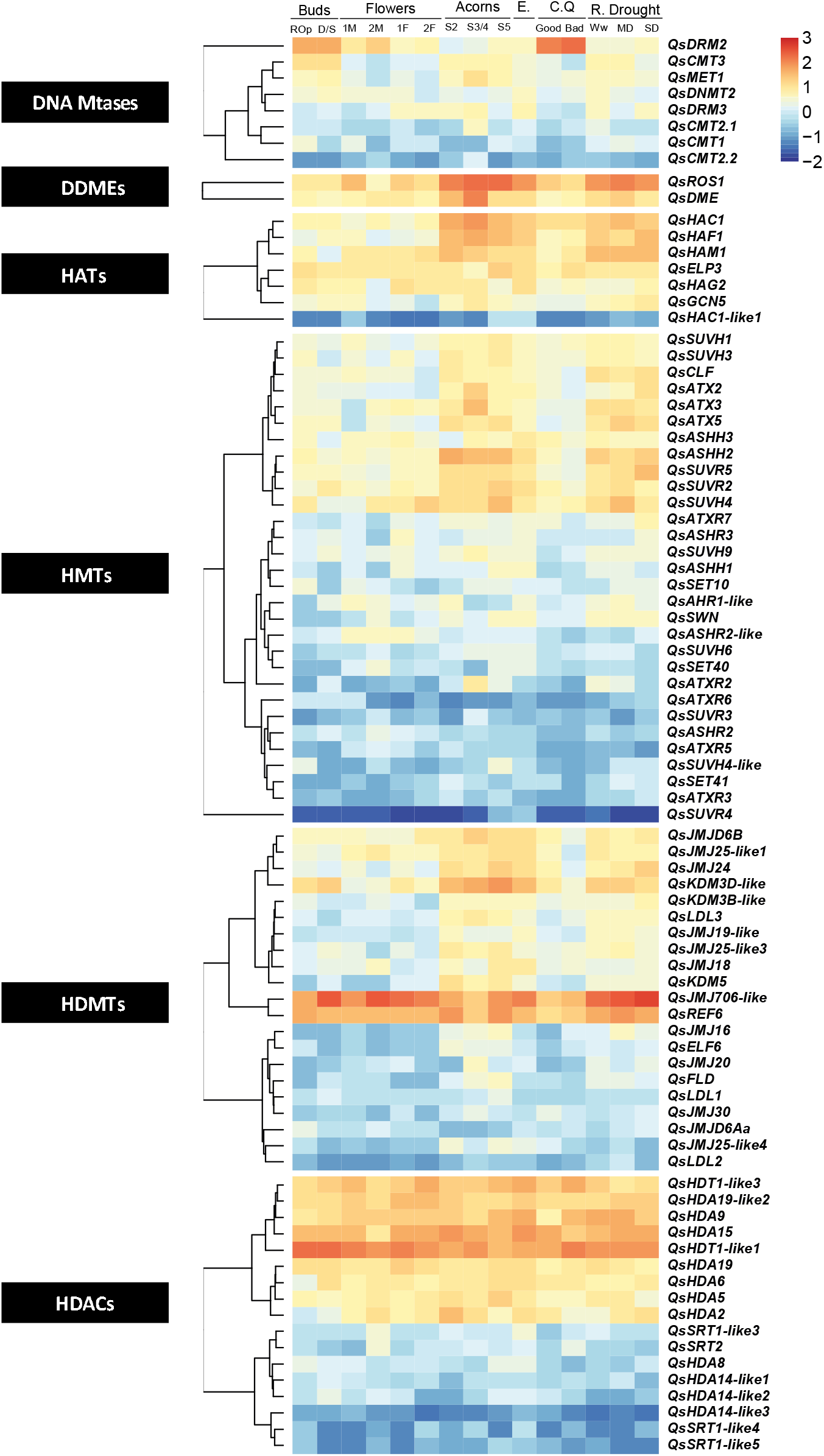
Expression profiles of the homologs of DNA methyltransferases, DNA demethylases,, histone acetyltransferases, histone methyltransferases, histone acetylases and histone demethylases in *Quercus suber*. The expression levels of the epigenetic regulators identified in diverse *Q. suber* databases were analyzed after mapping the 454 reads derived of the RNA sequencing of several tissues (categorized on the top) to the *Quercus suber* genome and the corresponding read count normalization. The tissues analyzed include: acorns in 3 development stages (S2, S3/4, S5), embryos (E), good and bad quality cork (C.Q), early (1F and 1M) and last stages (2F and 2M) of female (F) and male flower (M) development, roots (R.) with medium (MD), severe (SD) or without drought stress (Ww), samples with red and open buds (ROp) and samples with dormant and swollen buds (D/S). The results were plotted in an heatmap after Rlog transformation followed by Z-score computation. For interpretation of the expression patterns color, the chart is next to each heatmap. The gene name code was indexed in the heatmap, based on the phylogenetic analysis results.

## Discussion

Due to the recent publication of the *Q. suber* genome [63], it is now possible to identify the complete list of genes encoding chromatin packaging regulators in this species (Table S1). In this study we were able to identify eight DNA Mtases, three DNA demethylases, thirty-five HMTs, twenty-six HDMTs, eight HATs and twenty-two HDACs. In general, all the phylogenetic trees generated in this work are consistent with previous works in other species [17,38,65–67,39,41,42,48–50,53,64].

*Q. suber*, like other perennials, and in contrast to annual species, must adapt its vegetative and reproductive growth every year to fluctuating environmental conditions that occur over the different seasons. Individuals sense the decreasing photoperiod and temperature and anticipate the winter period by adjusting their own physiology to initiate a rest period. The bud dormancy period during the winter is a good example of a process that has been shown to be dependent of epigenetic gene silencing in several species [68,69]. The cold-induced establishment of trimethylation of lysine 27 of histone 3 (H3K27me3) in known regulators of dormancy, such as *DORMANCY-ASSOCIATED MADS-BOX* (*DAM*) genes, have been reported in several perennial species [68,69], which in turn, may affect the activity of bud bust and flowering inductor genes, such as *FLOWERING LOCUS T* (*FT*) [70–73]. The HMT members of the polycomb repressive complex (PRC) 2, responsible for H3K27me3 deposition), such as CLF are likely to regulate growth-dormancy transitions [74,75]. In *Q. suber* buds it was not possible to differentiate the expression of *QsCLF* between flowering inductive and non-inductive conditions because dormant and swollen buds were pooled together in the data of the transcriptomic study analyzed in this work [29].

Gene silencing by H3K27 methylation and by DNA methylation is essential in order to regulate genome stability by blocking transposable elements [77,78]. The establishment of these epigenetic marks is a result of the redundant action of ATXR5 and ATXR6 (H3K27me) and specific DNA Mtases. In *Q. suber*, *QsATXR5* and *QsATXR6* transcripts are less abundant in roots, acorns and embryos when compared with the other genes encoding HMTs (Table S3). The transcripts of *QsMET1* and *QsCMT3* genes were detected mainly in actively proliferating cell tissues such as buds (with flower meristems inside), acorns and roots (Fig 8, Table S3). The phylogenetic tree positioning and the conserved domain architecture of all the QsCMTs and QsMET1 proteins (Fig 1 and 2) suggests similar functions to the Arabidopsis homologs. So, *QsMET1* and *QsCMT3* may play a role in maintaining DNA methylation during DNA replication ensuring their correct transmission during subsequent cell divisions [47,78]. Maintaining DNA methylation is a decisive mechanism for *Q. suber* that develops new shoots every year to reinforce the canopy and to produce flowers and fruits ensuring reproductive success. Interestingly, in *Q. suber* the lower expression level of the presumed DNA methylation maintenance genes *QsCMT-* and *QsMET-* like is counterbalanced by the higher expression of the putative *de novo* DNA methylation gene *QsDRM2*. This result was already observed in previous works [78,79]. *QsDRM2* higher expression is well noticed in cork and buds (containing vegetative and reproductive tissues) (Fig 8, Table S3). Previous studies have proven that AtDRM2 exhibited *de novo* methylation activity through its UBA domain [7,8,47] QsDRM2 is phylogenetically close to AtDRM2 (Fig 2) and contains the UBA domain (Fig 1). So, the activation of putative *de novo* methylation gene *QsDRM2* may be associated with the establishment of new DNA methylation patterns in these tissues [9]. DNA methylation is in close association with H3K9 methylation, through SUVH proteins [81,82,83,84] due to the SRA domain that works as a 5mC-binding motif [55,56]. In *Q. suber*, eight proteins were identified as belonging to the SUVH clade (Fig 8), and all of them contain the domain structure reported for the Arabidopsis proteins. AtSUVH4 is the best known protein of the SUVH group and is involved mainly in the maintenance of cytosine methylation in a non-CG context [84,85]. DNA methylation variability may contribute to cork cell characteristics linked to quality [86]. Inácio et al. (2018) emphasized the presence of both DNA and H3K9 methylation silencing pathways in cork. Here, we show that *QsSUVH4* is expressed in cork (Fig 8) but also in the other tissues analyzed. The deacetylation of histones has also a negative role in gene expression. The *Q. suber* homologs of HDACs, *QsHDA9* and *QsHDA15* were highly expressed in several tissues but specific downregulation was found in good quality cork and in the last stages of male flower development, respectively. *QsHDA5* and *QsHDA2* are less expressed in the overall tissues, but were more expressed in the last and early stages of fruit development, respectively. AtHDA9 and AtHDA5 modulate flowering time [87,88] while AtHDA15 has been shown to be a crucial element of photomorphogenesis [90]. The impact of histone deacetylation catalyzed by AtHDA2 is not fully understood. It is possible that these enzymes in *Q. suber* may have the same roles as the Arabidopsis counterparts but also could function in other developmental processes specific of *Q. suber*, such as in cork formation and acorn production.

Lysine methylation of histones is also linked to gene expression activation, which is catalyzed by class II proteins, by methylation of Lysine 36 or 4 of histone 3 (H3K36/H3K4). AtASHH2 has been reported to control flowering induction leading to increased transcription of the flowering-repressor *FLOWERING LOCUS C* (*FLC*) by both H3K36 methylation [91,92] and H3K4 trimethylation [92]. AtASHH2 is essential for proper expression of pivotal genes throughout reproductive development stages, such as floral organ identity or embryo sac and pollen development [93]. *QsASHH2* is highly expressed during different development stages of fruit development and embryos (Fig 8). Since QsASHH2 groups together with the AtASHH2 (Supplementary Figure S6A) and contains similar domains (Fig 5), it likely performs similar functions. The class III HMT proteins like ATX1, ATX2, ATXR3 and ATXR7 are also involved in gene activation pathways [94–96]. AtATXR3 is required for appropriate root growth [97] and *QsATXR3* expression was higher in roots subjected to severe drought (Figure 14). *QsATXR7* and *QsATX2* transcripts were also detected in roots and its expression were higher during severe drought stress (Figure 14), suggesting a putative role in dehydration stress. In contrast, the expression of *QsATXR2* in roots was higher in well watering conditions decreasing with the increase of drought severity. The crosstalk between gene activation pathways and the environment is necessary to promote the expression of genes involved in abiotic stress resistance. The Mediterranean climate, in which *Q. suber* subsists, is very sensitive to hydrological changes such as drought and, as a consequence, the roots should adapt to supply the tree with its water and nutrient demands. In Arabidopsis, ATXR2 promotes accumulation of H3K36me3 enhancing transcription and is mainly related with the control of molecular components implicated in root organogenesis [98]. The expression differences of *QsATXR3, QsATXR7, QsATX2* and *QsATXR2* in contrastive watering condition of *Q. suber* roots suggest a putative role in root development, particularly during dehydration stress. Other genes whose proteins are involved in the deposition of euchromatic marks such as the MYST-like gene *QsHAM1* and the TAFII250 *QsHAF1* have also a strong expression in roots. High levels of expression for both genes were also observed in acorns at different developmental stages (Fig 8). Interestingly, in tomato *HAF1* homolog also has a strong expression in fruit and roots [49] and the *HAM1* homolog is more expressed in flowers and in very immature fruits [49]. In Arabidopsis, single *ham1* or *ham2* mutant plants have a wild-type phenotype, whilst *ham1ham2* double mutation caused mitotic defects in the mega and microgametophyte development [51]. So, a possible role for *QsHAM1* and *QsHAF1* in reproductive development should be considered.

Phylogenetic analysis allowed us to evidence the diversification of epigenetic regulators among the *Q. suber* genome, showing several cases of duplications such is the case of CMT2, SUVH5, SUVH4 and JMJ32 with two copies, HDA19, HAC1 and HDA14 with three copies, JMJ25 with four copies, and remarkably, SRT1 with five. The phylogenetic analysis showed that JMJ32 is not only duplicated in *Q. suber* but also in *Q. robur* and in *P. trichocarpa* in contrast with the other species used in the phylogenetic analysis that contain only JMJ32 single-copy genes (Supplementary Figure S9). Qian *et al*. (2015) reported the specificity of JMJ32 as a single copy gene in plants with the exception of a duplication of the *P. trichocarpa* JMJ32. Like oaks, *P. trichocarpa* is a woody perennial tree that requires several years to reach maturity, when cyclical transitions between vegetative and flowering phases occur. In Arabidopsis, JMJ32 mediates the demethylation of H3K27me3 in the flowering-repressor *FLC* locus, preventing premature early flowering under warm temperatures [100]. It is tempting to suggest that JMJ32 duplications may have evolved to overcome special requirements of perennials regarding flowering time control.

More than one SRT1 gene copy was found only in Fagaceae genomes (Fig 7). SRT1 is required for transposon repression and to regulate the expression of genes involved in plant stress-response and programmed cell death [101,102]. Unfortunately, it was not possible to obtain any clue about *QsSRT1* functional divergence since only *QsSRT1-like3* was significantly expressed in the *Q. suber* tissues analyzed in this work (more expressed in last stages of male flower development, in embryos and in well-watered roots) (Fig 8, Table S3). The closest homolog of *QsSRT1* in *Q. robur* (*QrSRT1-like1*) was not detected in any tissue analyzed but *QrSRT1-like2* had higher expression in roots when compared to other tissues. The higher number of SRT1 proteins in oaks could be related with genus-specific traits such as their high longevity and essential plasticity. These trees need to have the competence to modulate their own growth (by gene transcription regulation) during more than two hundred years in order to face multiple biotic and abiotic stresses. The duplication of CMT2, HDA14 and SUVH5 was only found in *Q. suber* (Fig 2B, Supplementary Figure S10 and Supplementary Figure S7). Moreover, no SUVH5 homologs were found in *C. mollissima* and *Q. robur*. CMT2 acts in the protection against genome instability by silencing transposons located in the pericentromeric regions of genomes [9,83,103,104]. HDA14 is involved in photosynthesis by regulating the expression of RuBisCO activase under low-light conditions [105]. SUVH5 acts in transposon silencing by methylation of histone H3 at Lysine 9 (H3K9me) and by mediating DNA methylation in non-CG sequences [106]. If the function of CMT2 and SUVH5 is indeed conserved in *Q. suber*, their duplication may have evolved due the important role of these enzymes in transposon silencing.

The high number of proteins here described and their expression profile in distinct developmental stages suggest an important role for these regulators in cork oak development. The transcriptomic data here represented is, however, not a full representation of the *Q. suber* transcriptome. In the future, the expression should be analyzed using a whole set of plant organs, at different developmental stages and collected at the same conditions to be accurately compared. The identification of a vast set of *Q. suber* epigenetic regulators in this study should greatly facilitate this type of expression analysis. Here, their likely roles were predicted based on the expression levels but also based on the phylogenetic analysis and domain architecture. The domain annotation is an important and valuable tool for discovering the homologs of protein families in plant species in which the functional characterization by mutant analysis is impossible, like in *Q. suber*. The next challenge should be to know if these genes have a similar function to the well characterized Arabidopsis homologs, and to study whether changes in their expression is the cause of an adaptation process.

## Acknowledgments

This work was supported by the FEDER funds through the Operational Competitiveness Programme-COMPETE and by National Funds through FCT—Fundação para a Ciência e a Tecnologia under the projects PTDC/AGR-GPL/118508/2010, “Characterization of Reproductive Development of Quercus suber” and PTDC/AGR-FOR/3356/2014, “Characterization of cork formation and reproductive biology in a cork oak hybrids population” and the PhD grant ref. SFRH/BD/111529/2015. Work supported by UID/MULTI/04046/2013 and UID/AGR/04129/2013 center grant from FCT, Portugal (to BioISI and LEAF, respectively).

## References

[1] Y. Gruenbaum, T. Naveh-Many, H. Cedar, A. Razin, Sequence specificity of methylation in higher plant DNA, Nature. 292 (1981) 860–862.

[2] A.J. Bewick, R.J. Schmitz, Gene body DNA methylation in plants, Curr Opin Plant Biol. (2017) 103–110. https://doi.org/10.1016/j.pbi.2016.12.007.Gene.

[3] X. Zhang, J. Yazaki, A. Sundaresan, S. Cokus, S.W.L. Chan, H. Chen, I.R. Henderson, P. Shinn, M. Pellegrini, S.E. Jacobsen, J.R.R. Ecker, Genome-wide High-Resolution Mapping and Functional Analysis of DNA Methylation in Arabidopsis, Cell. 126 (2006) 1189–1201. https://doi.org/10.1016/j.cell.2006.08.003.

[4] T. Chen, E. Li, Structure and Function of Eukaryotic DNA Methyltransferases, in: Curr. Top. Dev. Biol, 2004: pp. 55–89. http://arxiv.org/abs/quant-ph/9609014v1%5Cnhttp://www.ncbi.nlm.nih.gov/pubmed/15861675.

[5] S. Cokus, S. Feng, X. Zhang, Z. Chen, B. Merriman, C. Haudenschild, S. Pradhan, S. Nelson, M. Pellegrini, S. Jacobsen, Shotgun bisulfite sequencing of the Arabidopsis genome reveals DNA methylation patterning, Nature. 452 (2008) 215*219. https://doi.org/10.3390/ijms151222874.

[6] R. Lister, R.C.O. Malley, J. Tonti-Filippini, B.D. Gregory, C.C. Berry, A.H. Millar, J.R. Ecker, R.C. O’Malley, J. Tonti-Filippini, B.D. Gregory, C.C. Berry, A.H. Millar, J.R. Ecker, Highly Integrated Single-Base Resolution Maps of the Epigenome in Arabidopsis, Cell. 133 (2008) 523–536. https://doi.org/10.1016/j.cell.2008.03.029.Highly.

[7] X. Cao, S.E. Jacobsen, Locus-specific control of asymmetric and CpNpG methylation by the DRM and CMT3 methyltransferase genes, Proc. Natl. Acad. Sci. 99 (2002) 16491–16498. https://doi.org/10.1073/pnas.162371599.

[8] J.A. Law, S.E. Jacobsen, Establishing, maintaining and modifying DNA methylation patterns in plants and animals, Nat Rev Genet. 11 (2010) 204–220. https://doi.org/10.1038/nrg2719.Establishing.

[9] X. Cao, S. Jacobsen, Role of the Arabidopsis DRM methyltransferases in de novo DNA methylation and gene silencing, Curr. Biol. 12 (2002) 1138–1144. https://doi.org/10.1016/S0960-9822(02)00925-9.

[10] U. Naumann, L. Daxinger, T. Kanno, C. Eun, Q. Long, Z.J. Lorkovic, M. Matzke, A.J.M. Matzke, Genetic evidence that DNA methyltransferase DRM2 has a direct catalytic role in RNA-directed DNA methylation in Arabidopsis thaliana, Genetics. 187 (2011) 977–979. https://doi.org/10.1534/genetics.110.125401.

[11] Z. Gong, T. Morales-Ruiz, R.R. Ariza, T. Roldán-Arjona, L. David, J.K. Zhu, ROS1, a repressor of transcriptional gene silencing in Arabidopsis, encodes a DNA glycosylase/lyase., Cell. 111 (2002) 803–14. http://www.ncbi.nlm.nih.gov/pubmed/12526807.

[12] W. Xiao, M. Gehring, Y. Choi, L. Margossian, H. Pu, J.J. Harada, R.B. Goldberg, R.I. Pennell, R.L. Fischer, Imprinting of the MEA Polycomb Gene Is Controlled by Antagonism between MET1 Methyltransferase and DME Glycosylase, Dev. Biol. 5 (2003) 891–901.

[13] M. Gehring, J.H. Huh, T.F. Hsieh, J. Penterman, Y. Choi, J.J. Harada, R.B. Goldberg, R.L. Fischer, DEMETER DNA glycosylase establishes MEDEA polycomb gene self-imprinting by allele-specific demethylation, Cell. 124 (2006) 495–506. https://doi.org/10.1016/j.cell.2005.12.034.

[14] J. Penterman, D. Zilberman, J.H. Huh, T. Ballinger, S. Henikoff, R.L. Fischer, DNA demethylation in the Arabidopsis genome., PNAS. 104 (2007) 6752–6757. https://doi.org/10.1007/BF00507750.

[15] T.R. Hebbes, A.W. Thorne, C. Crane-Robinson, A direct link between core histone acetylation and transcriptionally active chromatin., EMBO J. 7 (1988) 1395–1402. https://doi.org/10.1002/j.1460-2075.1988.tb02956.x.

[16] D.E. Sterner, S.L. Berger, Acetylation of Histones and Transcription-Related Factors, Microbiol. Mol. Biol. Rev. 64 (2000) 435–459. https://doi.org/10.1128/mmbr.64.2.435-459.2000.

[17] R. Pandey, A. Mu, È. Ller, C.A. Napoli, D.A. Selinger, C.S. Pikaard, E.J. Richards, J. Bender, D.W. Mount, R.A. Jorgensen, Analysis of histone acetyltransferase and histone deacetylase families of Arabidopsis thaliana suggests functional diversification of chromatin modification among multicellular eukaryotes, Nucleic Acids Res. 30 (2002) 5036–5055. https://www.ncbi.nlm.nih.gov/pmc/articles/PMC137973/pdf/gkf660.pdf.

[18] D.W.K. Ng, T. Wang, M.B. Chandrasekharan, R. Aramayo, S. Kertbundit, T.C. Hall, Plant SET domain-containing proteins: Structure, function and regulation, Biochim. Biophys. Acta - Gene Struct. Expr. 1769 (2007) 316–329. https://doi.org/10.1016/j.bbaexp.2007.04.003.

[19] Y. Shi, F. Lan, C. Matson, P. Mulligan, J.R. Whetstine, P.A. Cole, R.A. Casero, Y. Shi, Histone Demethylation Mediated by the Nuclear Amine Oxidase Homolog LSD1 instance, histone H3 K9 (H3-K9) methylation is associ-ated with heterochromatin formation (Nakayama et al and also, Sidney Kimmel Compr. Cancer Cent. 119 (2004) 941–953. https://doi.org/10.1016/j.cell.2004.12.012.

[20] Y.I. Tsukada, J. Fang, H. Erdjument-Bromage, M.E. Warren, C.H. Borchers, P. Tempst, Y. Zhang, Histone demethylation by a family of JmjC domain-containing proteins, Nature. 439 (2006) 811–816. https://doi.org/10.1016/j.landurbplan.2017.03.007.

[21] R.J. Klose, Y. Zhang, Regulation of histone methylation by demethylimination and demethylation, Nat. Rev. Mol. Cell Biol. 8 (2007) 307–318. https://doi.org/10.1038/nrm2143.

[22] J.D. Thompson, D.G. Higgins, T.J. Gibson, CLUSTAL W: Improving the sensitivity of progressive multiple sequence alignment through sequence weighting, position-specific gap penalties and weight matrix choice, Nucleic Acids Res. 22 (1994) 4673–4680. https://doi.org/10.1093/nar/22.22.4673.

[23] B.G. Hall, Building phylogenetic trees from molecular data with MEGA, Mol. Biol. Evol. 30 (2013) 1229–1235. https://doi.org/10.1093/molbev/mst012.

[24] J.B. Pereira-Leal, I.A. Abreu, C.S. Alabaça, M.H. Almeida, P. Almeida, T. Almeida, M.I. Amorim, S. Araújo, H. Azevedo, A. Badia, D. Batista, A. Bohn, T. Capote, I. Carrasquinho, I. Chaves, A.C. Coelho, M.M.R. Costa, R. Costa, A. Cravador, C. Egas, C. Faro, A.M. Fortes, A.S. Fortunato, M.J. Gaspar, S. Gonçalves, J. Graça, M. Horta, V. Inácio, J.M. Leitão, T. Lino-Neto, L. Marum, J. Matos, D. Mendonça, A. Miguel, C.M. Miguel, L. Morais-Cecílio, I. Neves, F. Nóbrega, M.M. Oliveira, R. Oliveira, M.S. Pais, J.A.P. Paiva, O.S. Paulo, M. Pinheiro, J.A.P. Raimundo, J.C. Ramalho, A.I. Ribeiro, T. Ribeiro, M. Rocheta, A.I. Rodrigues, J.C. Rodrigues, N.J.M. Saibo, T.E. Santo, A.M. Santos, P. Sá-Pereira, M. Sebastiana, F. Simões, R.S. Sobral, R. Tavares, R. Teixeira, C. Varela, M.M. Veloso, C.P.P. Ricardo, A comprehensive assessment of the transcriptome of Cork oak (Quercus suber) through EST sequencing, BMC Genomics. 15 (2014) 1–14. https://doi.org/10.1186/1471-2164-15-371.

[25] A. Miguel, J. de Vega-Bartol, L. Marum, I. Chaves, T. Santo, J. Leitão, M.C. Varela, C.M. Miguel, Characterization of the cork oak transcriptome dynamics during acorn development, BMC Plant Biol. 15 (2015). https://doi.org/10.1186/s12870-015-0534-1.

[26] R.T. Teixeira, A.M. Fortes, C. Pinheiro, H. Pereira, Comparison of good- and bad-quality cork: Application of high-throughput sequencing of phellogenic tissue, J. Exp. Bot. 65 (2014) 4887–4905. https://doi.org/10.1093/jxb/eru252.

[27] M. Rocheta, R. Sobral, J. Magalhães, M.I. Amorim, T. Ribeiro, M. Pinheiro, C. Egas, L. Morais-Cecílio, M.M.R. Costa, Comparative transcriptomic analysis of male and female flowers of monoecious Quercus suber., Front. Plant Sci. 5 (2014) 1–16. https://doi.org/10.3389/fpls.2014.00599.

[28] A.P. Magalhães, N. Verde, F. Reis, I. Martins, D. Costa, T. Lino-Neto, P.H. Castro, R.M. Tavares, H. Azevedo, RNA-seq and gene network analysis uncover activation of an ABA-dependent signalosome during the cork oak root response to drought, Front. Plant Sci. 6 (2016) 1–17. https://doi.org/10.3389/fpls.2015.01195.

[29] A. Usié, F. Simões, P. Barbosa, B. Meireles, I. Chaves, S. Gonçalves, A. Folgado, M.H. Almeida, J. Matos, A.M. Ramos, Comprehensive analysis of the cork oak (Quercus suber) transcriptome involved in the regulation of bud sprouting, Forests. 8 (2017) 1–19. https://doi.org/10.3390/f8120486.

[30] A. Criscuolo, S. Brisse, AlienTrimmer: A tool to quickly and accurately trim off multiple short contaminant sequences from high-throughput sequencing reads, Genomics. 102 (2013) 500–506. https://doi.org/10.1016/j.ygeno.2013.07.011.

[31] R.K. Shrestha, B. Lubinsky, V.B. Bansode, M.B.J. Moinz, G.P. McCormack, S.A. Travers, QTrim: A novel tool for the quality trimming of sequence reads generated using the Roche/454 sequencing platform, BMC Bioinformatics. 15 (2014). https://doi.org/10.1186/1471-2105-15-33.

[32] H. Li, R. Durbin, Fast and accurate short read alignment with Burrows–Wheeler transform, Mass Genomics. 25 (2009) 1754–1760. https://doi.org/10.1093/bioinformatics/btp324.

[33] Y. Liao, G.K. Smyth, W. Shi, FeatureCounts: An efficient general purpose program for assigning sequence reads to genomic features, Bioinformatics. 30 (2014) 923–930. https://doi.org/10.1093/bioinformatics/btt656.

[34] M.I. Love, W. Huber, S. Anders, Moderated estimation of fold change and dispersion for RNA-seq data with DESeq2, Genome Biol. 15 (2014) 1–21. https://doi.org/10.1186/s13059-014-0550-8.

[35] I. Lesur, G. Le Provost, P. Bento, C. Da Silva, J.C. Leplé, F. Murat, S. Ueno, J. Bartholomé, C. Lalanne, F. Ehrenmann, C. Noirot, C. Burban, V. Léger, J. Amselem, C. Belser, H. Quesneville, M. Stierschneider, S. Fluch, L. Feldhahn, M. Tarkka, S. Herrmann, F. Buscot, C. Klopp, A. Kremer, J. Salse, J.M. Aury, C. Plomion, The oak gene expression atlas: Insights into Fagaceae genome evolution and the discovery of genes regulated during bud dormancy release, BMC Genomics. 16 (2015). https://doi.org/10.1186/s12864-015-1331-9.

[36] S. Kumar, X. Cheng, S. Klimasauskas, M. Sha, J. Posfai, R.J. Roberts, G.G. Wilson, The DNA (cytosine-5) methyltransferases, Nucleic Acids Res. 22 (1994) 1–10. https://doi.org/10.1093/nar/22.1.1.

[37] J. Pósfai, A.S. Bhagwat, G. Pósfai, R.J. Roberts, Predictive motifs derived from cytosine methyltransferases, Nucleic Acids Res. 17 (1989) 2421–2435. https://doi.org/10.1093/nar/17.7.2421.

[38] A. Pavlopoulou, S. Kossida, Plant cytosine-5 DNA methyltransferases: Structure, function, and molecular evolution, Curr. Top. Dev. Biol. 90 (2007) 530–541. https://doi.org/10.1016/j.ygeno.2007.06.011.

[39] D. Cao, Z. Ju, C. Gao, X. Mei, D. Fu, H. Zhu, Y. Luo, B. Zhu, Genome-wide identification of cytosine-5 DNA methyltransferases and demethylases in Solanum lycopersicum, Gene. 550 (2014) 230–237. https://doi.org/10.1016/j.gene.2014.08.034.

[40] L. Chang, Z. Zhang, B. Han, H. Li, H. Dai, P. He, H. Tian, Isolation of dna-methyltransferase genes from strawberry (fragaria 3 ananassa duch.) and their expression in relation to micropropagation, Plant Cell Rep. 28 (2009) 1373–1384. https://doi.org/10.1007/s00299-009-0737-8.

[41] R. Garg, R. Kumari, S. Tiwari, S. Goyal, Genomic survey, gene expression analysis and structural modeling suggest diverse roles of DNA methyltransferases in legumes, PLoS One. 9 (2014). https://doi.org/10.1371/journal.pone.0088947.

[42] S. Gianoglio, A. Moglia, A. Acquadro, C. Comino, E. Portis, The genome-wide identification and transcriptional levels of DNA methyltransferases and demethylases in globe artichoke, PLoS One. 12 (2017) 1–17. https://doi.org/10.1371/journal.pone.0181669.

[43] R. Sharma, R.K. Mohan Singh, G. Malik, P. Deveshwar, A.K. Tyagi, S. Kapoor, M. Kapoor, Rice cytosine DNA methyltransferases - Gene expression profiling during reproductive development and abiotic stress, FEBS J. 276 (2009) 6301–6311. https://doi.org/10.1111/j.1742-4658.2009.07338.x.

[44] J. Xu, H. Xu, Q. Xu, X. Deng, Characterization of DNA methylation variations during fruit development and ripening of sweet orange, Plant Mol Biol Rep. 33 (2015). https://doi.org/10.1007/s11105-014-0732-2.

[45] J. Eissenberg, Molecular biology of the chromo domain: an ancient chromatin module comes of age., Gene. 275 (2001) 19–29. http://www.wormbase.org/db/misc/paper?name=WBPaper00012893.

[46] I. Callebaut, J.C. Courvalin, J.P. Mornon, The BAH (bromo-adjacent homology) domain: A link between DNA methylation, replication and transcriptional regulation, FEBS Lett. 446 (1999) 189–193. https://doi.org/10.1016/S0014-5793(99)00132-5.

[47] X. Cao, W. Aufsatz, D. Zilberman, M.F. Mette, M.S. Huang, M. Matzke, S.E. Jacobsen, Role of the DRM and CMT3 Methyltransferases in RNA-Directed DNA Methylation, Curr. Biol. 13 (2003) 2212–2217. https://doi.org/10.1016/j.cub.2003.11.052.

[48] P. Wang, C. Gao, X. Bian, S. Zhao, C. Zhao, H. Xia, H. Song, L. Hou, S. Wan, X. Wang, Genome-Wide Identification and Comparative Analysis of Cytosine-5 DNA Methyltransferase and Demethylase Families in Wild and Cultivated Peanut, Front. Plant Sci. 7 (2016) 1–14. https://doi.org/10.3389/fpls.2016.00007.

[49] R. Aiese Cigliano, W. Sanseverino, G. Cremona, M.R. Ercolano, C. Conicella, F.M. Consiglio, Genome-wide analysis of histone modifiers in tomato: Gaining an insight into their developmental roles, BMC Genomics. 14 (2013). https://doi.org/10.1186/1471-2164-14-57.

[50] M. Peng, P. Ying, X. Liu, C. Li, R. Xia, J. Li, M. Zhao, Genome-Wide Identification of Histone Modifiers and Their Expression Patterns during Fruit Abscission in Litchi, Front. Plant Sci. 8 (2017) 1–16. https://doi.org/10.3389/fpls.2017.00639.

[51] D. Latrasse, M. Benhamed, Y. Henry, S. Domenichini, W. Kim, D.X. Zhou, M. Delarue, The MYST histone acetyltransferases are essential for gametophyte development in arabidopsis, BMC Plant Biol. 8 (2008) 1–16. https://doi.org/10.1186/1471-2229-8-121.

[52] X. De La Cruz, S. Lois, S. Sánchez-Molina, M.A. Martínez-Balbás, Do protein motifs read the histone code?, BioEssays. 27 (2005) 164–175. https://doi.org/10.1002/bies.20176.

[53] X. Liu, M. Luo, W. Zhang, J. Zhao, J. Zhang, K. Wu, L. Tian, J. Duan, Histone acetyltransferases in rice (Oryza sativa L.): phylogenetic analysis, subcellular localization and expression, BMC Plant Biol. 12 (2012) 1–17. https://doi.org/10.1186/1471-2229-12-145.

[54] R. Marmorstein, S.L. Berger, Structure and function of bromodomains in chromatin-regulating complexes, Gene. 272 (2001) 1–9. https://doi.org/10.1118/1.2761137.

[55] F. Pontivianne, T. Blevins, C.S. Pikaard, Arabidopsis Histone Lysine Methyltransferases, Adv Bot Res. 2296 (2010) 1–18. https://doi.org/10.1016/S0065-2296(10)53001-5.Arabidopsis.

[56] N. Springer, C. Napoli, D. Selinger, R. Pandey, K. Cone, V. Chandler, H. Kaeppler, S. Kaeppler, Comparative Analysis of SET Domain Proteins in Maize and Arabidopsis Reveals Multiple Duplications Preceding the Divergence of Monocots and Dicots, Plant Physiol. 132 (2003) 907–925. https://doi.org/10.1053/bega.2002.0324.

[57] Y. Huang, C. Liu, W.H. Shen, Y. Ruan, Phylogenetic analysis and classification of the Brassica rapa SET-domain protein family, BMC Plant Biol. 11 (2011) 175. https://doi.org/10.1186/1471-2229-11-175.

[58] V. Hoppmann, T. Thorstensen, P.E. Kristiansen, S.V. Veiseth, M.A. Rahman, K. Finne, R.B. Aalen, R. Aasland, The CW domain, a new histone recognition module in chromatin proteins, EMBO J. 30 (2011) 1939–1952. https://doi.org/10.1038/emboj.2011.108.

[59] Y. Liu, Y. Huang, Uncovering the mechanistic basis for specific recognition of monomethylated H3K4 by the CW domain of Arabidopsis histone methyltransferase SDG8, J. Biol. Chem. 293 (2018) 6470–6481. https://doi.org/10.1074/jbc.RA117.001390.

[60] T. Thorstensen, A. Fischer, S. V. Sandvik, S.S. Johnsen, P.E. Grini, G. Reuter, R.B. Aalen, The Arabidopsis SUVR4 protein is a nucleolar histone methyltransferase with preference for monomethylated H3K9, Nucleic Acids Res. 34 (2006) 5461–5470. https://doi.org/10.1093/nar/gkl687.

[61] L. Baumbusch, T. Thorstensen, V. Krauss, A. Fischer, K. Naumann, R. Assalkhou, The Arabidopsisthalianagenome contains at least 29 active genes encoding SET domain proteins that can be assigned to four evolutionarily conserved classes, 29 (2001) 4319–4333. https://doi.org/10.1093/nar/29.21.4319.

[62] Zhou, Ma, Evolutionary history of histone demethylase families: distinct evolutionary patterns suggest functional divergenc, MC Evol. Biol. 21 (2008) 210–214. https://doi.org/10.1186/1471-2148-8-294.

[63] A. Ramos, A. Usié, P. Barbosa, P.M. Barros, T. Capote, I. Chaves, F. Simões, I. Abreu, I. Carrasquinho, C. Faro, J.B. Guimarães, D. Mendonça, F. Nóbrega, L. Rodrigues, N.J.M. Saibo, M.C. Varela, C. Egas, J. Matos, C.M. Miguel, M.M. Oliveira, C.P. Ricardo, S. Gonçalves, The draft genome sequence of cork oak, Sci. Data. 5 (2018) 1–12. https://doi.org/10.1038/sdata.2018.69.

[64] G. Malik, M. Dangwal, S. Kapoor, M. Kapoor, Role of DNA methylation in growth and differentiation in Physcomitrella patens and characterization of cytosine DNA methyltransferases, FEBS J. 279 (2012) 4081–4094. https://doi.org/10.1111/febs.12002.

[65] F. Aquea, T. Timmermann, P. Arce-Johnson, Analysis of histone acetyltransferase and deacetylase families of Vitis vinifera, Plant Physiol. Biochem. 48 (2010) 194–199. https://doi.org/10.1016/j.plaphy.2009.12.009.

[66] L.C. Liew, M.B. Singh, P.L. Bhalla, An RNA-Seq Transcriptome Analysis of Histone Modifiers and RNA Silencing Genes in Soybean during Floral Initiation Process, PLoS One. 8 (2013) 1–23. https://doi.org/10.1371/journal.pone.0077502.

[67] D. Papaefthimiou, E. Likotrafiti, A. Kapazoglou, K. Bladenopoulos, A. Tsaftaris, Epigenetic chromatin modifiers in barley: III. Isolation and characterization of the barley GNAT-MYST family of histone acetyltransferases and responses to exogenous ABA, Plant Physiol. Biochem. 48 (2010) 98–107. https://doi.org/10.1016/j.plaphy.2010.01.002.

[68] D.P. Horvath, S. Sung, D. Kim, W. Chao, J. Anderson, Characterization, expression and function of DORMANCY ASSOCIATED MADS-BOX genes from leafy spurge, Plant Mol. Biol. 73 (2010) 169–179. https://doi.org/10.1007/s11103-009-9596-5.

[69] C. Leida, A. Conesa, G. Llácer, M.L. Badenes, G. Ríos, Histone modifications and expression of DAM6 gene in peach are modulated during bud dormancy release in a cultivar-dependent manner, New Phytol. 6 (2012) 67–80.

[70] X. Hao, W. Chao, Y. Yang, D. Horvath, Coordinated expression of FLOWERING LOCUS T and DORMANCY ASSOCIATED MADS-BOX-like genes in leafy spurge, PLoS One. 10 (2015) 1–18. https://doi.org/10.1371/journal.pone.0126030.

[71] A. Ito, T. Saito, D. Sakamoto, T. Sugiura, S. Bai, T. Moriguchi, Physiological differences between bud breaking and flowering after dormancy completion revealed by DAM and FT/TFL1 expression in Japanese pear (Pyrus pyrifolia), Tree Physiol. 36 (2015) 109–120. https://doi.org/10.1093/treephys/tpv115.

[72] Q. Niu, J. Li, D. Cai, M. Qian, H. Jia, S. Bai, S. Hussain, G. Liu, Y. Teng, X. Zheng, Dormancy-associated MADS-box genes and microRNAs jointly control dormancy transition in pear (Pyrus pyrifolia white pear group) flower bud, J. Exp. Bot. 67 (2016) 239–257. https://doi.org/10.1093/jxb/erv454.

[73] R.K. Singh, J.P. Maurya, A. Azeez, P. Miskolczi, S. Tylewicz, K. Stojkovic, N. Delhomme, V. Busov, R.P. Bhalerao, A genetic network mediating the control of bud break in hybrid aspen, Nat. Commun. 9 (2018) 1–10. https://doi.org/10.1038/s41467-018-06696-y.

[74] A. Lomax, D.P. Woods, Y. Dong, F. Bouché, Y. Rong, K.S. Mayer, X. Zhong, R.M. Amasino, An ortholog of CURLY LEAF/ENHANCER OF ZESTE like-1 is required for proper flowering in Brachypodium distachyon, Plant J. 93 (2018) 871–882. https://doi.org/10.1111/tpj.13815.

[75] T. Ruttink, M. Arend, K. Morreel, V. Storme, S. Rombauts, J. Fromm, R.P. Bhalerao, W. Boerjan, A. Rohde, A Molecular Timetable for Apical Bud Formation and Dormancy Induction in Poplar, Plant Cell Online. 19 (2007) 2370–2390. https://doi.org/10.1105/tpc.107.052811.

[76] Y. Jacob, S. Feng, C. LeBlanc, Y. Bernatavichute, H. Stroud, S. Cokus, L. Johnson, M. Pellegrini, S. Jacobsen, S. Michaels, ATXR5 and ATXR6 are novel H3K27 monomethyltransferases required for chromatin structure and gene silencing, Nat Struct Mol Biol. 2009. 16 (2009) 763–768. https://doi.org/10.1038/nsmb.1611.ATXR5.

[77] H. Saze, O.M. Scheid, J. Paszkowski, Maintenance of CpG methylation is essential for epigenetic inheritance during plant gametogenesis, Nat. Genet. 34 (2003) 65–69. https://doi.org/10.1038/ng1138.

[78] M. Ramos, M. Rocheta, L. Carvalho, V. Inácio, J. Graça, L. Morais-Cecilio, Expression of DNA methyltransferases is involved in Quercus suber cork quality, Tree Genet. Genomes. 9 (2013) 1481–1492. https://doi.org/10.1007/s11295-013-0652-6.

[79] V. Inácio, M.T. Martins, J. Graça, L. Morais-Cecílio, Cork Oak Young and Traumatic Periderms Show PCD Typical Chromatin Patterns but Different Chromatin-Modifying Genes Expression, Front. Plant Sci. 9 (2018) 1–18. https://doi.org/10.3389/fpls.2018.01194.

[80] L.M. Johnson, X. Cao, S.E. Jacobsen, Interplay between Two Epigenetic Marks: DNA Methylation and Histone H3 Lysine 9 Methylation, Curr. Biol. 12 (2002) 1360–1367. https://doi.org/10.1016/S0960-9822(02)00976-4.

[81] L. Johnson, M. Bostick, X. Zhang, E. Kraft, I. Henderson, J. Callis, S. Jacobsen, The SRA methyl-cytosine-binding domain links DNA and histone methylation, Curr Biol. 17 (2007) 379–384. https://doi.org/10.1038/jid.2014.371.

[82] L. Johnson, J. Law, A. Khattar, I.R. Henderson, S.E. Jacobsen, SRA-domain proteins required for DRM2-mediated de novo DNA methylation, PLoS Genet. 4 (2008) 1–13. https://doi.org/10.1371/journal.pgen.1000280.

[83] J. Jackson, Lindroth, AM, X. Cao, S. Jacobsen, Control of CpNpG DNA methylation by the KRYPTONITE histone H3 methyltransferase, Nature. 416 (2002) 556–560.

[84] F. Malagnac, L. Bartee, J. Bender, An Arabidopsis SET domain protein required for maintenance but not establishment of DNA methylation, 21 (2002).

[85] J. Du, L.M. Johnson, M. Groth, F. Suhua, J. Christopher, S. Li, A. Vashisht, J. Wohlschlegel, D. Patel, S.E. Jacobsen, Mechanism of DNA methylation-directed histone methylation by KRYPTONITE, Mol Cell. 55 (2014) 495–504. https://doi.org/10.1111/j.1746-1561.2010.00581.x.Are.

[86] V. Inácio, P.M. Barros, A. Costa, C. Roussado, E. Gonçalves, R. Costa, J. Graça, M.M. Oliveira, L. Morais-Cecílio, Differential DNA methylation patterns are related to phellogen origin and quality of Quercus suber cork, PLoS One. 12 (2017) 1–18. https://doi.org/10.1371/journal.pone.0169018.

[87] Y.J. Kim, R. Wang, L. Gao, D. Li, C. Xu, H. Mang, J. Jeon, X. Chen, X. Zhong, J.M. Kwak, B. Mo, L. Xiao, X. Chen, POWERDRESS and HDA9 interact and promote histone H3 deacetylation at specific genomic sites in Arabidopsis, Proc. Natl. Acad. Sci. U. S. A. 113 (2016) 14858–14863. https://doi.org/10.1073/pnas.1618618114.

[88] M. Luo, R. Tai, C.W. Yu, S. Yang, C.Y. Chen, W.D. Lin, W. Schmidt, K. Wu, Regulation of flowering time by the histone deacetylase HDA5 in Arabidopsis, Plant J. 82 (2015) 925–936. https://doi.org/10.1111/tpj.12868.

[89] X. Liu, C.-Y. Chen, K.-C. Wang, M. Luo, R. Tai, L. Yuan, M. Zhao, S. Yang, G. Tian, Y. Cui, H.-L. Hsieh, K. Wu, PHYTOCHROME INTERACTING FACTOR3 Associates with the Histone Deacetylase HDA15 in Repression of Chlorophyll Biosynthesis and Photosynthesis in Etiolated Arabidopsis Seedlings, Plant Cell. 25 (2013) 1258–1273. https://doi.org/10.1105/tpc.113.109710.

[90] Z. Zhao, Y. Yu, D. Meyer, C. Wu, W.H. Shen, Prevention of early flowering by expression of FLOWERING LOCUS C requires methylation of histone H3 K36, Nat. Cell Biol. 7 (2005) 1156–1160. https://doi.org/10.1038/ncb1329.

[91] L. Xu, Z. Zhao, A. Dong, L. Soubigou-Taconnat, J.-P. Renou, A. Steinmetz, W.-H. Shen, Di- and Tri-but Not Monomethylation on Histone H3 Lysine 36 Marks Active Transcription of Genes Involved in Flowering Time Regulation and Other Processes in Arabidopsis thaliana, Mol. Cell. Biol. 28 (2008) 1348–1360. https://doi.org/10.1128/MCB.01607-07.

[92] S.Y. Kim, Y. He, Y. Jacob, Y.S. Noh, S. Michaels, R. Amasino, Establishment of the Vernalization-Responsive, Winter-Annual Habit in Arabidopsis Requires a Putative Histone H3 Methyl Transferase, Plant Cell Online. 17 (2005) 3301–3310. https://doi.org/10.1105/tpc.105.034645.

[93] P.E. Grini, T. Thorstensen, V. Alm, G. Vizcay-Barrena, S.S. Windju, T.S. Jørstad, Z.A. Wilson, R.B. Aalen, The ASH1 HOMOLOG 2 (ASHH2) histone H3 methyltransferase is required for ovule and anther development in Arabidopsis, PLoS One. 4 (2009). https://doi.org/10.1371/journal.pone.0007817.

[94] J.Y. Yun, Y. Tamada, Y.E. Kang, R.M. Amasino, Arabidopsis trithorax-related3/set domain group2 is required for the winter-annual habit of arabidopsis thaliana, Plant Cell Physiol. 53 (2012) 834–846. https://doi.org/10.1093/pcp/pcs021.

[95] S. Napsucialy-Mendivil, R. Alvarez-Venegas, S. Shishkova, J.G. Dubrovsky, Arabidopsis homolog of trithorax1 (ATX1) is required for cell production, patterning, and morphogenesis in root development, J. Exp. Bot. 65 (2014) 6373–6384. https://doi.org/10.1093/jxb/eru355.

[96] A. Saleh, R. Alvarez-Venegas, M. Yilmaz, O. Le, G. Hou, M. Sadder, A. Al-Abdallat, Y. Xia, G. Lu, I. Ladunga, Z. Avramova, The Highly Similar Arabidopsis Homologs of Trithorax ATX1 and ATX2 Encode Proteins with Divergent Biochemical Functions, Plant Cell Online. 20 (2008) 568–579. https://doi.org/10.1105/tpc.107.056614.

[97] X. Yao, H. Feng, Y. Yu, A. Dong, W.H. Shen, SDG2-Mediated H3K4 Methylation Is Required for Proper Arabidopsis Root Growth and Development, PLoS One. 8 (2013) 1–11. https://doi.org/10.1371/journal.pone.0056537.

[98] K. Lee, O.S. Park, P.J. Seo, ATXR2 as a core regulator of de novo root organogenesis, Plant Signal. Behav. 13 (2018) 1–2. https://doi.org/10.1080/15592324.2018.1449543.

[99] S. Qian, Y. Wang, H. Ma, L. Zhang, Expansion and Functional Divergence of Jumonji C-Containing Histone Demethylases: Significance of Duplications in Ancestral Angiosperms and Vertebrates, Plant Physiol. 168 (2015) 1321–1337. https://doi.org/10.1104/pp.15.00520.

[100] E.S. Gan, Y. Xu, J.Y. Wong, J. Geraldine Goh, B. Sun, W.Y. Wee, J. Huang, T. Ito, Jumonji demethylases moderate precocious flowering at elevated temperature via regulation of FLC in Arabidopsis, Nat. Commun. 5 (2014) 1–13. https://doi.org/10.1038/ncomms6098.

[101] X. Liu, W. Wei, W. Zhu, L. Su, Z. Xiong, M. Zhou, Y. Zheng, D.X. Zhou, Histone Deacetylase AtSRT1 Links Metabolic Flux and Stress Response in Arabidopsis, Mol. Plant. 10 (2017) 1510–1522. https://doi.org/10.1016/j.molp.2017.10.010.

[102] L. Huang, Q. Sun, F. Qin, C. Li, Y. Zhao, D.X. Zhou, Down-regulation of a Silent Information Regulator2-related histone deacetylase gene, OsSRT1, induces DNA fragmentation and cell death in rice, Plant Physiol. 144 (2007) 1508–1519. https://doi.org/10.1104/pp.107.099473.

[103] A. Zemach, M. Kim, P. Hsieh, D. Coleman-Derr, L. Williams-Eshed, The nucleosome remodeler DDM1 allows DNA methyltransferases to access H1-containing heterochromatin, Cell. 153 (2013) 193–205. https://doi.org/10.1016/j.cell.2013.02.033.The.

[104] H. Stroud, T. Do, J. Du, X. Zhong, S. Feng, D.J. Patel, S.E. Jacobsen, The roles of non-CG methylation in Arabidopsis, Nat Struct Mol Biol. 21 (2014) 64–72. https://doi.org/10.1038/nsmb.2735.

[105] M. Hartl, M. Füβl, P.J. Boersema, J. Jost, K. Kramer, A. Bakirbas, J. Sindlinger, M. Plöchinger, D. Leister, G. Uhrig, G.B. Moorhead, J. Cox, M.E. Salvucci, D. Schwarzer, M. Mann, I. Finkemeier, Lysine acetylome profiling uncovers novel histone deacetylase substrate proteins in Arabidopsis, Mol. Syst. Biol. 13 (2017) 949. https://doi.org/10.15252/msb.20177819.

[106] M.L. Ebbs, Locus-Specific Control of DNA Methylation by the Arabidopsis SUVH5 Histone Methyltransferase, Plant Cell Online. 18 (2006) 1166–1176. https://doi.org/10.1105/tpc.106.041400.

