## Supplemental figures and tables for "Genome-Wide Identification of Epigenetic Regulators in *Quercus suber*"

| **2.8. Supplementary data**  **Supplementary File S1. List of all the DNA and histone epigenetic modifiers used in this study for the phylogenetic tree construction.** The protein name used in this study alongside with the annotation ID is represented. | | | |
| --- | --- | --- | --- |
| **Given name** | **Public Database ID** | **Given name** | **Public Database ID** |
| *VvCMT3* | XP_010651344.1 | *AtDRM3* | AT3G17310 |
| *CmCMT3* | XP_008451103.1 | *AtDRM2* | AT5G14620 |
| *GmCMT3* | XP_006572936.1 | *AtDRM1* | AT5G15380 |
| *JgCMT3* | XP_018834603.1 | *QrDRM2A* | Loc_1382_Tr_6/6_Conf_0.667_Len_2740 |
| *QrCMT3* | Loc_5202_Tr_1/2_Conf_0.889_Len_3208 | *QrDRM2B* | Loc_82954_Tr_2/2_Conf_0.750_Len_1158 |
| *PmCMT3* | XP_008226363.1 | *CmoDRM3* | maker-scaffold01664-augustus-gene-0.28 |
| *PpCMT3* | XP_020421159.1 | *CmoDRM2* | maker-scaffold03040-augustus-gene-0.18 |
| *MdCMT3* | NP_001280801.1 | *QrDRM3* | OCV4_rep_c18566 |
| *JcCMT3* | XP_012093320.1 | *VvDRM3* | XP_002264226.1 |
| *BrCMT3* | XP_009150572.1 | *PtDRM2* | XP_002300046.2 |
| *AtCMT3* | AT1G69770 | *PtDRM3* | XP_002316067.2 |
| *VvCMT1* | XP_002275932.1 | *GmDRM2* | XP_006583974.1 |
| *JcCMT1* | XP_012076052.1 | *PpDRM2* | XP_007201047.2 |
| *QrCMT1* | Loc_21057_Tr_1/5_Conf_0.667_Len_2822 | *PmDRM1* | XP_008223451.1 |
| *CmoCMT1* | maker-scaffold12039-augustus-gene-0.8 | *MdDRM2* | XP_008379233.1 |
| *GmCMT1a* | XP_006573534.1 | *CmDRM2* | XP_008458306.1 |
| *GmCMT1b* | XP_006590741.1 | *BrDRM1* | XP_009105496.1 |
| *BrCMT1* | XP_009148471.2 | *BrDRM2a* | XP_009121651.1 |
| *AtCMT1* | AT1G80740 | *BrDRM2b* | XP_009125988.1 |
| *GmCMT2* | XP_006599215.1 | *BrDRM3* | XP_009146023.1 |
| *PmCMT2* | XP_008238301.2 | *CsDRM2* | XP_011656312.1 |
| *PpCMT2* | ONI05788.1 | *JcDRM2* | XP_012072470.1 |
| *JrCMT2* | XP_018806585.1 | *JcDRM3* | XP_012084156.1 |
| *QrCMT2* | OCV4_rep_c4395 | *PmDRM2* | XP_016650960.1 |
| *CmCMT2* | XP_008448610.1 | *MdDRM1* | XP_017192312.1 |
| *CsCMT2* | XP_011650321.1 | *JgDRM2* | XP_018823107.1 |
| *JcCMT2* | XP_012066500.1 | *JgDRM1* | XP_018833888.1 |
| *VvCMT2* | XP_010665031.1 | *PpDRM3* | XP_020410202.1 |
| *BrCMT2* | XP_009132470.1 | *OsDRM1-2a* | LOC_Os03g02010.4 |
| *AtCMT2* | AT4G19020 | *OsDRM1-2b* | LOC_Os11g01810.1 |
| *OsCMT1-3b* | LOC_Os03g12570.1 | *OsDRM3* | LOC_Os05g04330.1 |
| *OsCMT1-3c* | LOC_Os10g01570.1 | *OsCMT1-3a* | LOC_Os05g13780.1 |
| *PpMET1* | AAM96952.1 | *VvDNMT2* | XP_002282454.2 |
| *PmMET1* | XP_008241936.1 | *JrDNMT2* | XP_018834793.1 |
| *MdMET1* | XP_008361333.1 | *QrDNMT2* | OCV4_rep_c6323 |
| *VvMET1a* | XP_002267200.1 | *CmoDNMT2* | maker-scaffold00063-snap-gene-0.34 |
| *VvMET1b* | XP_002267284.3 | *JcDNMT2* | XP_012076609.1 |
| *JgMET1* | XP_018821418.1 | *PpDNMT2* | XP_007223053.1 |
| *QrMET1* | OCV4_c25957 | *MdDNMT2* | XP_008394053.1 |
| *CmoMET1* | maker-scaffold03820-snap-gene-0.12 | *BrDNMT2* | XP_009151075.1 |
| *PtMET1* | XP_002305346.1 | *AtDNMT2* | AT5G25480 |
| *JcMET1* | XP_012088404.1 | *GmDNMT2* | NP_001242866.1 |
| *GmMET1a* | XP_014630288.1 | *CsDNMT2* | XP_004138735.1 |
| *GmMET1b* | XP_014632022.1 | *CmDNMT2* | XP_008445162.1 |
| *BrMET1a* | BAF34636.1 | *OsDNMT2* | LOC_Os01g42630.2 |
| *BrMET1b* | XP_009117317.1 | *AtMET3* | AT4G13610 |
| *AtMET1* | AT5G49160 | *OsMET1-3a* | LOC_Os03g58400.1 |
| *AtMET2a* | AT4G14140 | *OsMET1-3b* | LOC_Os07g08500.1 |
| *AtMET2b* | AT4G08990 | *CmoROS1* | augustus masked-scaffold02965-abinit-gene-0.4-mRNA-1 |
| *PtDME1.1* | XP_002316518.2 | *QrROS1* | OCV4_rep_c685 |
| *PtDME1.2* | XP_002311047.3 | *OsDME* | LOC_Os01g11900.1 |
| *JcDME* | XP_012089968.1 | *OsROS1.1* | LOC_Os05g37410.1 |
| *MdDME* | XP_028964401.1 | *OsROS1.2* | LOC Os05g37350.1 |
| *PmDME* | XP_008240460.1 | *AtROS1* | AT2G36490 |
| *PpDME* | XP_020423437.1 | *BrROS1.1* | XP_009143580.1 |
| *AtDME* | AT5G045609 | *BrROS1.2* | RID61705.1 |
| *JrDME1.1* | XP_018853480.1 | *AtDML2* | AT3G10010 |
| *JrDME1.2* | XP_018845375.1 | *CmoDML2* | maker-scaffold06611-augustus-gene-0.9-mRNA-1 |
| *CmoDME* | augustus masked-scaffold00254-abinit-gene-0.14-mRNA-1 | *AtDML3* | AT4G34060 |
| *QrDME1.1* | OCV4 _rep_c21323 | *BrDML3.1* | XP_009117384.1 |
| *QrDME1.2* | OCV4_rep_c3947 | *BrDML3.2* | RID77506.1 |
| *VvROS1* | CBI30244.3 | *GmROS1* | XP_006588821.1 |
| *JcROS1* | XP_012066501.1 | *QsROS1* | XP_023928322.1 |
| *MdHAC1* | XP_008380702.1 | *QrHAM1* | OCV3_prime_c11529 |
| *PmHAC1* | XP_008245061.1 | *CsHAM1* | XP_004140319.1 |
| *JrHAC1a* | XP_018859433.1 | *CmHAM1* | XP_008460476.1 |
| *JrHAC1b* | XP_018822549.1 | *CmoHAM1* | maker-scaffold04599-snap-gene-0.15 |
| *CmoHAC1* | maker-scaffold02425-snap-gene-0.24 | *PtHAM1a* | XP_002301018.1 |
| *QrHAC1* | OCV4_rep_c42029 | *PtHAM1b* | XP_002307411.1 |
| *JcHAC1* | XP_012087277.1 | *JcHAM1* | XP_012078147.1 |
| *VvHAC1* | XP_010655213.1 | *JrHAM1a* | XP_018849623.1 |
| *GmHAC1* | XP_006582962.1 | *JrHAM1* | XP_018850689.1 |
| *CsHAC1* | XP_011650235.1 | *VvHAM2* | XP_002285829.1 |
| *CmHAC1* | XP_008448424.1 | *MdHAM2* | XP_008338814.1 |
| *BrHAC1a* | XP_009128325.1 | *PmHAM2* | XP_008233928.1 |
| *BrHAC1b* | XP_009106616.1 | *PpHAM2* | XP_007222151.1 |
| *AtHAC1* | AT1G79000 | *GmHAM1* | XP_003522879.1 |
| *AtHAC12* | AT1G16710 | *BrHAM1* | XP_009112143.1 |
| *BrHAC5a* | XP_009135269.1 | *AtHAM1* | AT5G64610 |
| *BrHAC5b* | XP_009124693.1 | *BrHAM2* | XP_009122536.1 |
| *AtHAC5* | AT3G12980 | *AtHAM2* | AT5G09740 |
| *AtHAC4* | AT1G55970 | *OsHAM1* | LOC_Os07g43360.1 |
| *OsHAC1A* | LOC_Os06g49130.2 | *QsHAF1* | XP_023879413.1 |
| *OsHAC1b* | LOC_Os02g04490.1 | *MdHAF1* | XP_008369193.1 |
| *CmoHAC1-like1* | maker-scaffold04214-snap-gene-0.12 | *PmHAF1* | XP_008218889.1 |
| *CmoHAC1-like2* | maker-scaffold00359-snap-gene-1.12 | *PpHAF1* | XP_020410681.1 |
| *QrHAC1-like2* | Loc_39724_Tr_1/2_Conf_0.750_Len_4836 | *JrHAF1* | XP_018846793.1 |
| *BrHAC2b* | XP_009127535.1 | *CmoHAF1* | maker-scaffold00376-augustus-gene-0.24 |
| *BrHAC2a* | XP_009105610.1 | *QrHAF1* | OCV4_rep_c6162 |
| *AtHAC2* | AT1G67220 | *VvHAF1* | XP_010656962.1 |
| *OsHAC5* | LOC_Os01g14370.1 | *GmHAF1a* | XP_006578382.1 |
| *MdHAG2a* | XP_008387482.1 | *GmHAF1b* | XP_006587642.1 |
| *MdHAG2b* | XP_008337633.1 | *PtHAF1* | XP_002323740.2 |
| *PpHAG2* | XP_020423543.1 | *JcHAF1* | XP_020534041.1 |
| *JrHAG2* | XP_018846279.1 | *CmHAF1* | XP_008458065.1 |
| *CmoHAG2* | augustus_masked-scaffold09784-abinit-gene-0.0 | *BrHAF2* | XP_009112153.1 |
| *QrHAG2* | OCV4_rep_c26572 | *BrHAF1* | XP_009107984.1 |
| *PtHAG2* | XP_002319203.2 | *AtHAF1* | AT1G32750 |
| *JcHAG2* | XP_012076804.1 | *AtHAF2* | AT3G19040.1 |
| *VvHAG2* | XP_002282931.1 | *OsHAF1* | LOC_Os06g43790.1 |
| *GmHAG2a* | XP_003519360.1 | *JrELP3* | XP_018814787.1 |
| *GmHAG2b* | XP_003544183.1 | *CsELP3* | XP_004138068.1 |
| *CsHAG2* | XP_004141044.1 | *CmELP3* | XP_008464479.1 |
| *CmHAG2* | XP_008459220.1 | *PmELP3* | XP_008239412.1 |
| *BrHAG2a* | XP_009132185.1 | *PpELP3* | XP_007210504.1 |
| *BrHAG2b* | XP_009120185.1 | *GmELP3a* | XP_003523129.1 |
| *BrHAG2c* | XP_018510752.1 | *GmELP3b* | XP_003526977.1 |
| *AtHAG2* | AT5G56740 | *CmoELP3a* | maker-scaffold01033-snap-gene-0.30 |
| *OsHAG2* | LOC_Os09g17850.1 | *QrELP3* | Loc_4308_Tr_2/4_Conf_0.600_Len_2295 |
| *PtELP3* | XP_002318107.2 | *VvELP3* | XP_002262701.1 |
| *JcELP3* | XP_012079906.1 | *BrELP3* | XP_009134057.1 |
| *PmGCN5* | XP_008235336.1 | *AtELP3* | AT5G50320 |
| *PpGCN5* | XP_007201717.1 | *OsELP3* | LOC_Os04g40840.1 |
| *JrGCN5* | XP_018844562.1 | *CmoELP3b* | augustus_masked-scaffold36664-abinit-gene-0.0 |
| *QrGCN5* | OCV4_c7575 | *PtGCN5b* | XP_002306812.1 |
| *CmoGCN5* | maker-scaffold06069-snap-gene-0.16 | *PtGCN5a* | XP_006386247.1 |
| *GmGCN5a* | XP_003520580.1 | *JcGCN5* | XP_012092127.1 |
| *GmGCN5b* | XP_003553477.1 | *BrGCN5b* | XP_009139225.1 |
| *CsGCN5* | XP_004134529.1 | *BrGCN5a* | XP_009139228.1 |
| *VvGCN5* | XP_002275146.2 | *AtGCN5* | AT3G54610 |
| *OsGCN5* | LOC_Os10g28040.1 | *VvASHH1* | XP_019072374.1 |
| *PtSWN* | XP_002320296.1 | *GmASHH1a* | XP_003536414.1 |
| *PtSWN* | PNT50623.1 | *GmASHH1b* | XP_003556159.2 |
| *JcSWN* | XP_012082901.1 | *PtASHH1* | XP_002306713.2 |
| *JrSWN* | XP_018842484.1 | *PmASHH1* | XP_008237500.1 |
| *CmoSWN* | maker-scaffold05636-augustus-gene-0.11 | *PpASHH1* | XP_007199600.2 |
| *QrSWN* | Loc_1192_Tr_4/8_Conf_0.696_Len_3240 | *MdASHH1* | XP_008372737.1 |
| *GmSWNa* | XP_003519745.1 | *JrASHH1* | XP_018848121.1 |
| *GmSWNb* | XP_006588564.1 | *CmoASHH1* | maker-scaffold03331-augustus-gene-0.18 |
| *CsSWN* | XP_004152222.1 | *QrASHH1* | Loc_12859_Tr_1/4_Conf_0.769_Len_2491 |
| *CmSWN* | XP_008454285.1 | *JcASHH1* | XP_012075212.1 |
| *BrSWN* | XP_009111326.1 | *CmASHH1* | XP_008441493.1 |
| *AtSWN* | AT4G02020.1 | *CsASHH1* | XP_011656629.1 |
| *VvCLF* | CBI21398.3 | *BrASHH1* | XP_009106340.1 |
| *PtCLFa* | XP_002307273.2 | *GmASHH1* | AT1G76710.2 |
| *PtCLFb* | XP_002310129.2 | *GmASHH3* | XP_006587390.1 |
| *JcCLF* | XP_012079259.1 | *JrASHH3* | XP_018839332.1 |
| *PmCLF* | XP_008241485.1 | *CmoASHH3* | maker-scaffold00111-snap-gene-0.32 |
| *PpCLF* | XP_020424806.1 | *QrASHH3* | OCV4_rep_c40652 |
| *MdCLF* | XP_008340296.1 | *PpASHH3* | XP_007202086.1 |
| *QrCLF* | OCV4_rep_c16251 | *PmASHH3* | XP_016651961.1 |
| *CmCLF* | XP_008457756.1 | *MdASHH3* | XP_008394239.1 |
| *CsCLF* | XP_011649325.1 | *CsASHH3* | XP_011649104.1 |
| *BrCLF* | XP_009140462.1 | *CmASHH3* | XP_016902074.1 |
| *AtCLF* | AT2G23380.1 | *PtASHH3a* | XP_002310367.2 |
| *AtMEA* | AT1G02580.1 | *PtASHH3b* | PNS94539.1 |
| *OsCLF* | LOC_Os06g16390.1 | *JcASHH3* | XP_012081879.1 |
| *OsMEA\|SWN* | LOC_Os03g19480.1 | *CsASHR3* | XP_004146715.1 |
| *VvATXR7* | XP_019074011.1 | *CmASHR3* | XP_008443822.1 |
| *PtATXR7a* | PNT47006.1 | *PpASHR3* | XP_020410652.1 |
| *PtATXR7b* | PNT38856.1 | *PmASHR3* | XP_016647450.1 |
| *JcATXR7* | XP_012073523.1 | *PtASHR3* | XP_002325110.1 |
| *MdATXR7* | XP_008347884.1 | *JcASHR3* | XP_012071580.1 |
| *JrATXR7* | XP_018843102.1 | *CmoASHR3* | maker-scaffold06327-augustus-gene-0.9 |
| *CmoATXR7* | maker-scaffold01202-augustus-gene-0.28 | *QrASHR3* | OCV4_rep_c19155 |
| *QrATXR7* | Loc_9902_Tr_7/8_Conf_0.682_Len_7488 | *VvASHR3* | XP_002272781.1 |
| *BrATXR7* | XP_018510432.1 | *BrASHR3* | XP_018508962.1 |
| *AtATXR7* | AT5G42400.1 | *AtASHR3* | AT4G30860.1 |
| *VvATXR7* | XP_010661927.1 | *BrASHH3a* | XP_009133661.1 |
| *GmATX3a* | XP_006583237.1 | *BrASHH3b* | XP_009142998.1 |
| *GmATX3b* | XP_006598904.1 | *AtASHH3* | AT2G44150.1 |
| *PmATX3* | XP_008233180.2 | *JrASHR3* | XP_018818672.1 |
| *PpATX3* | XP_020411934.1 | *BrASHH4a* | XP_009104232.1 |
| *JrATX3a* | XP_018846718.1 | *BrASHH4b* | XP_018510238.1 |
| *JrATX3b* | XP_018846720.1 | *AtASHH4* | AT3G59960.1 |
| *QrATX3* | Loc_20589_Tr_3/6_Conf_0.688_Len_3877 | *JcASHH2* | XP_012078323.1 |
| *PtATX3a* | XP_002302628.2 | *JrASHH2a* | XP_018849175.1 |
| *PtATX3b* | XP_002320864.2 | *JrASHH2b* | XP_018811342.1 |
| *JcATX3* | XP_012083217.1 | *QrASHH2* | OCV4_rep_c21713 |
| *CmATX3* | XP_008459998.1 | *PmASHH2* | XP_008219574.1 |
| *CsATX3a* | KGN46349.1 | *PpASHH2* | XP_020411481.1 |
| *CsATX3b* | XP_011656748.1 | *MdASHH2* | XP_008378450.1 |
| *GmATX4a* | XP_003527980.1 | *VvASHH2* | XP_010664163.1 |
| *GmATX4b* | XP_006578910.1 | *BrASHH2* | XP_018508679.1 |
| *JcATX4* | XP_012090074.1 | *AtASHH2* | AT1G77300.1 |
| *PtATX5a* | XP_002318412.2 | *OsASHH1* | LOC_Os04g34976.2 |
| *PtATX5b* | XP_002321418.2 | *OsASHH2* | LOC_Os02g34850.1 |
| *PmATX5* | XP_008234739.1 | *OsASHH3-4* | LOC_Os09g13740.1 |
| *PpATX5* | XP_020413261.1 | *OsASHR3* | LOC_Os02g39800.1 |
| *MdATX4* | XP_008380864.1 | *VvATXR7* | XP_019074011.1 |
| *MdATX4-like* | XP_008347258.1 | *PtATXR7a* | PNT47006.1 |
| *JrATX5* | XP_018838727.1 | *PtATXR7b* | PNT38856.1 |
| *QrATX5* | OCV4_c26314 | *JcATXR7* | XP_012073523.1 |
| *JrATX4* | XP_018850756.1 | *MdATXR7* | XP_008347884.1 |
| *CmATX4* | XP_008466814.1 | *JrATXR7* | XP_018843102.1 |
| *CsATX4* | XP_011651328.1 | *CmoATXR7* | maker-scaffold01202-augustus-gene-0.28 |
| *BrATX4* | XP_018508983.1 | *QrATXR7* | Loc_9902_Tr_7/8_Conf_0.682_Len_7488 |
| *AtATX4* | AT4G27910.1 | *BrATXR7* | XP_018510432.1 |
| *BrATX5* | XP_009119888.1 | *AtATXR7* | AT5G42400.1 |
| *AtATX5* | AT5G53430.1 | *VvATXR7* | XP_010661927.1 |
| *BrATX3* | XP_009104342.1 | *GmATX3a* | XP_006583237.1 |
| *AtATX3* | AT3G61740.1 | *GmATX3b* | XP_006598904.1 |
| *VvATX1* | XP_002268621.1 | *PmATX3* | XP_008233180.2 |
| *PtATX1a* | XP_002301643.2 | *PpATX3* | XP_020411934.1 |
| *PtATX1b* | XP_002320433.2 | *JrATX3a* | XP_018846718.1 |
| *JcATX2* | XP_020533084.1 | *JrATX3b* | XP_018846720.1 |
| *JrATX2* | XP_018815781.1 | *QrATX3* | Loc_20589_Tr_3/6_Conf_0.688_Len_3877 |
| *CmoATX2* | maker-scaffold01217-snap-gene-0.28 | *PtATX3a* | XP_002302628.2 |
| *QrATX2* | OCV4_rep_c43523 | *PtATX3b* | XP_002320864.2 |
| *PpATX2* | XP_007225413.1 | *JcATX3* | XP_012083217.1 |
| *PmATX2* | XP_008244420.1 | *CmATX3* | XP_008459998.1 |
| *MdATX2* | XP_008337536.1 | *CsATX3a* | KGN46349.1 |
| *CmATX2* | XP_008464329.1 | *CsATX3b* | XP_011656748.1 |
| *CsATX2* | XP_011656479.1 | *GmATX4a* | XP_003527980.1 |
| *BrATX1* | XP_009141151.1 | *GmATX4b* | XP_006578910.1 |
| *AtATX1* | AT2G31650.1 | *JcATX4* | XP_012090074.1 |
| *BrATX2* | XP_009118571.1 | *PtATX5a* | XP_002318412.2 |
| *AtATX2* | AT1G05830. | *PtATX5b* | XP_002321418.2 |
| *OsATX1* | LOC_Os09g04890.1 | *PmATX5* | XP_008234739.1 |
| *OsATX2-5* | LOC_Os01g11952.1 | *PpATX5* | XP_020413261.1 |
| *OsATX4-5* | LOC_Os01g46700.1 | *MdATX4* | XP_008380864.1 |
| *OSATXR7* | LOC_Os12g41900.1 | *MdATX4-like* | XP_008347258.1 |
| *OsATXR3* | LOC_Os08g08210.1 | *JrATX5* | XP_018838727.1 |
| *PtATXR3* | XP_006372997.1 | *QrATX5* | OCV4_c26314 |
| *JcATXR3* | XP_012084659.1 | *JrATX4* | XP_018850756.1 |
| *PmATXR3* | XP_008230126.1 | *CmATX4* | XP_008466814.1 |
| *PpATXR3* | XP_020416287.1 | *CsATX4* | XP_011651328.1 |
| *MdATXR3* | XP_008341967.2 | *BrATX4* | XP_018508983.1 |
| *JrATXR3* | XP_018846411.1 | *AtATX4* | AT4G27910.1 |
| *CmoATXR3* | maker-scaffold02899-snap-gene-0.16 | *BrATX5* | XP_009119888.1 |
| *QrATXR3* | OCV4_rep_c4016 | *AtATX5* | AT5G53430.1 |
| *JrATXR3* | XP_018807833.1 | *BrATX3* | XP_009104342.1 |
| *VvATXR3* | XP_010657340.1 | *AtATX3* | AT3G61740.1 |
| *GmATXR3a* | XP_006582339.1 | *VvATX1* | XP_002268621.1 |
| *GmATXR3b* | XP_006592400.1 | *PtATX1a* | XP_002301643.2 |
| *GmATXR3c* | XP_006592825.1 | *PtATX1b* | XP_002320433.2 |
| *GmATXR3d* | XP_006594874.1 | *JcATX2* | XP_020533084.1 |
| *CmATXR3* | XP_008455393.1 | *JrATX2* | XP_018815781.1 |
| *CsATXR3* | XP_011658717.1 | *CmoATX2* | maker-scaffold01217-snap-gene-0.28 |
| *BrATXR3a* | XP_009147501.1 | *QrATX2* | OCV4_rep_c43523 |
| *BrATXR3b* | XP_009144717.1 | *PpATX2* | XP_007225413.1 |
| *AtATXR3* | AT4G15180.1 | *PmATX2* | XP_008244420.1 |
| *VvSUVR3* | XP_002277066.1 | *MdATX2* | XP_008337536.1 |
| *PtSUVR3* | XP_002319719. | *CmATX2* | XP_008464329.1 |
| *PmSUVR3* | XP_008244134.1 | *CsATX2* | XP_011656479.1 |
| *PpSUVR3* | XP_007203658.2 | *BrATX1* | XP_009141151.1 |
| *JrSUVR3* | XP_018807125.1 | *AtATX1* | AT2G31650.1 |
| *CmoSUVR3* | maker-scaffold23019-augustus-gene-0.5 | *BrATX2* | XP_009118571.1 |
| *QrSUVR3* | OCV4_rep_c24922 | *AtATX2* | AT1G05830. |
| *GmSUVR3* | XP_003555385.2 | *OsATX1* | LOC_Os09g04890.1 |
| *JcSUVR3* | XP_012070535.1 | *OsATX2-5* | LOC_Os01g11952.1 |
| *BrSUVR3* | XP_009130385.1 | *OsATX4-5* | LOC_Os01g46700.1 |
| *AtSUVR3* | AT3G03750.2 | *OSATXR7* | LOC_Os12g41900.1 |
| *CsSUVR3* | XP_004136662.1 | *OsATXR3* | LOC_Os08g08210.1 |
| *CmSUVR3* | XP_008443304.1 | *PtATXR3* | XP_006372997.1 |
| *VvSUVH8* | XP_010646790.1 | *JcATXR3* | XP_012084659.1 |
| *PmSUVH5* | XP_008238503.1 | *PmATXR3* | XP_008230126.1 |
| *PpSUVH5* | XP_020420071.1 | *PpATXR3* | XP_020416287.1 |
| *MdSUVH6* | XP_008386441.2 | *MdATXR3* | XP_008341967.2 |
| *JrSUVH6* | XP_018821426.1 | *JrATXR3* | XP_018846411.1 |
| *CmoSUVH6b* | snap_masked-scaffold01285-abinit-gene-0.28 | *CmoATXR3* | maker-scaffold02899-snap-gene-0.16 |
| *QsSUVH6* | OCV4_c18 | *QrATXR3* | OCV4_rep_c4016 |
| *CmoSUVH6a* | maker-scaffold04831-augustus-gene-0.25 | *JrATXR3* | XP_018807833.1 |
| *CmSUVH6* | XP_008448779.1 | *VvATXR3* | XP_010657340.1 |
| *CsSUVH6* | XP_011650376.1 | *GmATXR3a* | XP_006582339.1 |
| *PtSUVH5* | XP_006385561.1 | *GmATXR3b* | XP_006592400.1 |
| *JcSUVH6* | XP_012070094.2 | *GmATXR3c* | XP_006592825.1 |
| *BrSUVH6* | XP_009140380.1 | *GmATXR3d* | XP_006594874.1 |
| *AtSUVH6* | AT2G22740.2 | *CmATXR3* | XP_008455393.1 |
| *BrSUVH5* | XP_009143694.1 | *CsATXR3* | XP_011658717.1 |
| *AtSUVH5* | AT2G35160.1 | *BrATXR3a* | XP_009147501.1 |
| *VvSUVH4* | XP_010660678.1 | *BrATXR3b* | XP_009144717.1 |
| *JcSUVH4* | XP_012067355.1 | *AtATXR3* | AT4G15180.1 |
| *JrSUVH4* | XP_018834874.1 | *GmSET40* | XP_006596494.1 |
| *CmoSUVH4* | maker-scaffold07377-snap-gene-0.17 | *VvSET40* | XP_002269094.3 |
| *QrSUVH4* | OCV4_rep_c40958 | *PpSET40* | XP_007215291.1 |
| *PtSUVH4* | PNT04845.1 | *PmSET40* | XP_008227893.1 |
| *PmSUVH4* | XP_008231237.1 | *MdSET40* | XP_017179820.1 |
| *PpSUVH4* | XP_020415335.1 | *PtSET40* | PNT40156.1 |
| *MdSUVH4* | XP_008379740.1 | *JcSET40* | XP_012074708.1 |
| *CmSUVH4* | XP_008449992.1 | *CsSET40* | XP_004145844.1 |
| *CsSUVH4* | XP_011651591.1 | *CmSET40* | XP_008457029.1 |
| *GmSUVH4a* | XP_003548493.1 | *JrSET40* | XP_018846347. |
| *GmSUVH4b* | XP_006604199.1 | *CmoSET40* | maker-scaffold05691-snap-gene-0.16 |
| *BrSUVH4* | XP_009131370.1 | *QrSET40* | Loc_19922_Tr_2/5_Conf_0.636_Len_1707 |
| *AtSUVH4* | AT5G13960.1 | *BrSET40* | XP_009126173.1 |
| *QrSUVH4-like* | Loc_19624_Tr_8/9_Conf_0.609_Len_2652 | *AtSET40* | AT5G17240.1 |
| *VvSUVH1b* | XP_002278728.1 | *OsSET40* | LOC_Os07g28840.1 |
| *CmoSUVH1* | snap_masked-scaffold01698-abinit-gene-0.19 | *QrSET10* | Loc_11855_Tr_5/6_Conf_0.611_Len_2331 |
| *QrSUVH1* | OCV4_c19481 | *OsSET10* | LOC_Os06g03676.1 |
| *MdSUVH1* | XP_008387456.1 | *CmoSET10* | maker-scaffold04661-snap-gene-0.29 |
| *GmSUVH1a* | XP_006576757.1 | *AtSET10* | At1g01920 |
| *GmSUVH1b* | XP_014627153.1 | *BrSET10* | VDC64657.1 |
| *PtSUVH1a* | XP_002303967.1 | *JrSET10* | XP_018805358.1 |
| *PtSUVH1b* | XP_002299167.2 | *PpSET10* | XP_007217928.2 |
| *JcSUVH3* | XP_012068760.1 | *PmSET10* | XP_008232866.1 |
| *VvSUVH1a* | XP_002273935.1 | *PtSET10* | XP_024440252.1 |
| *JrSUVH1a* | XP_018818592.1 | *JcSET10* | XP_020538444.1 |
| *JrSUVH1b* | XP_018806577.1 | *GmSET10* | XP_006602773.1 |
| *CmoSUVH3* | augustus_masked-scaffold00709-abinit-gene-0.2 | *CmoSET10* | maker-scaffold07375-augustus-gene-0.17 |
| *QrSUVH3* | OCV4_rep_c21535 | *QrSET41* | OCV4_c22093 |
| *PpSUVH1* | XP_007204245.1 | *OsSET41* | LOC_Os04g34610.1 |
| *PmSUVH3* | XP_008242105.1 | *AtSET41* | AT1G43245.1 |
| *PpSUVH1b* | XP_007220227.1 | *BrSET41* | XP_009111402.1 |
| *PpSUVH1a* | ONI21243.1 | *CmSET41* | XP_008463080.1 |
| *PmSUVH1* | XP_008245334.1 | *CsSET41* | XP_011656459.1 |
| *MdSUVH1a* | XP_008346233.1 | *JcSET41* | XP_012080731.1 |
| *MdSUVH1b* | XP_008375400.1 | *JrSET41* | XP_018844201.1 |
| *BrSUVH1c* | XP_009125582.1 | *PtSET41* | XP_024457357.1 |
| *BrSUVH1a* | XP_009130849.1 | *GmSET41* | XP_006599489.1 |
| *BrSUVH1b* | XP_009122547.1 | *PpSET41* | XP_020425128.1 |
| *AtSUVH1* | AT5G04940.2 | *VvSET41* | XP_010665141.1 |
| *BrSUVH1d* | XP_009122548.1 | *CmSET41* | XP_008463081.1 |
| *BrSUVH3* | XP_009105965.1 | *VvASHR2* | XP_010657698.1 |
| *AtSUVH3* | AT1G73100.1 | *PtASHR2* | XP_002299366.2 |
| *BrSUVH7b* | XP_009149232.1 | *JcASHR2* | XP_012088668.1 |
| *BrSUVH7c* | XP_009149324.1 | *PpASHR2* | XP_007222652.1 |
| *BrSUVH7a* | XP_009105625.1 | *PmASHR2* | XP_008222297.1 |
| *AtSUVH7* | AT1G17770.1 | *MdASHR2a* | XP_008369584.1 |
| *AtSUVH8* | AT2G24740.1 | *MdASHR2b* | XP_008390264.1 |
| *VvSUVH9* | XP_002282386.1 | *GmASHR2* | XP_006585062.1 |
| *CsSUVH9* | XP_004134031.1 | *JrASHR2* | XP_018829239.1 |
| *CmSUVH9* | XP_008438443.1 | *CmoASHR2* | augustus_masked-scaffold05360-abinit-gene-0.3 |
| *JrSUVH9a* | XP_018807636.1 | *QrASHR2* | OCV4_rep_c11271 |
| *JrSUVH9b* | XP_018844439.1 | *BrASHR2* | XP_009112620.1 |
| *CmoSUVH9* | augustus_masked-scaffold02460-abinit-gene-0.3 | *AtASHR2* | AT2G19640.1 |
| *QrSUVH9* | Loc_7424_Tr_3/4_Conf_0.545_Len_5241 | *GmASHR2* | KRH76369.1 |
| *PpSUVH9* | XP_007226972.1 | *OsASHR2* | LOC_Os08g10470.1 |
| *PmSUVH9* | XP_008224025.1 | *VvASHR1* | XP_019072105.1 |
| *MdSUVH9a* | XP_008340678.1 | *JcASHR1* | XP_012078167.1 |
| *MdSUVH9b* | XP_008353179.1 | *JrASHR1* | XP_018814555.1 |
| *PtSUVH9a* | XP_002315593.2 | *CmoASHR1* | maker-scaffold01592-augustus-gene-0.38 |
| *PtSUVH9a* | PNT25203.1 | *QrASHR1* | OCV4_rep_c3041 |
| *JcSUVH9* | XP_012077634.1 | *PpASHR1* | XP_007222803.1 |
| *BrSUVH9* | XP_009116100.1 | *PmASHR1* | XP_008233908.1 |
| *AtSUVH9* | AT4G13460.1 | *MdASHR1* | XP_008338790.1 |
| *BrSUVH2* | XP_009141302.1 | *CmASHR1* | XP_008465815.1 |
| *AtSUVH2* | AT2G33290.1 | *CsASHR1* | XP_011655177.1 |
| *GmSUVR4* | XP_003520846.1 | *BrASHR1* | XP_009121063. |
| *QrSUVR4-like* | OCV4_c20491 | *AtASHR1* | AT2G17900.1 |
| *GmSYVR4a* | XP_014627854.1 | *OsASHR1* | LOC_Os03g49730.1 |
| *GmSUVR4b* | XP_014618709.1 | *VvASHR1* | XP_002274324.1 |
| *JcSUVR4* | KDP23728.1 | *JrASHHR1* | XP_018836331.1 |
| *PtSUVR4* | PNT06743.1 | *CmoASHR1* | maker-scaffold00558-snap-gene-0.38 |
| *VvSUVR4* | XP_010660173.1 | *QrASHR1* | OCV4_rep_c943 |
| *BrSUVR4a* | XP_009134761.1 | *PpATXR2* | XP_007215318.1 |
| *BrSUVR4b* | XP_009123996.1 | *PmATXR2* | XP_008230098.1 |
| *AtSUVR4* | AT3G04380 | *MdATXR2* | XP_008379613.1 |
| *QrSUVR4* | Loc_53227_Tr_1/1_Conf_1.000_Len_2710 | *CsATXR2* | XP_004151876.1 |
| *PtSUVR2a* | XP_002301851.2 | *CmATXR2* | XP_008455838.1 |
| *PtSUVR2b* | XP_002321292.2 | *PtATXR2* | XP_006372966.1 |
| *JcSUVR2* | XP_012085238.1 | *JcATXR2* | XP_012086379.1 |
| *MdSUVR2* | XP_008372385.1 | *BrATXR2* | XP_009145446.1 |
| *PpSUVR2* | XP_020425863.1 | *AtATXR2* | AT3G21820.1 |
| *JrSUVR2* | XP_018827288.1 | *GmATXR2* | XP_003540318.1 |
| *CmoSUVR2* | maker-scaffold04960-snap-gene-0.24 | *OsATXR2* | LOC_Os04g53700.1 |
| *QrSUVR2* | OCV4_rep_c16379 | *VvATXR4* | XP_002276611.1 |
| *BrSUVR2a* | XP_009101747.1 | *JrATXR4* | XP_018848850.1 |
| *BrSUVR2b* | XP_009123925.1 | *PtATXR4* | XP_002308346.2 |
| *AtSUVR2* | AT5G43990.1 | *JcATXR4* | XP_012090850.1 |
| *AtSUVR1* | AT1G04050.1 | *PmATXR4* | XP_008241157.1 |
| *GmSUVR5a* | XP_003516586.1 | *PpATXR4* | XP_007204308.2 |
| *GmSUVR5b* | XP_014619345.1 | *MdATXR4* | XP_008365789.1 |
| *VvSUVR5* | XP_010649212.1 | *CmoATXR4* | maker-scaffold14174-augustus-gene-0.4 |
| *PpSUVR5* | XP_007204800.1 | *QrATXR4* | Loc_30882_Tr_3/3_Conf_0.714_Len_832 |
| *PmSUVR5* | XP_008241605.1 | *BrATXR4* | XP_009125654.1 |
| *MdSUVR5a* | XP_008338988.1 | *AtATXR4* | AT5G06620.1 |
| *MdSUVR5b* | XP_008338178.1 | *OsATXR4* | LOC_Os10g27060.1 |
| *JrSUVR5a* | XP_018826472.1 | *VvATXR1* | XP_010655344.2 |
| *JrSUVR5b* | XP_018830205.1 | *JrATXR1* | XP_018850443.1 |
| *QrSUVR5* | Loc_7632_Tr_2/3_Conf_0.800_Len_7750 | *CmoATXR1* | augustus_masked-scaffold04895-abinit-gene-0.1 |
| *PtSUVR5a* | XP_002307228.2 | *QrATXR1* | Loc_33752_Tr_1/1_Conf_1.000_Len_2058 |
| *PtSUVR5b* | XP_006380742.1 | *PtATXR1* | XP_002312257.1 |
| *JcSUVR5a* | KDP31826.1 | *JcATXR1* | XP_012067684.1 |
| *JcSUVR5b* | XP_020537201.1 | *PmATXR1* | XP_008221586.1 |
| *BrSUVR5* | XP_009140488.1 | *PpATXR1* | XP_007222619.2 |
| *AtSUVR5* | AT2G23740.2 | *MdATXR1a* | XP_008389453.1 |
| *OsSUVH1\|SUVH3\|SUVH7-8* | LOC_Os11g38900.1 | *MdATXR1b* | XP_008339878.1 |
| *OsSUVH1* | LOC_Os01g59620.1 | *CsATXR1* | XP_004135447.1 |
| *OsSUVH1\|SUVH3* | LOC_Os05g41172.1 | *CmATXR1* | XP_008446386.1 |
| *OsSUVH2\|SUVH9* | LOC_Os07g25450.1 | *GmATXR1* | XP_003528581.1 |
| *OsSUVH4* | LOC_Os01g70220.1 | *BrATXR1* | XP_009115406.1 |
| *OsSUVH5-6* | LOC_Os09g19830.1 | *AtATXR1* | AT1G26760.1 |
| *OsSUVR1-2\|SUVR4* | LOC_Os02g40770.1 | *OsATXR1* | LOC_Os03g07260.1 |
| *OsSUVR3* | LOC_Os01g56540.1 | *OsSUVR5* | LOC_Os02g47900.1 |
| *AtLDL1* | AT1G62830.1 | *AtJMJ24* | AT1G09060.1 |
| *BrLDL1* | XP_009113042.1 | *BrJMJ24* | XP_009110857.1 |
| *JcLDL1* | XP_012085923.1 | *CmoJMJ24* | augustus_masked-scaffold04504 |
| *PtLDL1* | PNT47216.1 | *QrJMJ24* | OCV4_c26318 |
| *QrLDL1* | OCV4_rep_c28326 | *JrJMJ24* | XP_018837468. |
| *JrLDL1* | XP_018859441.1 | *PpJMJ24* | XP_020423075.1 |
| *VvLDL1* | XP_002265069.1 | *PmJMJ24* | XP_008246402.1 |
| *PpLDL1* | XP_007225269.1 | *VvJMJ24* | XP_002279731.2 |
| *PmLDL1* | XP_008221162.1 | *JcJMJ24* | XP_012089032.1 |
| *MdLDL1a* | XP_008377098.1 | *PtJMJ24a* | PNT06219.1 |
| *MdLDL1b* | XP_008387879.1 | *PtJMJ24b* | PNT34618.1 |
| *GmLDL1* | KRH39215.1 | *OsJMJ24* | LOC_Os03g22540.1 |
| *AtFLD* | AT3G10390.1 | *AtKDM3E* | AT1G11950.1 |
| *BrFLD* | XP_009135109.1 | *BrJMJ25-like3* | XP_009110682.1 |
| *OsFLD* | LOC_Os04g47270.1 | *AtKDM3F* | AT1G62310.1 |
| *CsFLD* | XP_011657505.1 | *BrJMJ25-like1* | XP_018510369.1 |
| *CmFLD* | XP_016902004.1 | *BrJMJ25-like2* | XP_009123379.1 |
| *CmoFLD* | maker-scaffold04812-snap-gene-0.17 | *CmoJMJ25-like1* | maker-scaffold01469-augustus-gene |
| *QrFLD* | OCV4_rep_c42703 | *QrJMJ25-like1* | OCV4_c27580 |
| *JrFLDa* | XP_018851616.1 | *JrJMJ25-like1* | XP_018844223.1 |
| *JrFLDb* | XP_018851624.1 | *MdJMJ25-like1* | XP_008371061.1 |
| *PpFLD* | XP_020413288.1 | *JcJMJ25-like1* | XP_020541213.1 |
| *PmFLD* | XP_008233274.1 | *AtJMJ25* | AT3G07610.1 |
| *MdFLD* | XP_008363874.1 | *BrJMJ25* | XP_009147050.1 |
| *VvFLD* | XP_010658356.1 | *VvJMJ25* | XP_010655917.1 |
| *PtFLDa* | PNT18181.1 | *CmoJMJ25-like2* | maker-scaffold03835-augustus-gene-0.8 |
| *PtFLDb* | PNT22486.1 | *PtJMJ25a* | PNT35579.1 |
| *AtLDL2* | AT3G13682.1 | *PtJMJ25b* | PNT35745.1 |
| *BrLDL2* | XP_009117559.1 | *AtKDM3D* | AT4G00990.1 |
| *JcLDL2a* | XP_012070642.1 | *BrKDM3D-like1* | XP_009111408.1 |
| *JcLDL2b* | XP_020534554.1 | *BrKDM3D-like2* | XP_009134497.1 |
| *PtLDL2* | XP_002316929.2 | *VvKDM3D-like* | CBI40561.3 |
| *CsLDL2* | XP_011652386.1 | *QrJMJ24-like1* | OCV4_rep_c9279 |
| *CmLDL2* | XP_008454649.1 | *CsKDM3D-like* | XP_011650678.1 |
| *CmoLDL2* | maker-scaffold10750-snap-gene-0.7 | *PpKDM3D-like* | XP_020412635.1 |
| *QrLDL2* | OCV4_c28695 | *PtKDM3D-like* | XP_002321404.2 |
| *JrLDL2* | XP_018823512.1 | *QrKDM3D-like1* | OCV4_rep_c15058 |
| *VvLDL2* | XP_002281860.3 | *QsKDM3E-F\|C-like1* | LOC_Os03g31594.1 |
| *PpLDL2* | XP_007213630.1 | *QsKDM3E-F\|C-like2* | LOC_Os02g58210.1 |
| *PmLDL2* | XP_008226688.1 | *AtKDM3B* | AT4G21430.1 |
| *MdLDL2* | XP_017186888.1 | *BrKDM3B-like1* | XP_009137189.1 |
| *GmLDL2* | XP_003527270.1 | *BtKDM3B-like2* | XP_009134735.1 |
| *OsLDL2* | LOC_Os08g04780.1 | *CsKDM3B-like* | XP_004134736.1 |
| *AtLDL3* | AT4G16310.1 | *CmKDM3B-like* | XP_008439919.1 |
| *BrLDL3* | XP_009124513.1 | *QrKDM3B-like* | Loc_11265_Tr_3/5_Conf_0.692_Len_5052 |
| *JcLDL3* | XP_020534824.1 | *CmoKDM3B-like* | maker-scaffold00476-augustus-gene-0.35 |
| *PtLDL3a* | PNT38359.1 | *JrKDM3B-like* | XP_018815718.1 |
| *PtLDL3b* | PNT47463.1 | *PtKDM3B-like* | PNT11725.1 |
| *QrLDL3* | Loc_2373_Tr_8/10_Conf_0.706_Len_11006 | *AtJMJD6B* | AT1G78280.1 |
| *CmoLDL3* | maker-scaffold00352-snap-gene-0.38 | *BrJMJD6B* | XP_009106539.1 |
| *PpLDL3* | XP_007225485.1 | *VvJMJD6B* | XP_010664345.1 |
| *PmLDL3* | XP_008221314.1 | *CmoJMJD6B* | maker-scaffold04948-snap-gene-0.21 |
| *MdLDL3a* | XP_008387975.1 | *QrJMJD6B* | OCV4_rep_c21199 |
| *MdLDL3b* | XP_017181870.1 | *JrJMJD6B* | XP_018848967.1 |
| *OsLDL3* | LOC_Os10g38850.1 | *PpJMJD6B* | XP_007227026.1 |
| *JcJMJ30* | XP_012092780.1 | *PmJMJD6B* | XP_008219330.1 |
| *VvJMJ30* | XP_010651266.1 | *MdJMJD6Ba* | XP_008372999.1 |
| *CsJMJ30* | XP_011660118.1 | *MdJMJD6Bb* | XP_017191354.1 |
| *PpJMJ30* | XP_007207823.1 | *GmJMJD6B* | XP_003526572.1 |
| *PmJMJ30* | XP_008243615.1 | *PtJMJD6B* | XP_002301069.2 |
| *OsJMJ30* | LOC_Os09g31380.1 | *JcJMJD6B* | XP_012067900.1 |
| *JrJMJ32* | XP_018814587.1 | *CsJMJD6B* | XP_011649670.1 |
| *AtJMJ32* | AT3G45880.1 | *CmJMJD6B* | XP_008444850.1 |
| *BrJMJ32* | XP_009150057.2 | *OsJMJD6B* | LOC_Os03g27250.1 |
| *CmoJMJ32* | augustus_masked-scaffold03265-abinit-gene-0.5 | *AtJMJD6A* | AT5G06550.1 |
| *QrJMJ32* | Loc_19662_Tr_4/5_Conf_0.500_Len_1503 | *BrJMJD6A* | XP_009122265.1 |
| *QrJMJ32-like* | OCV4_rep_c1929 | *CmoJMJD6A* | augustus_masked-scaffold07539-abinit-gene-0.1 |
| *PpJMJ32* | XP_007212899.1 | *QrJMJD6A* | OCV4_rep_c28281 |
| *PmJMJ32* | XP_008226259.1 | *JrJMJD6A* | XP_018851141.1 |
| *MdJMJ32* | XP_008372265.1 | *PpJMJD6A* | XP_007204829.1 |
| *VvJMJ32* | XP_010644362.1 | *PmJMJD6A* | XP_008241108.1 |
| *JcJMJ32* | XP_012067091.1 | *MdJMJD6A* | XP_008387150.1 |
| *PtJMJ32a* | PNT19189.1 | *PtJMJD6A* | PNS98217.1 |
| *PtJMJ32b* | PNT56197.1 | *JcJMJD6A* | XP_012074254.1 |
| *GmJMJ32* | XP_003538923.1 | *GmJMJD6Aa* | XP_003554495.1 |
| *OsJMJ32* | LOC_Os09g31050.1 | *GmJMJD6Ab* | XP_014629416.1 |
| *AtJMJ20* | AT5G63080.1 | *CsJMJD6A* | XP_004136539.1 |
| *BrJMJ20* | XP_009112057.1 | *CmJMJD6A* | XP_008442976.1 |
| *VvJMJ20* | XP_010663128.1 | *VvJMJD6A* | XP_002269129.1 |
| *JcJMJ20* | XP_012080013.1 | *OsJMJD6A* | LOC_Os11g36450.1 |
| *PtJMJ20* | PNT01062.1 | *AtJMJ30* | AT3G20810.1 |
| *CmoJMJ20* | maker-scaffold01888-augustus-gene-0.16 | *BrJMJ30* | XP_009145582.1 |
| *QrJMJ20* | OCV4_rep_c11681 | *GmMJ30* | XP_006590955.1 |
| *PpJMJ20* | XP_007209966.1 | *CmoJMJ30* | maker-scaffold07485-snap-gene-0.11 |
| *PmJMJ20* | XP_008244994.1 | *QrJMJ30* | OCV3_prime_c1898 |
| *MdJMJ20* | XP_008393006.1 | *PtJMJ30* | PNT52161.1 |
| *CsJMJ20* | XP_004138273.2 | *AtKDM5* | AT1G63490.1 |
| *CmJMJ20* | XP_008464540.1 | *BrKDM5* | XP_009112942.1 |
| *OsJMJ20* | LOC_Os01g36630.1 | *VvKDM5* | XP_010660757.1 |
| *PpJMJ16* | XP_020421533.1 | *QrKDM5* | Loc_1909_Tr_59/61_Conf_0.057_Len_4367 |
| *PmJMJ16* | XP_008218326.1 | *JrKDM5* | XP_018811890.1 |
| *MdJMJ16a* | XP_008370740.1 | *PpKDM5* | XP_020419317.1 |
| *MdJMJ16b* | XP_008388723.1 | *PmKDM5* | XP_008238846.1 |
| *QrJMJ16* | Loc_970_Tr_1/3_Conf_0.727_Len_4447 | *MdKDM5* | XP_008392572.1 |
| *CmoJMJ16* | maker-scaffold04342-augustus-gene-0.14 | *JcKDM5* | XP_012086900.1 |
| *JrJMJ16* | XP_018829796.1 | *PtKDM5* | PNT45248.1 |
| *VvJMJ16* | XP_002266063.2 | *GmJMJ18a* | XP_003535005.1 |
| *JcJMJ16* | XP_012089330.1 | *GmJMJ18b* | XP_006585229.1 |
| *PtJMJ16a* | PNS89961.1 | *AtJMJ18* | AT1G30810.1 |
| *PtJMJ16b* | PNT06703.1 | *BrJMJ18a* | XP_009107759.1 |
| *AtJMJ16* | AT1G08620.1 | *BrJMJ18b* | XP_009115044.1 |
| *BrJMJ16a* | XP_009110879.1 | *AtJMJ15* | AT2G34880.1 |
| *BrJMJ16b* | XP_009148114.1 | *BrJMJ15a* | XP_009141408.1 |
| *GmJMJ16a* | XP_006606422.1 | *BrJMJ15b* | XP_009141530.1 |
| *GmJMJ16b* | XP_003535393.2 | *AtJMJ14* | AT4G20400.1 |
| *CsJMJ16* | XP_004152824.1 | *BrJMJ14* | XP_009133798.1 |
| *CmJMJ16* | XP_008441838.1 | *VvJMJ18* | CBI39010.3 |
| *VvJMJ16* | XP_010652378.1 | *QrJMJ18* | Loc_9658_Tr_3/4_Conf_0.700_Len_4718 |
| *CmoJMJ19-like* | maker-scaffold08383-snap-gene-0.13 | *CmoJMJ18* | maker-scaffold01802-snap-gene-0.27 |
| *QrJMJ19-like* | OCV4_rep_c18633 | *JrJMJ18* | XP_018838947.1 |
| *JrJMJ19-like1* | XP_018827196.1 | *PpJMJ18* | XP_007213709.2 |
| *JrJMJ19-like2* | XP_018860254.1 | *MdJMJ14* | XP_008383353.1 |
| *JcJMJ19-like* | XP_012085298.1 | *JcJMJ18* | XP_012075546.1 |
| *PtJMJ19-like* | PNT47663.1 | *PtJMJ18a* | PNT13662.1 |
| *OsJMJ16* | LOC_Os05g10770.1 | *PtJMJ18b* | XP_006370484.1 |
| *JcJMJ706* | XP_012081065.1 | *OsJMJ18* | LOC_Os05g23670.1 |
| *PtJMJ706a* | PNT44650.1 | *AtREF6* | AT3G48430.1 |
| *PtJMJ706b* | PNT54370.1 | *BrREF6* | XP_009149819.1 |
| *QrJMJ706* | Loc_2899_Tr_3/5_Conf_0.417_Len_2953 | *CmoREF6* | maker-scaffold00547-snap-gene-0.28 |
| *CmoJMJ706* | maker-scaffold02280-snap-gene-0.10 | *QrREF6* | OCV4_c5323 |
| *JrJMJ706* | XP_018840882.1 | *PpREF6* | XP_020419346.1 |
| *PpJMJ706* | XP_020419985.1 | *MdREF6a* | XP_008351755.1 |
| *PmJMJ706* | XP_008238182.1 | *MdREF6b* | XP_008374335.1 |
| *MdJMJ706a* | XP_008341057.1 | *GmREF6a* | XP_003528125.1 |
| *MdJMJ706b* | XP_008373685.1 | *GmREF6b* | XP_006578679.1 |
| *OsJMJ706* | LOC_Os10g42690.1 | *CsREF6a* | XP_011651913.1 |
| *JctELF6* | XP_020532451.1 | *CmREF6b* | XP_008439230.1 |
| *PtELF6a* | PNT18125.1 | *OsREF6a* | LOC_Os01g67970.1 |
| *PtELF6b* | PNT22549.1 | *OsREF6b* | LOC_Os12g18150.1 |
| *QrELF6* | OCV4_rep_c3275 | *AtELF6* | AT5G04240.1 |
| *CmoELF6* | snap_masked-scaffold00318-abinit-gene-0.28 | *BrELF6* | XP_009122623.1 |
| *JrELF6* | XP_018828783.1 | *AtJMJ706* | AT5G46910.1 |
| *PpELF6* | XP_020412564.1 | *BrJMJ706a* | XP_009128909.1 |
| *PmELF6* | XP_008233302.1 | *BrJMJ706b* | XP_009114312.1 |
| *MdELF6* | XP_008366377.1 | *AtJMJ19* | AT2G38950.1 |
| *OsELF6* | LOC_Os03g05680.1 | *BrJMJ19a* | XP_009133319.1 |
| *GmELF6a* | XP_006606255.1 | *BrJMJ19b* | XP_018514859.1 |
| *GmELF6b* | XP_006589417.1 | *CmELF6* | XP_008456505.1 |
| *CsELF6* | XP_011657498.1 | *PtHDT1-like1* | XP_006381322.1 |
| *PtHDA19a* | XP_024464772.1 | *VvHDT1-like1* | XP_010654035.1 |
| *PtHDA19b* | XP_024455581.1 | *QrHDT1-like2* | Loc_1428_Tr_2_6_Conf_0.650_Len_1466 |
| *JcHDA19* | XP_012077112.1 | *CmoHDT1-like2* | maker-scaffold02883-augustus-gene-0.26 |
| *CmoHDA19-like1* | augustus_masked-scaffold00370-abinit-gene-0.1 | *JrHDT1-like3* | XP_018812385.1 |
| *QrHDA19-like1* | OCV4_rep_c21849 | *PpHDT1-like3* | XP_007202381.1 |
| *VvHDA19a* | XP_019080589.1 | *PmHDT1-like2* | XP_008241303.1 |
| *VvHDA19b* | XP_002283371.1 | *BrHDT3-like1* | RID42953.1 |
| *JrHDA19* | XP_018829140.1 | *BrHDT3-like2* | VDD21221.1 |
| *QrHDA19* | OCV3_primec7262 | *AtHDT3* | AT5G03740 |
| *CmoHDA19* | maker-scaffold02368-snap-gene-0.29 | *Os5-12-C2H2* | LOC_Os05g51830.1 |
| *PmHDA19* | XP_008235847.1 | *QrHDT1-like1* | Loc_8805_Tr_2_4_Conf_0.833_Len_1959 |
| *MdHDA19-like2* | XP_008348955.1 | *CmoHDT1-like1* | maker-scaffold15072-augustus-gene-0.5 |
| *MdHDA19-like1* | XP_008358446.1 | *JrHDT1-like1* | XP_018833188.1 |
| *MdHDA19a* | XP_008382752.1 | *JrHDT1-like2* | XP_018848633.1 |
| *MdHDA19b* | XP_008364370.1 | *PpHDT1-like1* | ONI01050.1 |
| *CsHDA19* | XP_011656897.1 | *PpHDT1-like2* | XP_007205675.1 |
| *CmHDA19* | XP_008448147.1 | *PmHDT1-like1* | XP_008227126.1 |
| *BrHDA19* | XP_009124989.1 | *MdHDT1-like* | XP_008355217.1 |
| *AtHDA19* | AT4G38130 | *MdHDT1-like* | XP_008385460.1 |
| *OsHDA19* | LOC_Os06g38470.3 | *GmHDT1-like1* | NP_001241996.1 |
| *OsHDA19-like1* | LOC_Os02g12350.1 | *CsHDT1-like1* | XP_004145781.1 |
| *OsHDA19-like2* | LOC_Os02g12380.1 | *BrHDT2-like2* | XP_009120640.1 |
| *QrHDA6* | OCV4_rep_c26599 | *BrHDT2-like3* | XP_009126557.1 |
| *CmoHDA6* | maker-scaffold00127-augustus-gene-0.30 | *BrHDT2-like1* | XP_009131883.1 |
| *JrHDA6a* | XP_018808184.1 | *AtHDT2* | AT5G22650 |
| *JrHDA6b* | XP_018837700.1 | *BrHDT1-like1* | RID79607.1 |
| *PtHDA6* | XP_002322192.2 | *AtHDT1* | AT3G44750 |
| *CsHDA6* | XP_004138094.1 | *AtHDT4* | AT2G27840 |
| *GmHDA6a* | XP_003549603.1 | *VvSRT1* | XP_002265837.1 |
| *GmHDA6b* | XP_003525556.1 | *JcSRT1* | XP_012071004.1 |
| *PpHDA6* | XP_007209104.1 | *CmoSRT1-like1* | maker-scaffold09820-augustus-gene-0.10 |
| *MdHDA6* | XP_008351786.1 | *PtSRT1* | XP_024449740.1 |
| *VvHDA6* | XP_010663108.1 | *CsSRT1* | XP_004148924.1 |
| *BrHDA6b* | XP_009112060.1 | *CmSRT1* | XP_008463003.1 |
| *BrHDA6a* | XP_009150370.1 | *JrSRT1* | XP_018813165.1 |
| *AtHDA6* | AT5G63110 | *QrSRT1-like1* | Loc_9090_Tr_3_7_Conf_0.611_Len_2531 |
| *OsHDA6* | LOC_Os08g25570.1 | *PpSRT1* | XP_007211402.1 |
| *GmHDA9a* | XP_003539814.1 | *PmSRT1* | XP_008227160.1 |
| *GmHDA9b* | XP_014619623.1 | *GmSRT1* | XP_003551434.1 |
| *VvHDA9a* | XP_002266492.1 | *BrSRT1* | XP_009132354.1 |
| *VvHDA9b* | XP_010651716.1 | *AtSRT1* | AT5G55760 |
| *CmoHDA9* | maker-scaffold00239-augustus-gene-1.22 | *QrSRT1-like2* | Loc_26807_Tr_3_3_Conf_0.778_Len_1873 |
| *QrHDA9-like* | OCV3_prime_rep_c68588 | *CmoSRT1-like1* | maker-scaffold10306-augustus-gene-0.6 |
| *PpHDA9* | XP_007205234.1 | *OsSRT1* | LOC_Os04g20270.1 |
| *PmHDA9* | XP_008227393.1 | *QrSRT2* | OCV3_prime_c3720 |
| *MdHDA9* | XP_008349785.1 | *CmoSRT2-like2* | maker-scaffold00071-snap-gene-1.37 |
| *JcHDA9* | XP_012075872.1 | *CmoSRt2-like1* | maker-scaffold00796-snap-gene-0.53 |
| *PtHDA9* | XP_002300554.1 | *JrSRT2* | XP_018810536.1 |
| *CsHDA9* | XP_004145792.1 | *PpSRT2* | XP_007211470.2 |
| *CmHDA9* | XP_008458605.1 | *PmSRT2* | XP_008227541.1 |
| *JrHDA9* | XP_018844531.1 | *MdSTR2* | XP_008343302.1 |
| *QrHDA9* | OCV4_rep_c14185 | *PtSRT2* | XP_002306275.2 |
| *CmoHDA9* | XP_018514955.1 | *GmSRT2a* | XP_003528059.2 |
| *CmoHDA9* | AT3G44680 | *GmSRT2b* | XP_003522478.1 |
| *AtHDA10* | AT3G44660 | *BrSRT2* | XP_018512480.1 |
| *AtHDA17* | AT3G44490 | *AtSRT2* | AT5G09230 |
| *OsHDA9\|10\|17* | LOC_Os04g33480.1 | *VvSRT2* | XP_010652926.1 |
| *BrHDA7a* | XP_018515014.1 | *OsSRT2* | LOC_Os12g07950.1 |
| *BrHDA7b* | XP_009150219.2 | *VvHDA14* | XP_002267516.1 |
| *AtHDA7* | AT5G35600 | *QrHDA14* | OCV3_prime_c9769 |
|  | augustus_masked-scaffold41118-abinit-gene-0.0 | *CmoHDA14* | maker-scaffold01029-snap-gene-0.41 |
| *VvHDA5* | XP_019082029.1 | *JrHDA14* | XP_018827767.1 |
| *QrHDA5* | Loc_6160_Tr_2_9_Conf_0.625_Len_2930 | *JcHDA14* | XP_012070591.1 |
| *CmoHDA5* | maker-scaffold00133-snap-gene-1.31 | *PtHDA14* | XP_002307035.2 |
| *JrHDA5* | XP_018824848.1 | *PpHDA14* | XP_007211505.2 |
| *PpHDA5* | XP_020420171.1 | *PmHDA14* | XP_008227633.1 |
| *PmHDA5* | XP_008240023.1 | *MdHDA14* | XP_017178842.1 |
| *MdHDA5* | XP_008374619.1 | *BrHDA14* | XP_009122119.1 |
| *JcHDA5* | XP_012080374.1 | *AtHDA14* | AT4G33470 |
| *PtHDA5* | XP_024455788.1 | *OSHDA14* | LOC_Os12g08220.1 |
| *OsHDA5* | LOC_Os07g41090.3 | *QrHDA8* | Loc_15145_Tr_8_9_Conf_0.280_Len_1833 |
| *BrHDA5-like1* | XP_009111972.1 | *CmoHDA8* | maker-scaffold26220-snap-gene-0.4 |
| *BrHDA5-like2* | XP_009136431.1 | *JrHDA8* | XP_018829758.1 |
| *AtHDA5* | AT5G61060 | *PpHDA8* | XP_007205347.1 |
| *BrHDA5-like2* | XP_009130144.1 | *PmHDA8* | XP_008230191.1 |
| *BrHDA5-like3* | XP_009130142.1 | *MdHDA8* | XP_008344371.1 |
| *AtHDA18* | AT5G61070 | *JcHDA8* | XP_012078899.1 |
| *VvHDA15* | XP_002274270.2 | *PtHDA8* | XP_002313479.1 |
| *QrHDA15* | OCV4_rep_c12421 | *OSHDA8* | LOC_Os05g36920.1 |
| *CmoHDA15* | maker-scaffold02241-augustus-gene-0.24 | *BrHDA8* | XP_009118374.1 |
| *JrHDA15* | XP_018840168. | *AtHDA8* | AT1G08460 |
| *PpHDA15* | XP_020419500.1 | *CmoHDA8-like1* | maker-scaffold13571-augustus-gene-0.6 |
| *PmHDA15* | XP_008244027.1 | *CmoHDA8-like2* | maker-scaffold17774-augustus-gene-0.3 |
| *MdHDA15a* | XP_008374725.1 | *VvHDA2* | XP_002277742.1 |
| *MdHDA15b* | XP_008393285.1 | *PtHDA2* | XP_006382022.1 |
| *JcHDA15a* | XP_012092970.1 | *JcHDA2* | XP_020536570.1 |
| *JcHDA15b* | XP_012092971.1 | *QrHDA2* | OCV4_rep_c15012 |
| *PtHDA15* | XP_024438220.1 | *CmoHDA2* | maker-scaffold01226-snap-gene-0.40 |
| *BrHDA15* | XP_009145883.1 | *PpHDA2* | XP_007225703.1 |
| *AtHDA15* | AT3G18520 | *PmHDA2* | XP_008220462.1 |
| *CsHDA15* | XP_004139132.1 | *GmHDA2a* | XP_003550277.1 |
| *CmHDA15* | XP_008450327.1 | *GmHDA2b* | XP_006601239.1 |
| *OsHDA15* | LOC_Os07g06980.1 | *CsHDA2* | XP_004141312.1 |
| *AtHDA2* | AT5G26040 | *CmHDA2* | XP_008452709.1 |
| *OsHDA2* | LOC_Os06g37420.1 | *BrHDA2* | XP_009151116.2 |

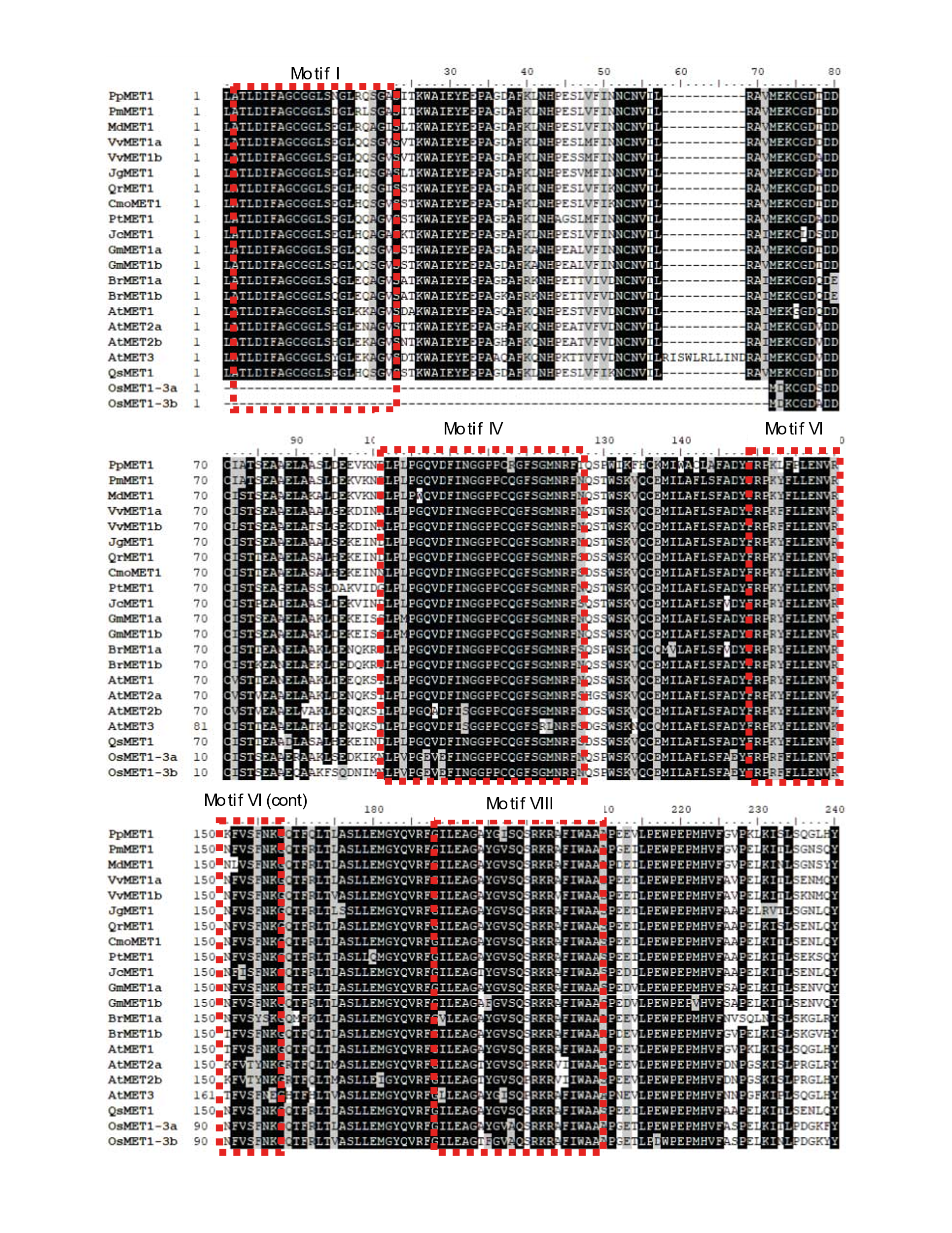

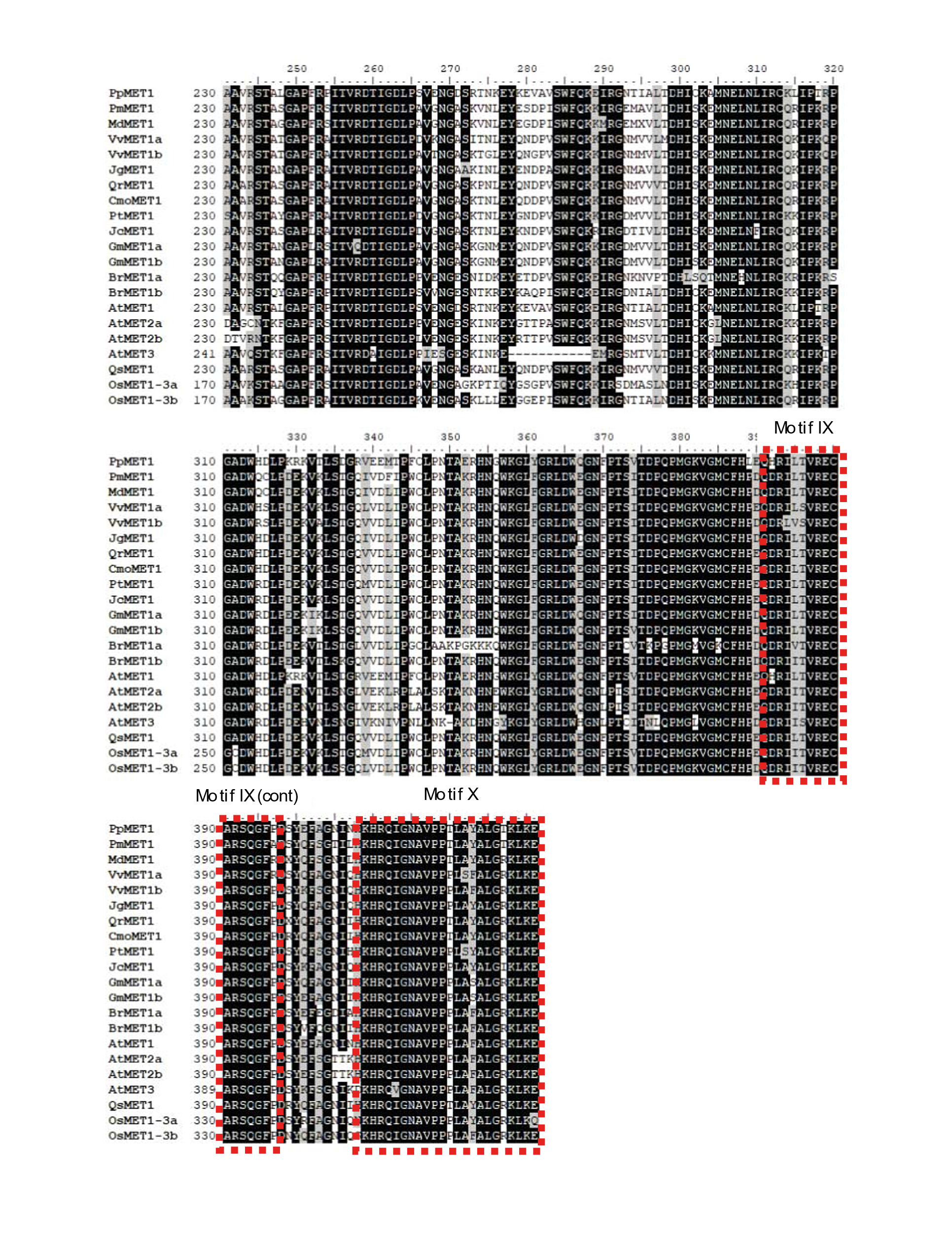

**Supplementary Figure S1. Alignment of the amino acid sequences of MET family proteins.** Dotted red boxes indicate the conserved methyltransferase catalytic motifs I, IV, and VI, VIII-X. Black background shows identical amino acid sequences among MET1 proteins.

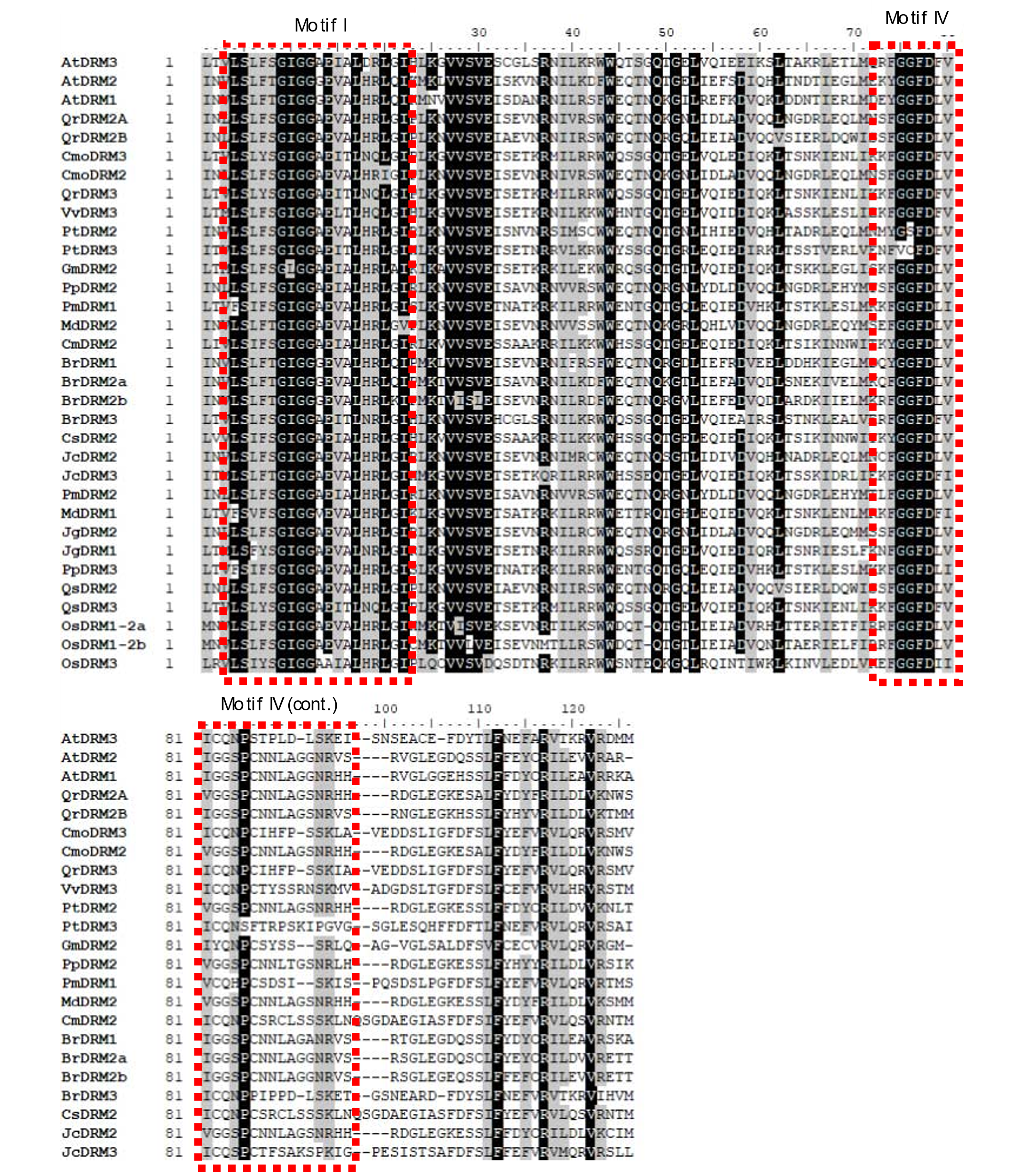

**Supplementary Figure S2. Alignment of the amino acid sequences of DRM family proteins.** Dotted red boxes indicate the conserved methyltransferase catalytic motifs I, and IV. Black background shows identical amino acid sequences among DRM proteins.

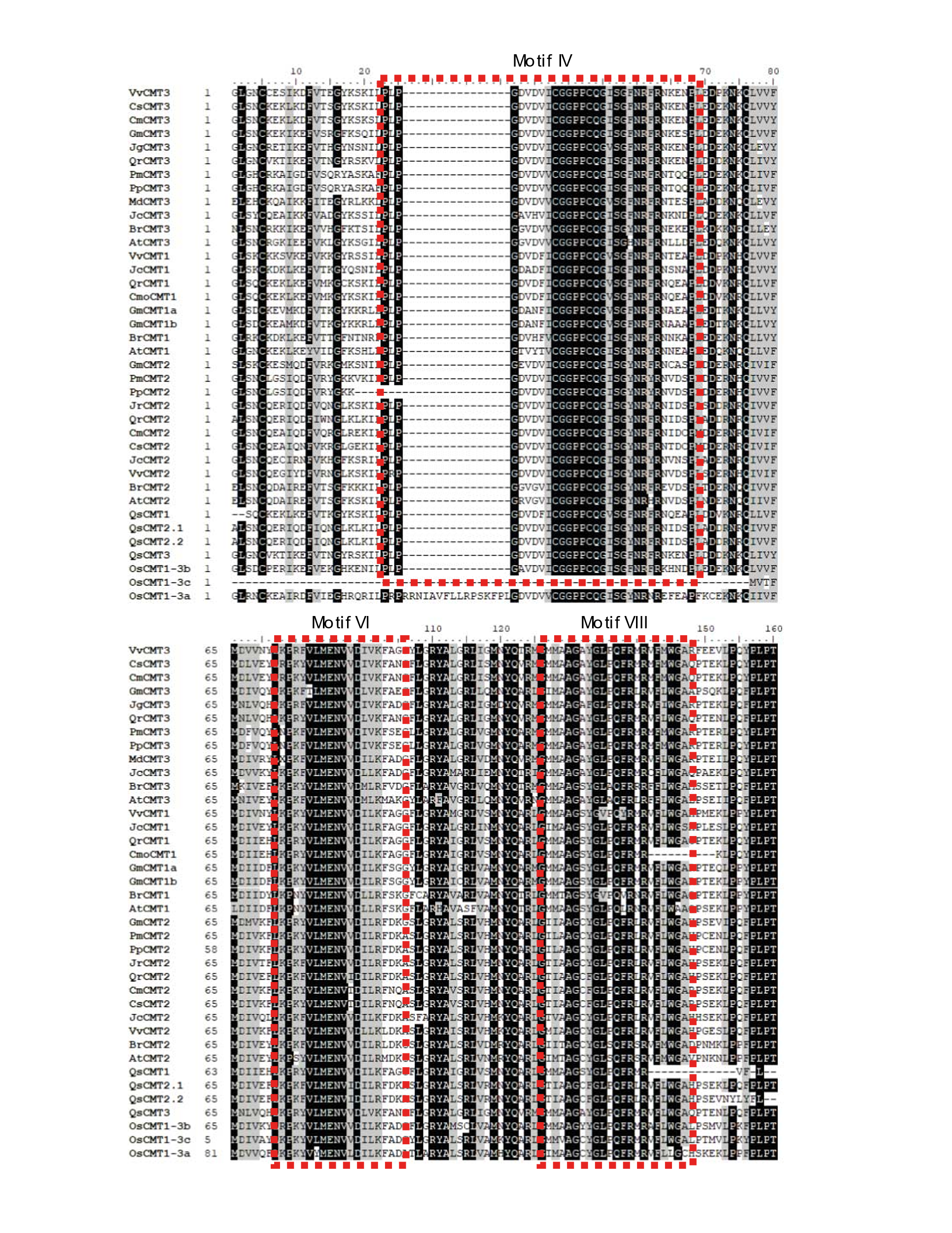

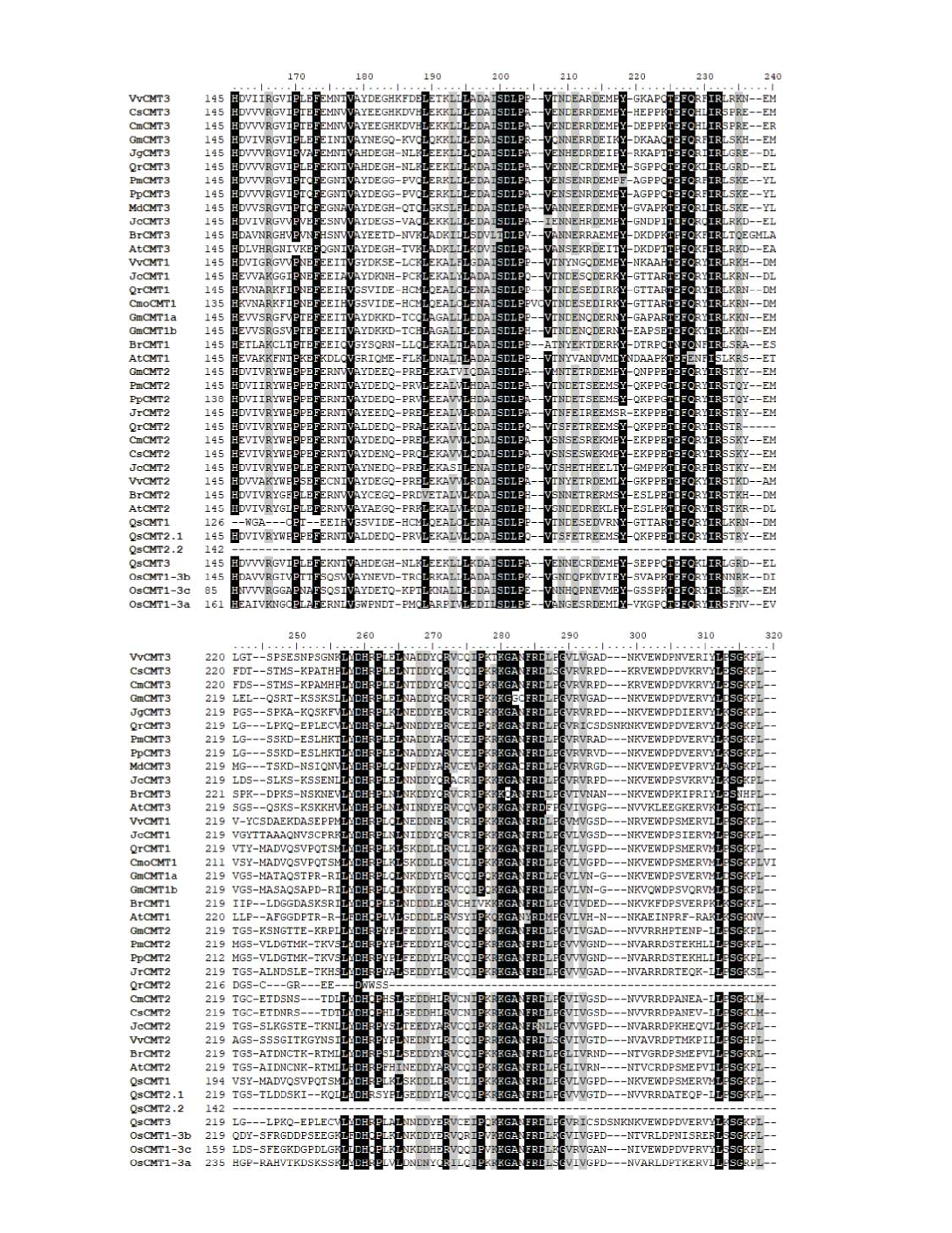

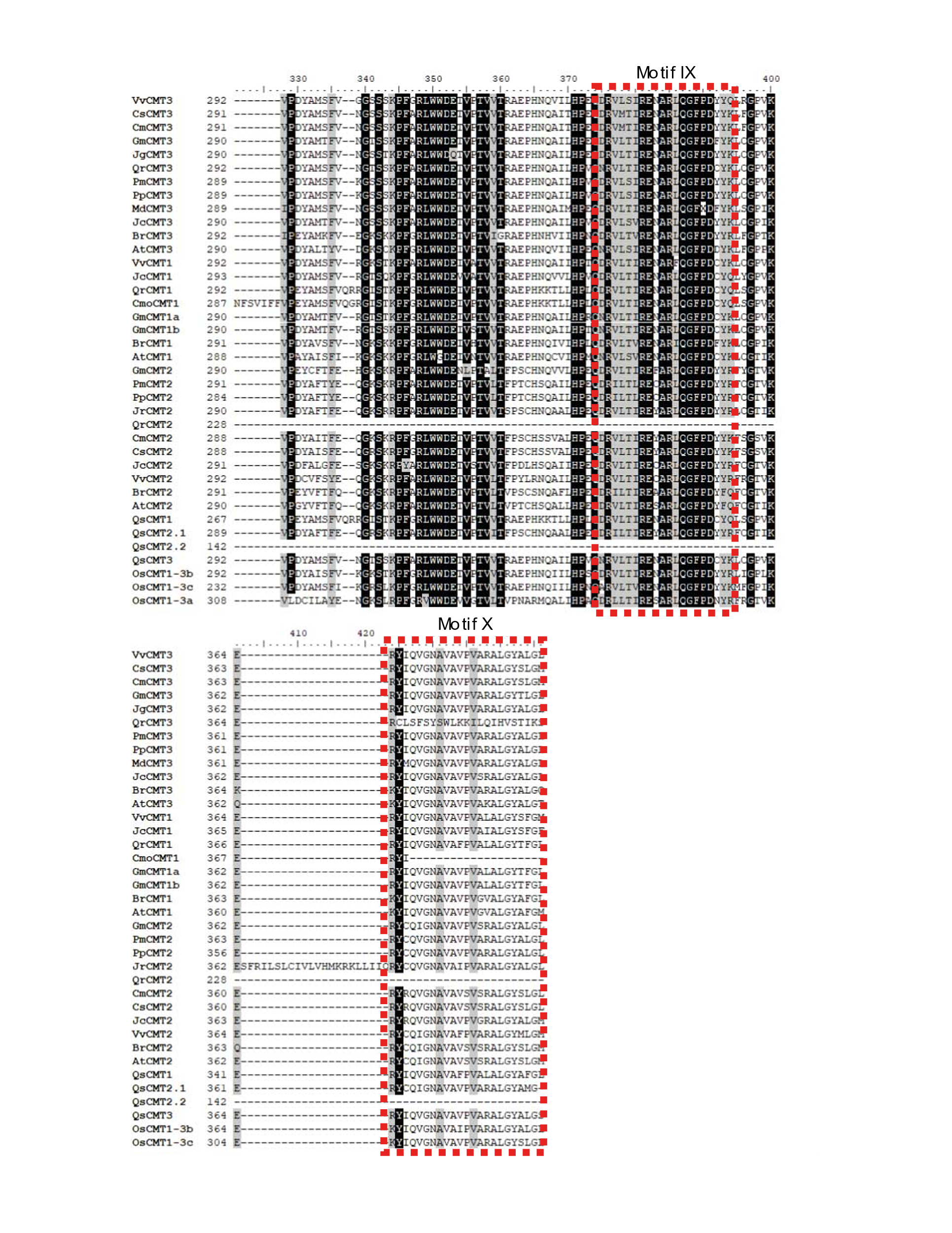

**Supplementary Figure S3.** **Alignment of the amino acid sequences of CMT family proteins.** Dotted red boxes indicate the conserved methyltransferase catalytic motifs IV, VI and VIII-X. Black background shows identical amino acid sequences among CMT proteins.

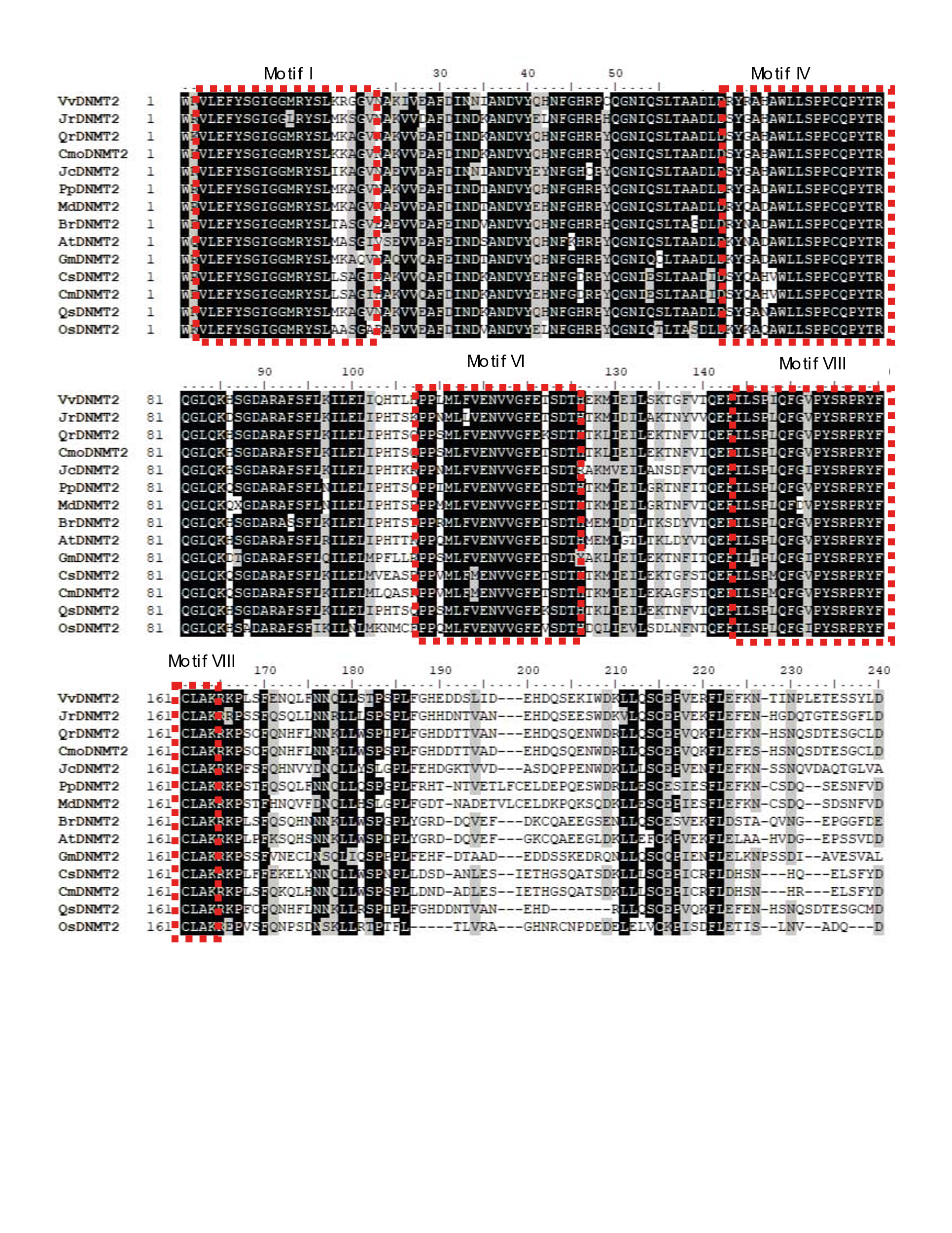

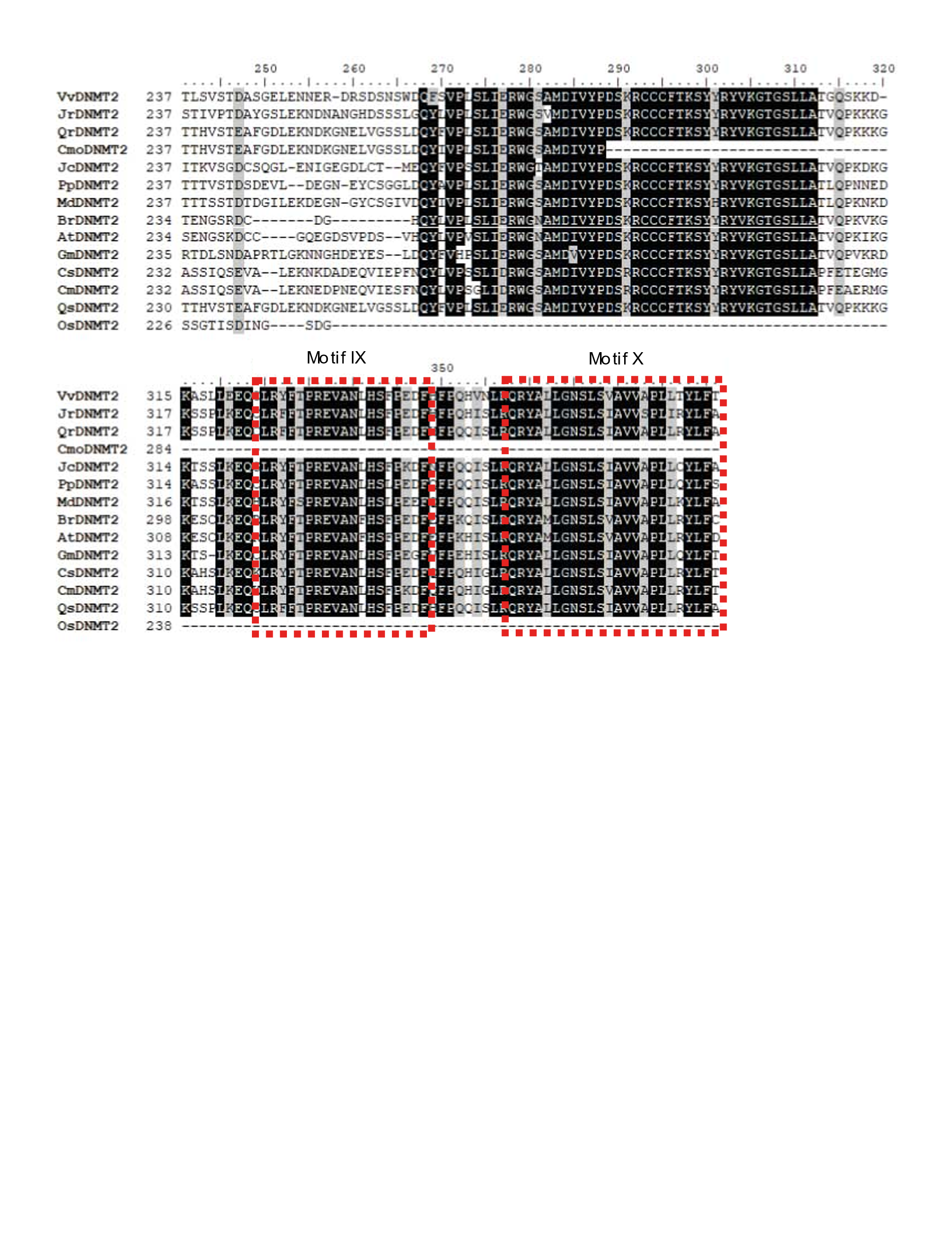

**Supplementary Figure S4.** **Alignment of the amino acid sequences of DNMT family proteins.** Dotted red boxes indicate the conserved methyltransferase catalytic motifs I, IV, VI and VIII-X. Black background shows identical amino acid sequences among DNMT proteins.

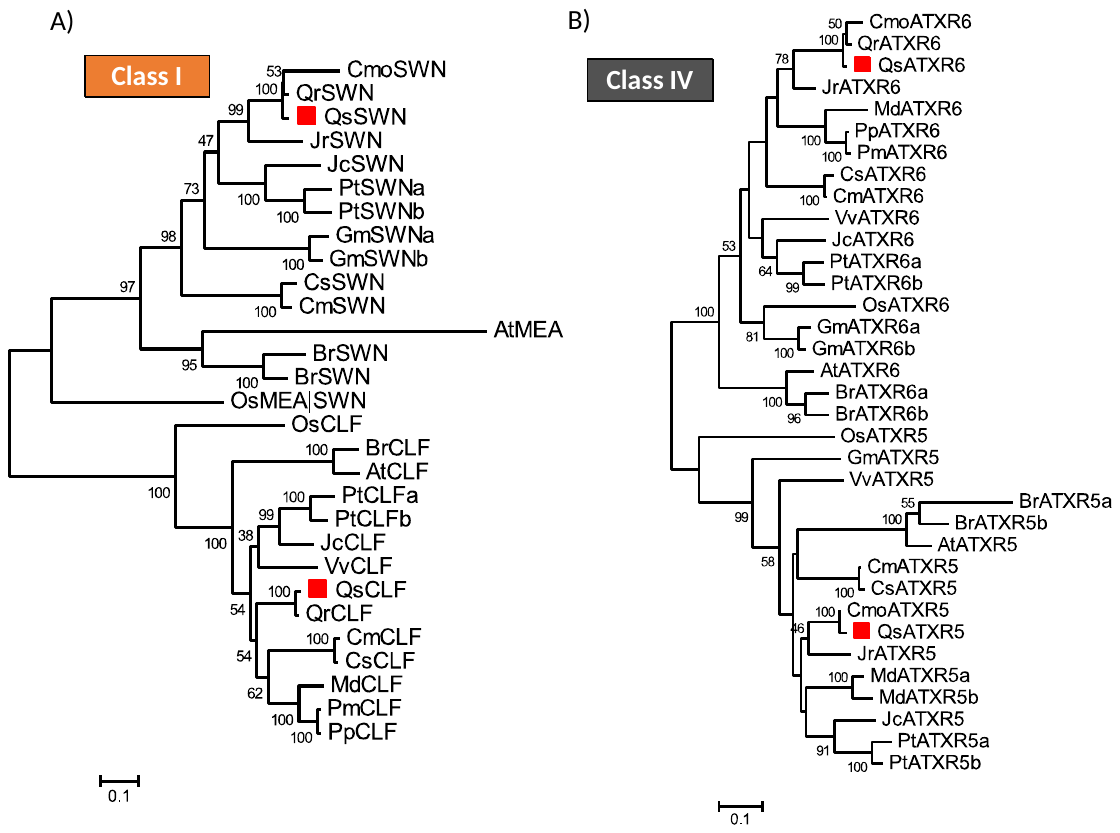

**Supplementary Figure S5.** **Phylogeny of HMT proteins of class I and IV.** The SET domain of *Quercus suber* (Qs), *Arabidopsis thaliana* (At), *Brassica rapa* (Br), *Glycine max* (Gm), *Jatropha curcas* (Jc), *Populus trichocarpa* (Pt), *Prunus persica* (Pp), *Prunus mume* (Pm), *Vitis vinifera* (Vv), *Cucumis melo* (Cm), *Cucumis sativus* (Cs), *Juglans regia* (Jr), *Castanea mollissima* (Cmo), *Quercus robur* (Qr) *Oryza sativa* (Os) were aligned using ClustalW and used to infer the evolutionary history using the Maximum-likelihood method. The evolutionary distances (left side scale bar) were computed using the Jones-Taylor-Thornton (JTT) correction model. The numbers at the nodes represent bootstrap values from 1000 replicates. The *Quercus suber* SET domain contain proteins were indicated with a red square. Phylogenetic analyses were conducted in MEGA7.

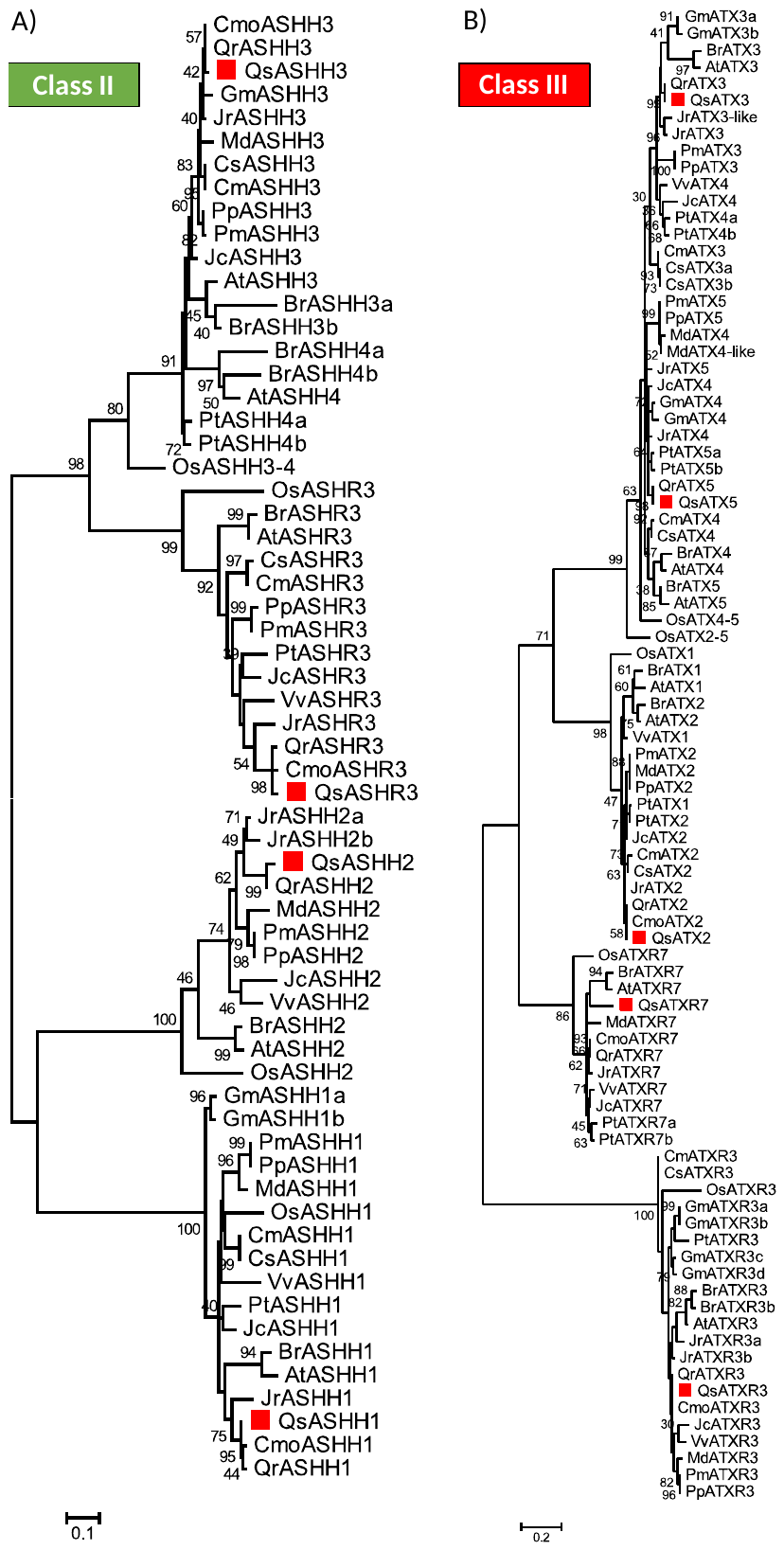

**Supplementary Figure S6. Phylogeny of HMT proteins of class II and III.** The SET domain of *Quercus suber* (Qs), *Arabidopsis thaliana* (At), *Brassica rapa* (Br), *Glycine max* (Gm), *Jatropha curcas* (Jc), *Populus trichocarpa* (Pt), *Prunus persica* (Pp), *Prunus mume* (Pm), *Vitis vinifera* (Vv), *Cucumis melo* (Cm), *Cucumis sativus* (Cs), *Juglans regia* (Jr), *Castanea mollissima* (Cmo), *Quercus robur* (Qr) *Oryza sativa* (Os) were aligned using ClustalW and used to infer the evolutionary history using the Maximum-likelihood method. The evolutionary distances (left side scale bar) were computed using the Jones-Taylor-Thornton (JTT) correction model. The numbers at the nodes represent bootstrap values from 1000 replicates. The *Quercus suber* SET domain contain proteins were indicated with a red square. Phylogenetic analyses were conducted in MEGA7.

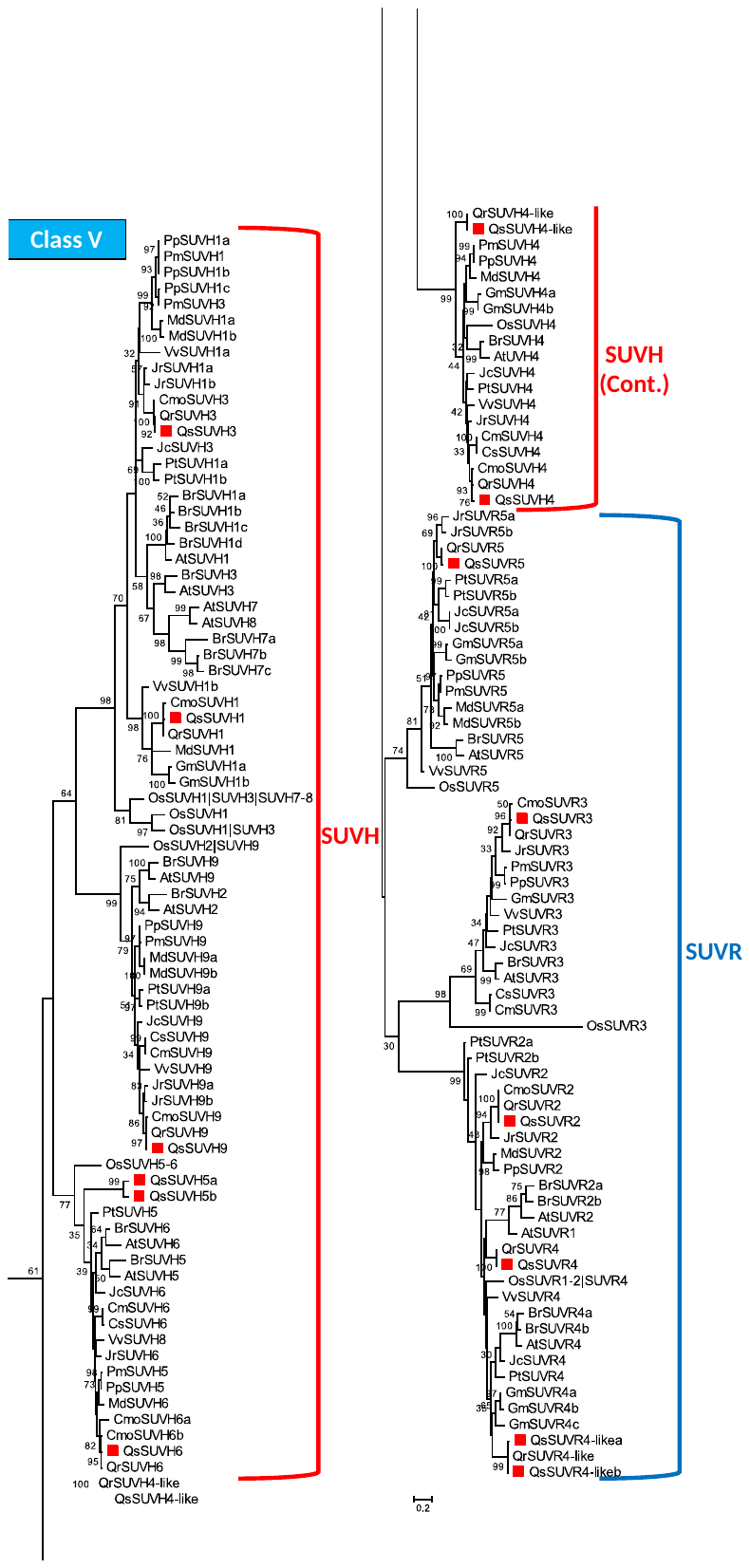

**Supplementary Figure S7. Phylogeny of HMT proteins of class V.** The SET domain of *Quercus suber* (Qs), *Arabidopsis thaliana* (At), *Brassica rapa* (Br), *Glycine max* (Gm), *Jatropha curcas* (Jc), *Populus trichocarpa* (Pt), *Prunus persica* (Pp), *Prunus mume* (Pm), *Vitis vinifera* (Vv), *Cucumis melo* (Cm), *Cucumis sativus* (Cs), *Juglans regia* (Jr), *Castanea mollissima* (Cmo), *Quercus robur* (Qr) *Oryza sativa* (Os) were aligned using ClustalW and used to infer the evolutionary history using the Maximum-likelihood method. The evolutionary distances (left side scale bar) were computed using the Jones-Taylor-Thornton (JTT) correction model. The numbers at the nodes represent bootstrap values from 1000 replicates. The *Quercus suber* SET domain contain proteins were indicated with a red square. Phylogenetic analyses were conducted in MEGA7.

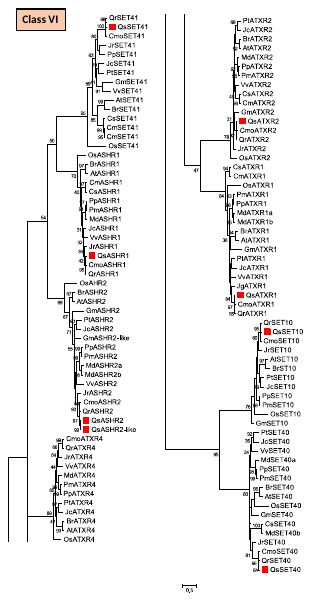

**Supplementary Figure S8. Phylogeny of other SET containing proteins.** The SET domain of *Quercus suber* (Qs), *Arabidopsis thaliana* (At), *Brassica rapa* (Br), *Glycine max* (Gm), *Jatropha curcas* (Jc), *Populus trichocarpa* (Pt), *Prunus persica* (Pp), *Prunus mume* (Pm), *Vitis vinifera* (Vv), *Cucumis melo* (Cm), *Cucumis sativus* (Cs), *Juglans regia* (Jr), *Castanea mollissima* (Cmo), *Quercus robur* (Qr) *Oryza sativa* (Os) were aligned using ClustalW and used to infer the evolutionary history using the Maximum-likelihood method. The evolutionary distances (left side scale bar) were computed using the Jones-Taylor-Thornton (JTT) correction model. The numbers at the nodes represent bootstrap values from 1000 replicates. The *Quercus suber* SET domain contain proteins were indicated with a red square. Phylogenetic analyses were conducted in MEGA7.

**Supplementary Figure S9.** **Phylogeny of HDMT proteins of JmJC family.** The JmJC domain of *Quercus suber* (Qs), *Arabidopsis thaliana* (At), *Brassica rapa* (Br), *Glycine max* (Gm), *Jatropha curcas* (Jc), *Populus trichocarpa* (Pt), *Prunus persica* (Pp), *Prunus mume* (Pm), *Vitis vinifera* (Vv), *Cucumis melo* (Cm), *Cucumis sativus* (Cs), *Juglans regia* (Jr), *Castanea mollissima* (Cmo), *Quercus robur* (Qr) *Oryza sativa* (Os) were aligned using ClustalW and used to infer the evolutionary history using the Maximum-likelihood method. The evolutionary distances (left side scale bar) were computed using the Jones-Taylor-Thornton (JTT) correction model. The numbers at the nodes represent bootstrap values from 1000 replicates. The *Quercus suber* proteins were indicated with a red square. Phylogenetic analyses were conducted in MEGA7.

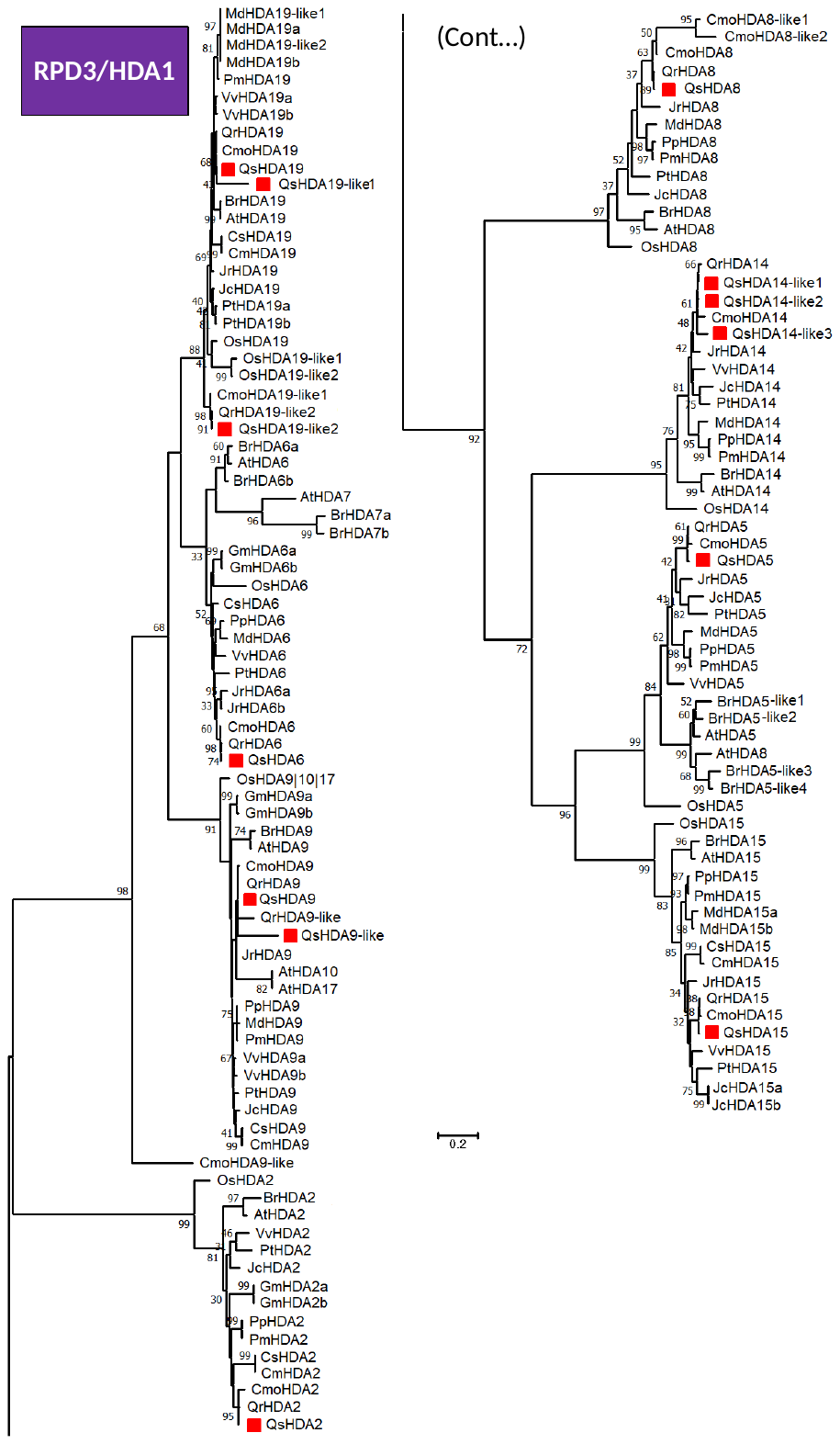

**Supplementary Figure S10. Phylogeny of HDAC proteins of RPD3/HDA1 family.** The HDAC domain of *Quercus suber* (Qs), *Arabidopsis thaliana* (At), *Brassica rapa* (Br), *Glycine max* (Gm), *Jatropha curcas* (Jc), *Populus trichocarpa* (Pt), *Prunus persica* (Pp), *Prunus mume* (Pm), *Vitis vinifera* (Vv), *Cucumis melo* (Cm), *Cucumis sativus* (Cs), *Juglans regia* (Jr), *Castanea mollissima* (Cmo), *Quercus robur* (Qr) *Oryza sativa* (Os) were aligned using ClustalW and used to infer the evolutionary history using the Maximum-likelihood method. The evolutionary distances (left side scale bar) were computed using the Jones-Taylor-Thornton (JTT) correction model. The numbers at the nodes represent bootstrap values from 1000 replicates. The *Quercus suber* proteins were indicated with a red square. Phylogenetic analyses were conducted in MEGA7.

**Table S1.** **The identification of the loci encoding DNA and histone epigenetic modifier homologs of *Quercus suber*.** Gene name, the correspondent locus tag, gene symbol and protein IDs (including splicing forms) are represented.

| **Gene name** | **Locus tag** | **Gene symbol** | **Proteins** |
| --- | --- | --- | --- |
| **DNA Methyltransferases** | | | |
| ***QsMET1*** | CFP56_38722 | LOC111984282 | XP_023871673.1 |
| ***QsCMT1*** | CFP56_24702 | LOC112002245 | XP_023890169.1  XP_023890170.1 |
| ***QsCMT2.1*** | CFP56_06331 | LOC112001521 | XP_023889464.1  XP_023889465.1 |
| ***QsCMT3*** | CFP56_35720 | LOC112031183 | XP_023919638.1 |
| ***QsDRM2*** | CFP56_35392 | LOC112033986 | XP_023922537.1 |
| ***QsDRM3*** | CFP56_66797 | LOC112032791 | XP_023921322.1  XP_023921323.1  XP_023921324.1  XP_023921321.1 |
| ***QsDNMT2*** | CFP56_40985 | LOC112003961 | XP_023891955.1 |
| ***QsCMT2.2*** | CFP56_59471 | LOC112034184 | XP_023922750.1 |
| **DNA Demethylases** | | | |
| ***QsDME*** | CFP56_06273 | LOC111983123 | XP_023870550.1 |
| ***QsROS1*** | CFP56_05353 | LOC112039671 | XP_023928322.1 |
| ***QsDML2*** | CFP56_35285 | LOC112020085 | XP_023908412.1 |
| **Histone Acetyltransferases** | | | |
| ***QsHAC1*** | CFP56_16103 | LOC112034234 | XP_023922811.1  XP_023922818.1 |
| ***QsHAC1-like1*** | CFP56_07357 | LOC111988497 | XP_023876070.1 |
| ***QsHAC1-like2*** | CFP56_07360 | LOC111988316 | XP_023875877.1 |
| ***QsGCN5*** | CFP56_29027 | LOC111998855 | XP_023886737.1 |
| ***QsHAG2*** | CFP56_43084 | LOC112010794 | XP_023898930.1 |
| ***QsHAM1*** | CFP56_58450 | LOC112012489 | XP_023900637.1 |
| ***QsHAF1*** | CFP56_26298 | LOC111991837 | XP_023879413.1  XP_023879414.1 |
| ***QsELP3*** | CFP56_52694 | LOC112000045 | XP_023887919.1 |
| **Histone Methyltransferases** | | | |
| ***QsSET40*** | CFP56_02401 | LOC111984868 | XP_023872255.1 |
| ***QsASHR2*** | CFP56_23528 | LOC111985014 | XP_023872401.1 |
| ***QsASHR2-like*** | CFP56_23526 | LOC111984972 | XP_023872355.1 |
| ***QsSUVR5*** | CFP56_22604 | LOC111985175 | XP_023872585.1 |
| ***QsCLF*** | CFP56_48440 | LOC111988803 | XP_023876366.1  XP_023876365.1 |
| ***QsATX3*** | CFP56_04889 | LOC111989353 | XP_023876913.1 |
| ***QsSUVH6*** | CFP56_40587 | LOC111989703 | XP_023877264.1  XP_023877266.1  XP_023877265.1 |
| ***QsSUVR2*** | CFP56_56207 | LOC111992201 | XP_023879822.1  XP_023879817.1  XP_023879818.1  XP_023879819.1  XP_023879820.1  XP_023879823.1 |
| ***QsATX2*** | CFP56_26820 | LOC111994171 | XP_023881801.1  XP_023881799.1  XP_023881802.1  XP_023881800.1 |
| ***QsATXR2*** | CFP56_29061 | LOC111996077 | XP_023883778.1 |
| ***QsSUVR3*** | CFP56_29051 | LOC111996085 | XP_023883794.1 |
| ***QsSUVH9*** | CFP56_44980 | LOC111996522 | XP_023884276.1  XP_023884277.1 |
| ***QsASHH2*** | CFP56_13855 | LOC111999241 | XP_023887134.1  XP_023887133.1 |
| ***QsSUVR4*** | CFP56_60043 | LOC111999549 | XP_023887448.1 |
| ***QsASHH3*** | CFP56_26485 | LOC112002855 | XP_023890787.1  XP_023890783.1  XP_023890785.1  XP_023890784.1  XP_023890788.1 |
| ***QsATXR1*** | CFP56_15756 | LOC112005587 | XP_023893650.1 |
| ***QsSUVH4-like*** | CFP56_01500 | LOC112036034 | XP_023924619.1 |
| ***QsATXR6*** | CFP56_57230 | LOC112008849 | XP_023896969.1 |
| ***QsASHR3*** | CFP56_53193 | LOC112009409 | XP_023897510.1 |
| ***QsATXR7*** | CFP56_14674 | LOC112011828 | XP_023899934.1  XP_023899935.1 |
| ***QsATX5*** | CFP56_25529 | LOC112017971 | XP_023906233.1  XP_023906232.1 |
| ***QsSUVH1*** | CFP56_34169 | LOC112020905 | XP_023909243.1 |
| ***QsSUVH3*** | CFP56_28232 | LOC112025017 | XP_023913440.1 |
| ***QsASHH1*** | CFP56_54040 | LOC112035360 | XP_023923957.1  XP_023923958.1  XP_023923956.1  XP_023923959.1 |
| ***QsSUVH4*** | CFP56_37589 | LOC112035570 | XP_023924166.1  XP_023924168.1  XP_023924167.1 |
| ***QSASHR1*** | CFP56_31286 | LOC112037474 | XP_023926081.1 |
| ***QsATXR5*** | CFP56_53906 | LOC112038577 | XP_023927165.1 |
| ***QsSWN*** | CFP56_32659 | LOC112038731 | XP_023927343.1 |
| ***QsSET41*** | CFP56_40485 | LOC112039440 | XP_023928085.1  XP_023928086.1 |
| ***QsSUVH5b*** | CFP56_65722 | LOC112015057 | XP_023903188.1 |
| ***QsSUVH5a*** | CFP56_65724 | LOC112015060 | XP_023903190.1 |
| ***QsSUVR4-likeb*** | CFP56_76543 | LOC112015520 | XP_023903694.1 |
| ***QsSUVR4-likea*** | CFP56_36057 | LOC112032443 | XP_023920971.1 |
| ***QsSET10*** | CFP56_45284 | LOC112020643 | XP_023908960.1 |
| ***QSATXR3*** | CFP56_27073 | - | QSP042949.0 |
| **Histone Demethylases** | | | |
| ***QsJMJ18*** | CFP56_59257 | LOC111985798 | XP_023873223.1  XP_023873225.1  XP_023873224.1 |
| ***QsKDM3B-like*** | CFP56_10640 | LOC111989488 | XP_023877041.1  XP_023877042.1  XP_023877040.1 |
| ***QsJMJ25-like1*** | CFP56_48908 | LOC112003991 | XP_023891992.1  XP_023891991.1  XP_023891990.1 |
| ***QsJMJ25-like2*** | CFP56_47938 | LOC112018887 | XP_023907188.1 |
| ***QsJMJ25-like3*** | CFP56_18193 | LOC112030617 | XP_023919056.1 |
| ***QsJMJ25-like4*** | CFP56_18195 | LOC112030626 | XP_023919066.1  XP_023919065.1 |
| ***QsJMJ24*** | CFP56_37500 | LOC111995675 | XP_023883369.1  XP_023883368.1  XP_023883365.1  XP_023883367.1 |
| ***QsKDM3D-like*** | CFP56_34178 | LOC112020902 | XP_023909239.1 |
| ***QsJMJD6Aa*** | CFP56_37891 | LOC111991092 | XP_023878641.1 |
| ***QsJMJD6Ab*** | CFP56_50389 | LOC112013333 | XP_023901496.1 |
| ***QsJMJ16*** | CFP56_76189 | LOC112004294 | XP_023892291.1  XP_023892289.1  XP_023892290.1 |
| ***QsJMJ19-like*** | CFP56_17215 | LOC112019264 | XP_023907567.1  XP_023907566.1 |
| ***QsJMJ706*** | CFP56_36127/6 | LOC112010540 | XP_023898652.1  XP_023898654.1  XP_023898653.1 |
| ***QsJMJ706-like*** | CFP56_37564 | LOC112030921 | XP_023919361.1 |
| ***QsLDL3*** | CFP56_08883 | LOC112022596 | XP_023910985.1 |
| ***QsELF6*** | CFP56_41197 | LOC112025120 | XP_023913552.1 |
| ***QsJMJD6B*** | CFP56_61743 | LOC112026327 | XP_023914775.1  XP_023914776.1 |
| ***QsJMJ32-like*** | CFP56_26267 | LOC112034921 | XP_023923513.1 |
| ***QsJMJ32*** | CFP56_26269 | LOC112034640 | XP_023923230.1 |
| ***QsREF6*** | CFP56_69927 | LOC112029048 | XP_023917830.1 |
| ***QsLDL2*** | CFP56_19117 | LOC112032113 | XP_023920630.1  XP_023920629.1 |
| ***QsLDL1*** | CFP56_55051 | LOC112033571 | XP_023922114.1 |
| ***QsKDM5*** | CFP56_46886 | LOC112036450 | XP_023925013.1 |
| ***QsJMJ30*** | CFP56_23213 | LOC112002893 | XP_023890817.1 |
| ***QsFLD*** | CFP56_03400 | LOC112023717 | XP_023912111.1 |
| ***QsJMJ20*** | CFP56_45215 | LOC111991489 | XP_023879031.1 |
| **Histone Deacetylases** | | | |
| ***QsHDA2*** | CFP56_55621 | LOC112000524 | XP_023888449.1 |
| ***QsHDA5*** | CFP56_01537 | LOC112036007 | XP_023924594.1 |
| ***QsHDA6*** | CFP56_50628 | LOC112035676 | XP_023924274.1 |
| ***QsHDA19-like2*** | CFP56_14676 | LOC112011819 | XP_023899923.1  XP_023899924.1  XP_023899925.1 |
| ***QsHDA19*** | CFP56_26357 | LOC112003171 | XP_023891130.1  XP_023891137.1 |
| ***QsHDA9*** | CFP56_22884 | LOC111996728 | XP_023884497.1  XP_023884496.1 |
| ***QsHDA19-like1*** | CFP56_75988 | LOC112009697 | XP_023897791.1 |
| ***QsHDA8*** | CFP56_03621 | LOC112005173 | XP_023893191.1 |
| ***QsHDA14-like1*** | CFP56_64318 | LOC112012023 | XP_023900151.1  XP_023900150.1  XP_023900153.1  XP_023900152.1 |
| ***QsHDA14-like2*** | CFP56_32080 | LOC112004322 | XP_023892317.1 |
| ***QsHDA14-like3*** | CFP56_65290 | LOC112011809 | XP_023899914.1 |
| ***QsHDA15*** | CFP56_10834 | LOC112024699 | XP_023913091.1  XP_023913093.1  XP_023913092.1 |
| ***QsHDT1-like1*** | CFP56_42573 | LOC112002966 | XP_023890901.1 |
| ***QsHDT1-like3*** | CFP56_05349 | LOC112039702 | XP_023928367.1 |
| ***QsSRT1-like3*** | CFP56_78126 | LOC112020807 | XP_023909132.1 |
| ***QsSRT1-like5*** | CFP56_38596 | LOC112028975 | XP_023917433.1 |
| ***QsSRT1-like4*** | CFP56_42400 | LOC112017795 | XP_023906010.1  XP_023906011.1 |
| ***QsSRT1-like2*** | - | LOC112025841 | XP_023914295.1 |
| ***QsSRT1-like1*** | - | LOC112027314 | XP_023915762.1 |
| ***QsSRT2*** | CFP56_42602 | LOC112032808 | XP_023921347.1  XP_023921348.1 |
| ***QsHDT1-like1*** | CFP56_51836 | LOC112035297 | XP_023923893.1 |
| ***QsHDA9-like*** | - | [LOC112031837](https://www.ncbi.nlm.nih.gov/sites/entrez?db=gene&cmd=retrieve&list_uids=112031837) | XP_023920316.1 |

| **Table S2: Normalized read counts of DNA De/methyltransferases and histone modifiers detected in ecodormant bud (ecodB), swelling bud (swB), differentiating secondary xylem (XY), root (RO), leaf (LE) and in vitro dedifferentiated callus (CA) of *Quercus robur*** [35]**.** | | | | | | | |
| --- | --- | --- | --- | --- | --- | --- | --- |
| ***Gene name*** | **Gene ID** | **EcoDB** | **swB** | **XY** | **RO** | **LE** | **CA** |
| **DNA methyltransferases** | | | | | | | |
| *QrCMT3* | Loc_5202_Tr_1/2_Conf_0.889_Len_3208 | 0.0 | 44.9 | 0.8 | 3.1 | 7.8 | 4.1 |
| *QrCMT1* | Loc_21057_Tr_1/5_Conf_0.667_Len_2822 | 7.7 | 185.6 | 5.9 | 96.7 | 45.6 | 54.0 |
| *QrCMT2* | OCV4_rep_c4395 | ND | ND | ND | ND | ND | ND |
| *QrMET1* | OCV4_c25957 | 874.6 | 2363.3 | 811.8 | 819.9 | 717.4 | 828.9 |
| *QrDNMT2* | OCV4_rep_c6323/Loc_8252_Tr_6/6_Conf_0.600_Len_1795 | 1.9 | 0.8 | 2.5 | 3.1 | 1.3 | 1.0 |
| *QrDRM2A* | Loc_1382_Tr_6/6_Conf_0.667_Len_2740 | 8.7 | 20.8 | 16.0 | 9.3 | 9.1 | 15.3 |
| *QrDRM2B* | Loc_82954_Tr_2/2_Conf_0.750_Len_1158 | 1.9 | 0.0 | 3.4 | 0.0 | 0.0 | 0.0 |
| *QrDRM3* | OCV4_rep_c18566 | ND | ND | ND | ND | ND | ND |
| **Histone acetyltransferases** | | | | | | | |
| *QrHAC1* | OCV4_rep_c42029 | ND | ND | ND | ND | ND | ND |
| *QrHAC1-like2* | Loc_39724_Tr_1/2_Conf_0.750_Len_4836 | 259.5 | 61.6 | 174.7 | 143.0 | 148.4 | 24.4 |
| *QrHAM1* | OCV3_prime_c11529 | 366.2 | 435.2 | 232.1 | 524.7 | 623.6 | 299.4 |
| *QrHAG2* | OCV4_rep_c26572 | ND | ND | ND | ND | ND | ND |
| *QrELP3* | Loc_4308_Tr_2/4_Conf_0.600_Len_2295 | 74.0 | 59.9 | 123.2 | 94.6 | 74.2 | 279.0 |
| *QrGCN5* | OCV4_c7575 | 938.1 | 1235.8 | 966.2 | 825.1 | 757.7 | 1486.7 |
| *QrHAF1* | OCV4_rep_c6162 | ND | ND | ND | ND | ND | ND |
| **Histone methyltransferases** | | | | | | | |
| *QrSWN* | Loc_1192_Tr_4/8_Conf_0.696_Len_3240 | 26.0 | 10.8 | 27.0 | 18.5 | 13.0 | 26.5 |
| *QrCLF* | OCV4_rep_c16251/Loc_19971_Tr_7/10_Conf_0.467_Len_2918 | 16.3 | 24.1 | 12.7 | 14.4 | 5.2 | 6.1 |
| *QrASHH1* | Loc_12859_Tr_1/4_Conf_0.769_Len_2491 | 36.5 | 20.8 | 30.4 | 26.7 | 7.8 | 12.2 |
| *QrASHH3* | OCV4_rep_c40652/OCV4_rep_c14852 | 83.6 | 66.6 | 75.1 | 67.9 | 72.9 | 116.1 |
| *QrASHR3* | OCV4_rep_c19155 | ND | ND | ND | ND | ND | ND |
| *QrASHH2* | OCV4_rep_c21713 | ND | ND | ND | ND | ND | ND |
| *QrATXR7* | Loc_9902_Tr_7/8_Conf_0.682_Len_7488 | 100.0 | 34.1 | 61.6 | 39.1 | 57.3 | 73.3 |
| *QrATX3* | Loc_20589_Tr_3/6_Conf_0.688_Len_3877 | 936.2 | 912.9 | 813.5 | 788.0 | 1070.2 | 643.6 |
| *QrATX5* | OCV4_c26314 | 353.7 | 290.4 | 505.5 | 259.2 | 251.3 | 242.4 |
| *QrATX2* | OCV4_rep_c43523 | ND | ND | ND | ND | ND | ND |
| *QrATXR3* | OCV4_rep_c4016 | ND | ND | ND | ND | ND | ND |
| *QrATXR6* | OCV4_rep_c10294 | 0.0 | 94.9 | 0.8 | 11.3 | 29.9 | 8.1 |
| *QrSUVR3* | OCV4_rep_c24922 | ND | ND | ND | ND | ND | ND |
| *QsSUVH6* | OCV4_c18 | 894.8 | 1056.0 | 1145.1 | 667.7 | 820.2 | 809.5 |
| *QrSUVH4* | OCV4_rep_c40958/OCV4_rep_c14813 | 380.6 | 687.4 | 457.4 | 475.3 | 407.5 | 280.0 |
| *QrSUVH4-like* | Loc_19624_Tr_8/9_Conf_0.609_Len_2652 | 34.6 | 51.6 | 34.6 | 32.9 | 52.1 | 44.8 |
| *QrSUVH1* | OCV4_c19481 | 1715.7 | 1368.1 | 1861.6 | 1154.3 | 1394.4 | 2238.2 |
| *QrSUVH3* | OCV4_rep_c21535 | ND | ND | ND | ND | ND | ND |
| *QrSUVH9* | Loc_7424_Tr_3/4_Conf_0.545_Len_5241 | 75.9 | 37.4 | 88.6 | 38.1 | 20.8 | 101.8 |
| *QrSUVR4-like* | OCV4_c20491 | 6.7 | 9.2 | 15.2 | 32.9 | 10.4 | 4.1 |
| *QrSUVR4* | Loc_53227_Tr_1/1_Conf_1.000_Len_2710 | 50.0 | 45.8 | 0.8 | 22.6 | 6.5 | 4.1 |
| *QrSUVR2* | OCV4_rep_c16379 | ND | ND | ND | ND | ND | ND |
| *QrSUVR5* | Loc_7632_Tr_2/3_Conf_0.800_Len_7750 | 127.8 | 164.8 | 161.2 | 123.5 | 89.8 | 334.0 |
| *QrSET40* | Loc_19922_Tr_2/5_Conf_0.636_Len_1707 | 336.4 | 374.5 | 351.9 | 153.3 | 149.7 | 291.2 |
| *QrSET10* | Loc_11855_Tr_5/6_Conf_0.611_Len_2331 | 120.1 | 89.9 | 122.4 | 71.0 | 80.7 | 97.8 |
| *QrSET41* | OCV4_c22093 | 321.0 | 461.8 | 378.9 | 452.7 | 416.6 | 356.4 |
| *QrASHR2* | OCV4_rep_c11271 | 556.5 | 1317.3 | 582.3 | 935.1 | 565.1 | 635.4 |
| *QrASHR1* | OCV4_rep_c3041 | ND | ND | ND | ND | ND | ND |
| *QrASHR1* | OCV4_rep_c943/Loc_12215_Tr_7/9_Conf_0.583_Len_3513 | 41.3 | 20.0 | 34.6 | 26.7 | 20.8 | 42.8 |
| *QrATXR4* | Loc_30882_Tr_3/3_Conf_0.714_Len_832 | 0.0 | 2.5 | 0.8 | 13.4 | 6.5 | 0.0 |
| *QrATXR1* | Loc_33752_Tr_1/1_Conf_1.000_Len_2058 | 2.9 | 126.5 | 1.7 | 85.4 | 61.2 | 51.9 |
| **Histone demethyltransferases** | | | | | | | |
| *QrJMJ24* | OCV4_c26318 | 674.7 | 501.8 | 426.2 | 310.7 | 665.3 | 506.1 |
| *QrJMJ25-like1* | OCV4_c27580 | 68.2 | 167.3 | 54.9 | 204.7 | 112.0 | 118.1 |
| *QrKDM3D-like2* | OCV4_rep_c9279 | ND | ND | ND | ND | ND | ND |
| *QrKDM3D-like1* | OCV4_rep_c15058 | 483.5 | 1126.7 | 458.2 | 412.5 | 266.9 | 290.2 |
| *QrKDM3B-like* | Loc_11265_Tr_3/5_Conf_0.692_Len_5052 | 19.2 | 67.4 | 19.4 | 8.2 | 14.3 | 22.4 |
| *QrJMJD6B* | OCV4_rep_c21199 | ND | ND | ND | ND | ND | ND |
| *QrJMJD6A* | OCV4_rep_c28281 | ND | ND | ND | ND | ND | ND |
| *QrJMJ32* | Loc_19662_Tr_4/5_Conf_0.500_Len_1503 | 20.2 | 16.6 | 25.3 | 21.6 | 10.4 | 23.4 |
| *QrJMJ32-like* | OCV4_rep_c1929 | ND | ND | ND | ND | ND | ND |
| *QrJMJ20* | OCV4_rep_c11681 | 19.2 | 13.3 | 23.6 | 26.7 | 43.0 | 15.3 |
| *QrJMJ16* | Loc_970_Tr_1/3_Conf_0.727_Len_4447 | 167.2 | 150.6 | 344.3 | 309.7 | 157.5 | 263.7 |
| *QrJMJ19-like* | OCV4_rep_c18633 | ND | ND | ND | ND | ND | ND |
| *QrKDM5* | Loc_1909_Tr_59/61_Conf_0.057_Len_4367 | 36.5 | 18.3 | 31.2 | 12.3 | 19.5 | 36.7 |
| *QrJMJ18* | Loc_9658_Tr_3/4_Conf_0.700_Len_4718 | 11.5 | 28.3 | 18.6 | 19.5 | 24.7 | 35.6 |
| *QrREF6* | OCV4_c5323 | 1482.1 | 3231.3 | 2221.1 | 1945.4 | 1809.7 | 1652.7 |
| *QrJMJ706* | Loc_2899_Tr_3/5_Conf_0.417_Len_2953 | 79.8 | 163.9 | 60.8 | 451.6 | 287.7 | 36.7 |
| *QrELF6* | OCV4_rep_c3275 | ND | ND | ND | ND | ND | ND |
| *QrLDL1* | OCV4_rep_c28326 | ND | ND | ND | ND | ND | ND |
| *QrFLD* | OCV4_rep_c42703 | ND | ND | ND | ND | ND | ND |
| *QrLDL2* | OCV4_c28695 | 706.4 | 877.9 | 418.6 | 445.5 | 497.4 | 478.6 |
| *QrLDL3* | Loc_2373_Tr_8/10_Conf_0.706_Len_11006 | 428.7 | 561.7 | 449.8 | 699.6 | 841.1 | 717.9 |
| **Histone deacetyltransferases** | | | | | | | |
| ***Gene name*** | **Gene ID** | **EcoDB** | **swB** | **XY** | **RO** | **LE** | **CA** |
| *QrHDA19-like1* | OCV4_rep_c21849 | ND | ND | ND | ND | ND | ND |
| *QrHDA19* | OCV3_primec7262/OCV4_rep_c13389 | 44.2 | 66.6 | 36.3 | 56.6 | 341.1 | 28.5 |
| *QrHDA6* | OCV4_rep_c26599 | ND | ND | ND | ND | ND | ND |
| *QrHDA9-like* | OCV3_prime_rep_c68588 | 8.7 | 5.8 | 7.6 | 50.4 | 26.0 | 6.1 |
| *QrHDA9* | OCV4_rep_c14185 | 39.4 | 63.2 | 31.2 | 37.0 | 39.1 | 11.2 |
| *QrHDA5* | Loc_6160_Tr_2_9_Conf_0.625_Len_2930 | 20.2 | 24.1 | 40.5 | 25.7 | 31.2 | 24.4 |
| *QrHDA15* | OCV4_rep_c12421 | 136.5 | 169.8 | 186.5 | 370.4 | 158.8 | 308.5 |
| *QrHDA14* | OCV3_prime_c9769 | 14.4 | 56.6 | 31.2 | 56.6 | 354.1 | 186.3 |
| *QrHDA8* | Loc_15145_Tr_8_9_Conf_0.280_Len_1833 | 275.9 | 428.6 | 419.4 | 254.1 | 742.1 | 523.4 |
| *QrHDA2* | OCV4_rep_c15012 | 1747.4 | 1342.3 | 942.6 | 679.0 | 854.1 | 421.6 |
| *QrHDT1-like2* | Loc_1428_Tr_2_6_Conf_0.650_Len_1466 | 369.1 | 422.7 | 449.8 | 256.2 | 225.2 | 353.3 |
| *QrHDT1-like1* | Loc_8805_Tr_2_4_Conf_0.833_Len_1959 | 2822.9 | 3538.3 | 2427.0 | 3209.7 | 1852.7 | 3649.6 |
| *QrSRT1-like1* | Loc_9090_Tr_3_7_Conf_0.611_Len_2531 | 0.0 | 0.0 | 0.0 | 1.0 | 0.0 | 0.0 |
| *QrSRT1-like2* | Loc_26807_Tr_3_3_Conf_0.778_Len_1873 | 20.2 | 11.7 | 15.2 | 159.5 | 53.4 | 76.4 |
| *QrSRT2* | OCV3_prime_c3720 | 300.8 | 327.9 | 318.1 | 163.6 | 962.2 | 251.5 |

| **Table S3: Normalized read counts of Quercus suber homologs of DNA and histone epigenetic modifiers as a result of the genome mapping of previously 454 sequenced libraries.** The tissues analyzed include: acorns in 3 development stages (S2, S3/4, S5), in embryos (E), in good and bad quality cork (C.Q), in first (1F and 1M) and last stages (2F and 2M) of female (F) and male flower (M) development, in roots (R.) with medium (MD), severe (SD) or without drought stress (Ww), in samples with red and open buds (ROp) and in samples with dormant and swollen buds (D/Sw). | | | | | | | | | | | | | | | | |
| --- | --- | --- | --- | --- | --- | --- | --- | --- | --- | --- | --- | --- | --- | --- | --- | --- |
|  |  | **Buds** | | **Male Flowers** | | **Female Flowers** | | **Acorns** | | | **Embryos** | **Cork Quality** | | **Roots** | | |
| **Gene name** | **Locus tag** | **ROp** | **DSw** | **1M** | **2M** | **1F** | **2F** | **S2** | **S3S4** | **S5** |  | **Good** | **Bad** | **WW** | **MD** | **SD** |
| **DNA Methyltransferases** | | | | | | | | | | | | | | | | |
| *QsMET1* | CFP56_38722 | 19.0 | 37.8 | 9.7 | 0.0 | 6.6 | 2.7 | 21.5 | 61.8 | 29.4 | 12.0 | 10.0 | 11.3 | 18.7 | 19.3 | 10.4 |
| *QsCMT1* | CFP56_24702 | 24.8 | 2.8 | 16.1 | 0.0 | 5.5 | 6.7 | 0.0 | 0.0 | 10.2 | 8.7 | 0.0 | 0.9 | 12.5 | 15.1 | 11.7 |
| *QsCMT2.1* | CFP56_06331 | 5.8 | 7.0 | 1.6 | 1.9 | 5.5 | 1.3 | 2.6 | 37.7 | 3.4 | 9.8 | 2.5 | 5.7 | 18.7 | 3.3 | 3.3 |
| *QsCMT3* | CFP56_35720 | 67.2 | 71.4 | 4.8 | 0.0 | 5.5 | 5.4 | 35.2 | 32.6 | 29.4 | 14.9 | 6.7 | 0.9 | 29.1 | 30.6 | 9.1 |
| *QsDRM2* | CFP56_35392 | 213.4 | 219.7 | 32.2 | 69.6 | 12.1 | 18.8 | 0.9 | 3.4 | 12.4 | 10.9 | 352.9 | 627.0 | 6.7 | 5.7 | 9.8 |
| *QsDRM3* | CFP56_66797 | 2.9 | 8.4 | 1.6 | 5.8 | 18.8 | 21.5 | 26.6 | 37.7 | 6.8 | 29.0 | 3.3 | 0.9 | 20.0 | 7.5 | 19.5 |
| *QsDNMT2* | CFP56_40985 | 16.1 | 30.8 | 14.5 | 15.5 | 15.4 | 2.7 | 8.6 | 5.1 | 7.9 | 23.6 | 6.7 | 11.3 | 19.6 | 10.4 | 5.2 |
| *QsCMT2.2* | CFP56_59471 | 0.0 | 0.0 | 1.6 | 5.8 | 1.1 | 0.0 | 4.3 | 20.6 | 0.0 | 3.3 | 0.8 | 2.8 | 3.3 | 1.9 | 0.7 |
| **DNA Demethylases** | | | | | | | | | | | | | | | | |
| *QsDME* | CFP56_06273 | 23.4 | 15.4 | 19.3 | 29.0 | 30.9 | 20.1 | 94.5 | 272.8 | 40.7 | 35.9 | 10.0 | 17.9 | 52.0 | 54.2 | 29.3 |
| *QsROS1* | CFP56_05353 | 16.1 | 15.4 | 66.0 | 5.8 | 39.7 | 26.9 | 209.6 | 332.8 | 369.8 | 117.4 | 51.6 | 30.2 | 166.8 | 199.2 | 168.1 |
| *QsDML2* | CFP56_35285 | 0.0 | 0.0 | 0.0 | 0.0 | 0.0 | 0.0 | 0.0 | 0.0 | 0.0 | 0.0 | 0.0 | 0.0 | 0.0 | 0.0 | 0.0 |
| **Histone Acetyltransferases** | | | | | | | | | | | | | | | | |
| *QsHAC1* | CFP56_16103 | 21.9 | 16.8 | 4.8 | 21.3 | 5.5 | 12.1 | 128.0 | 168.1 | 109.7 | 57.3 | 48.3 | 45.3 | 63.2 | 83.8 | 73.6 |
| *QsHAC1-like1* | CFP56_07357 | 0.0 | 0.0 | 4.8 | 0.0 | 0.0 | 0.0 | 0.9 | 0.0 | 11.3 | 13.4 | 0.0 | 0.0 | 0.8 | 2.8 | 2.0 |
| *QsHAC1-like2* | CFP56_07360 | 0.0 | 0.0 | 0.0 | 0.0 | 0.0 | 0.0 | 0.0 | 0.0 | 0.0 | 0.0 | 0.0 | 0.0 | 0.0 | 0.0 | 0.0 |
| *QsGCN5* | CFP56_29027 | 14.6 | 21.0 | 16.1 | 5.8 | 8.8 | 0.0 | 23.2 | 49.7 | 18.1 | 12.7 | 6.7 | 7.5 | 17.9 | 26.4 | 45.0 |
| *QsHAG2* | CFP56_43084 | 51.2 | 33.6 | 29.0 | 3.9 | 58.5 | 30.9 | 30.1 | 37.7 | 6.8 | 50.4 | 8.3 | 17.9 | 10.8 | 8.5 | 27.4 |
| *QsHAM1* | CFP56_58450 | 23.4 | 0.0 | 29.0 | 30.9 | 24.3 | 34.9 | 97.1 | 66.9 | 40.7 | 34.4 | 11.7 | 28.3 | 87.8 | 103.6 | 107.5 |
| *QsHAF1* | CFP56_26298 | 5.8 | 15.4 | 11.3 | 1.9 | 4.4 | 8.1 | 118.5 | 145.8 | 95.0 | 66.7 | 14.2 | 21.7 | 69.1 | 46.6 | 104.9 |
| *QsELP3* | CFP56_52694 | 52.6 | 35.0 | 32.2 | 38.6 | 24.3 | 24.2 | 25.8 | 10.3 | 57.7 | 36.6 | 25.0 | 49.0 | 34.1 | 28.7 | 30.0 |
| **Histone Methyltransferases** | | | | | | | | | | | | | | | | |
| *QsSET40* | CFP56_02401 | 0.0 | 0.0 | 12.9 | 25.1 | 3.3 | 6.7 | 4.3 | 0.0 | 20.4 | 15.6 | 5.0 | 2.8 | 3.3 | 3.8 | 2.0 |
| *QsASHR2* | CFP56_23528 | 4.4 | 14.0 | 4.8 | 19.3 | 11.0 | 6.7 | 2.6 | 5.1 | 2.3 | 4.0 | 0.0 | 0.0 | 5.4 | 1.9 | 2.0 |
| *QsASHR2-like* | CFP56_23526 | 4.4 | 7.0 | 24.1 | 36.7 | 28.7 | 13.4 | 6.0 | 8.6 | 9.0 | 9.8 | 1.7 | 0.0 | 3.3 | 1.4 | 6.5 |
| *QsSUVR5* | CFP56_22604 | 20.5 | 18.2 | 6.4 | 5.8 | 13.2 | 13.4 | 45.5 | 63.5 | 58.8 | 25.4 | 10.8 | 6.6 | 41.2 | 44.7 | 100.3 |
| *QsCLF* | CFP56_48440 | 13.2 | 11.2 | 19.3 | 13.5 | 8.8 | 1.3 | 39.5 | 30.9 | 22.6 | 17.4 | 10.0 | 3.8 | 59.9 | 33.0 | 61.2 |
| *QsATX3* | CFP56_04889 | 10.2 | 9.8 | 0.0 | 15.5 | 24.3 | 13.4 | 39.5 | 128.7 | 26.0 | 13.8 | 6.7 | 6.6 | 53.2 | 54.2 | 34.5 |
| *QsSUVH6* | CFP56_40587 | 0.0 | 2.8 | 3.2 | 13.5 | 2.2 | 2.7 | 7.7 | 15.4 | 18.1 | 13.4 | 4.2 | 4.7 | 7.9 | 8.5 | 2.0 |
| *QsSUVR2* | CFP56_56207 | 10.2 | 35.0 | 16.1 | 11.6 | 22.1 | 16.1 | 55.0 | 42.9 | 65.6 | 48.2 | 24.1 | 6.6 | 29.5 | 41.4 | 24.1 |
| *QsATX2* | CFP56_26820 | 17.5 | 9.8 | 6.4 | 3.9 | 5.5 | 1.3 | 28.3 | 85.8 | 30.5 | 27.5 | 14.2 | 3.8 | 12.5 | 16.0 | 55.4 |
| *ATXR2* | CFP56_29061 | 0.0 | 12.6 | 0.0 | 0.0 | 1.1 | 0.0 | 8.6 | 72.1 | 15.8 | 3.6 | 1.7 | 0.0 | 20.4 | 17.4 | 3.3 |
| *QsSUVR3* | CFP56_29051 | 0.0 | 1.4 | 3.2 | 11.6 | 1.1 | 4.0 | 0.0 | 17.2 | 2.3 | 2.2 | 3.3 | 0.9 | 2.9 | 0.0 | 2.6 |
| *QsSUVH9* | CFP56_44980 | 5.8 | 15.4 | 6.4 | 13.5 | 12.1 | 5.4 | 16.3 | 30.9 | 18.1 | 12.3 | 1.7 | 4.7 | 14.6 | 17.4 | 13.0 |
| *QsASHH2* | CFP56_13855 | 23.4 | 8.4 | 11.3 | 7.7 | 14.3 | 14.8 | 132.3 | 101.2 | 98.4 | 44.6 | 13.3 | 4.7 | 69.9 | 51.8 | 59.3 |
| *QsSUVR4* | CFP56_60043 | 0.0 | 0.0 | 0.0 | 0.0 | 0.0 | 0.0 | 0.0 | 0.0 | 10.2 | 25.7 | 0.0 | 0.0 | 0.8 | 0.0 | 0.0 |
| *QsASHH3* | CFP56_26485 | 20.5 | 2.8 | 16.1 | 17.4 | 28.7 | 12.1 | 4.3 | 30.9 | 20.4 | 43.5 | 25.0 | 14.1 | 25.4 | 18.8 | 20.2 |
| *QsATXR1* | CFP56_15756 | 0.0 | 0.0 | 0.0 | 0.0 | 0.0 | 0.0 | 0.0 | 0.0 | 0.0 | 0.0 | 0.0 | 0.0 | 0.0 | 0.0 | 0.0 |
| *QsSUVH4-like* | CFP56_01500 | 21.9 | 0.0 | 0.0 | 5.8 | 1.1 | 0.0 | 1.7 | 3.4 | 26.0 | 6.2 | 0.8 | 0.0 | 0.8 | 10.4 | 7.8 |
| *QsATXR6* | CFP56_57230 | 13.2 | 16.8 | 12.9 | 0.0 | 0.0 | 2.7 | 0.0 | 0.0 | 0.0 | 2.9 | 0.0 | 0.0 | 0.8 | 2.8 | 4.6 |
| *QsASHR3* | CFP56_53193 | 10.2 | 14.0 | 6.4 | 0.0 | 27.6 | 6.7 | 5.2 | 10.3 | 13.6 | 16.7 | 5.0 | 3.8 | 3.7 | 4.7 | 18.9 |
| *QsATXR7* | CFP56_14674 | 2.9 | 1.4 | 8.0 | 0.0 | 9.9 | 6.7 | 17.2 | 20.6 | 9.0 | 18.5 | 20.8 | 4.7 | 9.6 | 12.2 | 32.6 |
| *QsATX5* | CFP56_25529 | 16.1 | 18.2 | 0.0 | 11.6 | 5.5 | 2.7 | 50.7 | 46.3 | 53.2 | 15.9 | 5.0 | 8.5 | 36.6 | 72.5 | 50.8 |
| *QsSUVH1* | CFP56_34169 | 13.2 | 9.8 | 16.1 | 9.7 | 4.4 | 12.1 | 34.4 | 25.7 | 39.6 | 16.7 | 12.5 | 16.0 | 26.2 | 24.5 | 20.2 |
| *QsSUVH3* | CFP56_28232 | 19.0 | 2.8 | 9.7 | 3.9 | 16.5 | 4.0 | 46.4 | 29.2 | 45.2 | 18.5 | 12.5 | 15.1 | 23.7 | 26.8 | 20.2 |
| *QsASHH1* | CFP56_54040 | 5.8 | 0.0 | 8.0 | 0.0 | 16.5 | 6.7 | 10.3 | 8.6 | 29.4 | 26.8 | 0.8 | 1.9 | 13.3 | 12.2 | 5.9 |
| *QsSUVH4* | CFP56_37589 | 35.1 | 4.2 | 3.2 | 29.0 | 32.0 | 67.1 | 52.4 | 46.3 | 96.1 | 29.4 | 15.0 | 25.5 | 48.7 | 89.0 | 31.9 |
| *QSASHR1* | CFP56_31286 | 0.0 | 9.8 | 24.1 | 17.4 | 7.7 | 10.7 | 30.9 | 0.0 | 1.1 | 18.8 | 4.2 | 8.5 | 17.5 | 26.4 | 11.1 |
| *QsATXR5* | CFP56_53906 | 1.5 | 0.0 | 9.7 | 15.5 | 8.8 | 2.7 | 12.0 | 5.1 | 4.5 | 4.3 | 0.0 | 0.0 | 0.8 | 0.9 | 0.0 |
| *QsSWN* | CFP56_32659 | 0.0 | 0.0 | 1.6 | 19.3 | 2.2 | 5.4 | 20.6 | 8.6 | 24.9 | 20.3 | 6.7 | 6.6 | 30.4 | 22.1 | 22.2 |
| *QsSET41* | CFP56_40485 | 0.0 | 0.0 | 1.6 | 0.0 | 2.2 | 4.0 | 16.3 | 6.9 | 1.1 | 5.4 | 5.0 | 0.0 | 2.9 | 13.7 | 10.4 |
| *QsSUVH5b* | CFP56_65722 | 0.0 | 0.0 | 0.0 | 0.0 | 0.0 | 0.0 | 0.0 | 0.0 | 0.0 | 0.0 | 0.0 | 0.0 | 0.0 | 0.0 | 0.0 |
| *QsSUVH5a* | CFP56_65724 | 0.0 | 0.0 | 0.0 | 0.0 | 0.0 | 0.0 | 0.0 | 0.0 | 0.0 | 0.0 | 0.0 | 0.0 | 0.0 | 0.0 | 0.0 |
| *QsSUVR4-likeb* | CFP56_76543 | 0.0 | 0.0 | 0.0 | 0.0 | 0.0 | 0.0 | 0.0 | 0.0 | 0.0 | 0.0 | 0.0 | 0.0 | 0.0 | 0.0 | 0.0 |
| *QsSUVR4-likea* | CFP56_36057 | 0.0 | 0.0 | 0.0 | 0.0 | 0.0 | 0.0 | 0.0 | 0.0 | 0.0 | 0.0 | 0.0 | 0.0 | 0.0 | 0.0 | 0.0 |
| *QsSET10* | CFP56_45284 | 21.9 | 0.0 | 11.3 | 9.7 | 2.2 | 0.0 | 3.4 | 13.7 | 15.8 | 10.1 | 10.0 | 2.8 | 3.3 | 14.6 | 14.3 |
| *QSATXR3* | CFP56_27073 | 1.5 | 2.8 | 0.0 | 0.0 | 1.1 | 4.0 | 13.7 | 1.7 | 6.8 | 2.2 | 0.0 | 0.0 | 5.0 | 4.7 | 10.4 |
| **Histone Demethylases** | | | | | | | | | | | | | | | | |
| *QsJMJ18* | CFP56_59257 | 4.4 | 8.4 | 8.0 | 25.1 | 2.2 | 4.0 | 27.5 | 6.9 | 45.2 | 44.6 | 10.8 | 12.3 | 9.2 | 7.1 | 13.7 |
| *QsKDM3B-like* | CFP56_10640 | 8.8 | 2.8 | 3.2 | 9.7 | 5.5 | 0.0 | 25.8 | 20.6 | 23.7 | 20.3 | 10.8 | 17.9 | 12.1 | 16.0 | 28.0 |
| *QsJMJ25-like1* | CFP56_48908 | 10.2 | 7.0 | 20.9 | 42.5 | 13.2 | 18.8 | 42.1 | 41.2 | 49.8 | 46.4 | 15.8 | 0.9 | 35.4 | 26.4 | 26.7 |
| *QsJMJ25-like2* | CFP56_47938 | 0.0 | 0.0 | 0.0 | 0.0 | 0.0 | 0.0 | 0.0 | 0.0 | 0.0 | 0.0 | 0.0 | 0.0 | 0.0 | 0.0 | 0.0 |
| *QsJMJ25-like3* | CFP56_18193 | 5.8 | 12.6 | 9.7 | 0.0 | 7.7 | 2.7 | 42.1 | 27.4 | 40.7 | 15.9 | 10.8 | 13.2 | 18.7 | 28.3 | 11.1 |
| *QsJMJ25-like4* | CFP56_18195 | 4.4 | 0.0 | 1.6 | 0.0 | 1.1 | 1.3 | 23.2 | 8.6 | 26.0 | 16.3 | 15.0 | 1.9 | 1.7 | 3.8 | 0.7 |
| *QsJMJ24* | CFP56_37500 | 8.8 | 4.2 | 12.9 | 1.9 | 11.0 | 2.7 | 55.0 | 39.5 | 83.7 | 47.5 | 30.0 | 2.8 | 28.7 | 35.3 | 81.4 |
| *QsKDM3D-like* | CFP56_34178 | 59.9 | 72.8 | 3.2 | 5.8 | 22.1 | 8.1 | 104.8 | 137.2 | 229.6 | 64.9 | 28.3 | 5.7 | 59.5 | 56.0 | 33.9 |
| *QsJMJD6Aa* | CFP56_37891 | 17.5 | 4.2 | 4.8 | 11.6 | 3.3 | 5.4 | 0.0 | 0.0 | 1.1 | 6.5 | 5.0 | 5.7 | 2.9 | 5.7 | 9.1 |
| *QsJMJD6Ab* | CFP56_50389 | 0.0 | 0.0 | 0.0 | 0.0 | 0.0 | 0.0 | 0.0 | 0.0 | 0.0 | 0.0 | 0.0 | 0.0 | 0.0 | 0.0 | 0.0 |
| *QsJMJ16* | CFP56_76189 | 0.0 | 0.0 | 1.6 | 0.0 | 1.1 | 1.3 | 16.3 | 22.3 | 37.3 | 3.6 | 0.0 | 10.4 | 9.2 | 37.2 | 13.0 |
| *QsJMJ19-like* | CFP56_17215 | 2.9 | 5.6 | 3.2 | 3.9 | 2.2 | 4.0 | 28.3 | 13.7 | 10.2 | 21.7 | 7.5 | 2.8 | 25.4 | 24.5 | 15.6 |
| *QsJMJ706* | CFP56_36126 | 0.0 | 0.0 | 0.0 | 0.0 | 0.0 | 0.0 | 0.0 | 0.0 | 0.0 | 0.0 | 0.0 | 0.0 | 0.0 | 0.0 | 0.0 |
| *QsJMJ706-like* | CFP56_37564 | 80.4 | 454.9 | 130.3 | 394.1 | 200.8 | 155.8 | 88.5 | 22.3 | 139.1 | 110.9 | 40.8 | 52.8 | 281.2 | 331.1 | 562.9 |
| *QsLDL3* | CFP56_08883 | 5.8 | 0.0 | 4.8 | 5.8 | 2.2 | 5.4 | 36.1 | 60.0 | 35.1 | 13.0 | 2.5 | 11.3 | 18.7 | 23.1 | 23.5 |
| *QsELF6* | CFP56_41197 | 7.3 | 0.0 | 1.6 | 0.0 | 1.1 | 1.3 | 28.3 | 13.7 | 15.8 | 8.0 | 0.8 | 7.5 | 9.2 | 7.1 | 8.5 |
| *QsJMJD6B* | CFP56_61743 | 19.0 | 14.0 | 12.9 | 9.7 | 8.8 | 37.6 | 42.9 | 75.5 | 61.1 | 41.0 | 20.8 | 6.6 | 27.5 | 25.9 | 37.1 |
| *QsJMJ32-like* | CFP56_26267 | 0.0 | 0.0 | 0.0 | 0.0 | 0.0 | 0.0 | 0.0 | 0.0 | 0.0 | 0.0 | 0.0 | 0.0 | 0.0 | 0.0 | 0.0 |
| *QsJMJ32* | CFP56_26269 | 0.0 | 0.0 | 0.0 | 0.0 | 0.0 | 0.0 | 0.0 | 0.0 | 0.0 | 0.0 | 0.0 | 0.0 | 0.0 | 0.0 | 0.0 |
| *QsREF6* | CFP56_69927 | 119.8 | 89.6 | 67.6 | 69.6 | 60.7 | 67.1 | 128.0 | 51.5 | 151.5 | 48.6 | 35.8 | 49.0 | 104.8 | 130.5 | 95.1 |
| *QsLDL2* | CFP56_19117 | 1.5 | 0.0 | 0.0 | 0.0 | 1.1 | 0.0 | 1.7 | 5.1 | 3.4 | 4.7 | 0.8 | 1.9 | 5.8 | 15.5 | 2.0 |
| *QsLDL1* | CFP56_55051 | 4.4 | 14.0 | 3.2 | 3.9 | 3.3 | 4.0 | 4.3 | 8.6 | 18.1 | 2.9 | 2.5 | 1.9 | 3.7 | 3.3 | 3.9 |
| *QsKDM5* | CFP56_46886 | 0.0 | 7.0 | 0.0 | 0.0 | 7.7 | 6.7 | 82.5 | 30.9 | 58.8 | 15.2 | 3.3 | 1.9 | 13.7 | 16.0 | 18.2 |
| *QsJMJ30* | CFP56_23213 | 1.5 | 2.8 | 1.6 | 0.0 | 5.5 | 0.0 | 8.6 | 15.4 | 5.7 | 4.7 | 15.0 | 5.7 | 4.2 | 8.0 | 9.8 |
| *QsFLD* | CFP56_03400 | 0.0 | 5.6 | 4.8 | 3.9 | 0.0 | 0.0 | 9.4 | 22.3 | 28.3 | 4.0 | 3.3 | 1.9 | 12.5 | 15.5 | 9.8 |
| *QsJMJ20* | CFP56_45215 | 0.0 | 1.4 | 3.2 | 7.7 | 12.1 | 1.3 | 0.0 | 37.7 | 6.8 | 12.7 | 0.0 | 0.9 | 18.3 | 5.7 | 18.2 |
| **Histone Deacetylases** | | | | | | | | | | | | | | | | |
| *QsHDA2* | CFP56_55621 | 0.0 | 5.6 | 22.5 | 59.9 | 24.3 | 38.9 | 98.8 | 49.7 | 15.8 | 42.4 | 40.0 | 10.4 | 8.7 | 27.8 | 48.2 |
| *QsHDA5* | CFP56_01537 | 20.5 | 14.0 | 19.3 | 40.6 | 18.8 | 29.5 | 55.8 | 36.0 | 63.3 | 26.1 | 19.1 | 14.1 | 30.0 | 35.3 | 5.9 |
| *QsHDA6* | CFP56_50628 | 2.9 | 57.4 | 17.7 | 36.7 | 36.4 | 18.8 | 39.5 | 30.9 | 54.3 | 44.6 | 32.5 | 35.8 | 17.9 | 21.7 | 31.9 |
| *QsHDA19-like2* | CFP56_14676 | 39.5 | 46.2 | 49.9 | 42.5 | 67.3 | 94.0 | 67.0 | 41.2 | 36.2 | 54.0 | 40.0 | 39.6 | 38.3 | 53.2 | 62.5 |
| *QsHDA19* | CFP56_26357 | 43.8 | 54.6 | 29.0 | 27.0 | 60.7 | 41.6 | 35.2 | 34.3 | 44.1 | 31.9 | 17.5 | 30.2 | 40.4 | 26.4 | 18.2 |
| *QsHDA9* | CFP56_22884 | 20.5 | 46.2 | 48.3 | 65.7 | 51.8 | 53.7 | 55.8 | 30.9 | 74.6 | 79.7 | 13.3 | 111.3 | 113.1 | 100.3 | 61.9 |
| *QsHDA19-like1* | CFP56_75988 | 0.0 | 0.0 | 0.0 | 0.0 | 0.0 | 0.0 | 0.0 | 0.0 | 0.0 | 0.0 | 0.0 | 0.0 | 0.0 | 0.0 | 0.0 |
| *QsHDA8* | CFP56_03621 | 16.1 | 9.8 | 12.9 | 3.9 | 5.5 | 6.7 | 4.3 | 1.7 | 13.6 | 18.5 | 1.7 | 0.0 | 2.1 | 0.9 | 5.9 |
| *QsHDA14-like1* | CFP56_64318 | 5.8 | 16.8 | 8.0 | 5.8 | 3.3 | 5.4 | 0.0 | 5.1 | 3.4 | 5.1 | 2.5 | 3.8 | 2.5 | 8.0 | 0.0 |
| *QsHDA14-like2* | CFP56_32080 | 5.8 | 23.8 | 6.4 | 13.5 | 6.6 | 0.0 | 0.0 | 3.4 | 11.3 | 14.5 | 6.7 | 2.8 | 0.0 | 0.0 | 1.3 |
| *QsHDA14-like3* | CFP56_65290 | 2.9 | 2.8 | 4.8 | 1.9 | 1.1 | 0.0 | 0.0 | 0.0 | 1.1 | 5.8 | 1.7 | 0.0 | 0.0 | 0.5 | 0.0 |
| *QsHDA15* | CFP56_10834 | 67.2 | 89.6 | 54.7 | 23.2 | 75.0 | 83.2 | 137.4 | 87.5 | 80.3 | 105.1 | 124.0 | 61.3 | 74.9 | 105.0 | 89.3 |
| *QsHDT1-like1* | CFP56_42573 | 371.2 | 340.1 | 194.7 | 156.5 | 142.3 | 111.4 | 78.2 | 126.9 | 47.5 | 58.0 | 100.7 | 216.9 | 131.9 | 119.6 | 138.1 |
| *QsHDT1-like3* | CFP56_05349 | 49.7 | 63.0 | 74.0 | 40.6 | 46.3 | 102.0 | 48.1 | 48.0 | 64.5 | 88.1 | 70.8 | 138.6 | 43.3 | 16.0 | 39.7 |
| *QsSRT1-like3* | CFP56_78126 | 2.9 | 4.2 | 3.2 | 25.1 | 3.3 | 4.0 | 8.6 | 6.9 | 5.7 | 21.7 | 0.8 | 5.7 | 10.0 | 2.8 | 3.3 |
| *QsSRT1-like5* | CFP56_38596 | 5.8 | 0.0 | 0.0 | 3.9 | 0.0 | 4.0 | 0.9 | 1.7 | 6.8 | 6.9 | 1.7 | 0.0 | 1.7 | 0.0 | 0.0 |
| *QsSRT1-like4* | CFP56_42400 | 4.4 | 0.0 | 0.0 | 1.9 | 0.0 | 6.7 | 1.7 | 6.9 | 0.0 | 12.7 | 0.0 | 2.8 | 0.0 | 0.0 | 2.6 |
| *QsSRT1-like2* | - | ND | ND | ND | ND | ND | ND | ND | ND | ND | ND | ND | ND | ND | ND | ND |
| *QsSRT1-like1* | - | ND | ND | ND | ND | ND | ND | ND | ND | ND | ND | ND | ND | ND | ND | ND |
| *QsSRT2* | CFP56_42602 | 4.4 | 1.4 | 0.0 | 27.0 | 2.2 | 8.1 | 6.0 | 12.0 | 5.7 | 10.9 | 2.5 | 14.1 | 2.1 | 1.9 | 1.3 |
| *QsHDT1-like1* | CFP56_51836 | ND | ND | ND | ND | ND | ND | ND | ND | ND | ND | ND | ND | ND | ND | ND |
| *QsHDA9-like* | - | ND | ND | ND | ND | ND | ND | ND | ND | ND | ND | ND | ND | ND | ND | ND |
